# Estimating the changing risks of low crop yield using non-stationary generalized Pareto distributions

**DOI:** 10.1101/2025.05.10.653234

**Authors:** Akio Onogi

**Author notes:** **Corresponding author:** Akio Onogi, Postal address: 1-5, Yokotani, Oe-cho, Seta, Otsu, Shiga, 520-2194, Japan.

## Abstract

Estimating the changing risk of low crop yield in a changing climate is an important task in various fields of agricultural research. According to the extreme value theory, the probability of extreme events can be approximated using generalized Pareto distributions. In this study, non-stationary generalized Pareto distributions were used to estimate the changing risk of low crop yield. The proposed methods were applied to global yield data for maize, wheat, rice, and soybean collected from 1961 to 2022, as well as local yield data for wheat, rice, and soybean in Japan, from a start date of either 1948 or 1958 and running to 2020. The results illustrated exacerbated trends of low-yield risk in maize crops in Africa; maize, wheat, and rice in Americas; maize and wheat in Western, Central, and Southern Asia; maize and wheat in Europe; and soybean in Japan. Only wheat in Japan showed trends of mitigating the risk of low yields. The proposed models were also validated through simulations. The results showed that the models can generally estimate the changing risks accurately, and the precision depends on the size of the data set. Although there is still room for improvement in the models, the present study demonstrates that it is possible to estimate changes in the risk of low yield using a data-driven approach based on extreme value theory without assumptions about climate and crop physiology.

## 1. Introduction

Climate is a crucial factor controlling crop yield variability. Many studies have shown that climate variability, such as variability in temperature and rainfall, affects the crop yield. For example, Ray *et al* (2015) reported that approximately 60% or more of yield variability in main crops in major growing regions can be explained by climate variability. Zampieri *et al* (2017) reported that 42 % of the interannual variability in wheat yield is explained by the variability of a climate index for drought and heat stress, and Vogel *et al* (2019) reported that 18 % to 43 % of the variance in yield anomalies of major crops can be explained by climate extremes. Together, these findings highlight how global climate change can affect the risk of crop yield loss or anomalies.

Estimating the risk of yield loss and anomalies is a focus area in agricultural research and the agricultural industry and related professions. For farmers, risk information helps changes in management (planting time, crop variety, or management practices) to mitigate yield losses (Howden *et al* 2007, Deressa *et al* 2009). Companies in food supply chains can use risk estimation to manage their supply, reduce financial losses, and make stable contracts (Jüttner *et al* 2003, Tang 2006). Yield risk estimation is a useful tool for policymakers working to mitigate the risk of food insecurity (Funk *et al* 2008). Additionally, insurance companies need reliable risk assessments to design agricultural insurance and protect farmers from significant financial damage.

Yield variability or yield loss/anomalies are often analyzed using crop growth models or their emulators (e.g., Sultan *et al* 2014, Liu *et al* 2021, Kamali *et al* 2022, Hsiao *et al* 2024). Future yield variability can then be predicted under climate conditions forecasted by global climate models (Faye *et al* 2018). However, the results often vary depending on the crop models used (Faye *et al* 2018), and the predicted yield is inevitably affected by the uncertainties introduced by both types of models: crop growth and global climate models (Müller *et al* 2021). Non-crop-growth model-based approaches have also been applied to yield variability analyses; for example, Rathore *et al* (2024) used copula functions to discover the relationship between crop yield variability in the United States and a drought stress index and then estimated the yield loss risk. Ceglar *et al* (2016) used partial least squares regression to relate the interannual variability of wheat and maize in France to meteorological information. Leng and Hall (2020) compared crop growth models, linear regression, and random forest to predict maize yield variability in the United States and reported the superiority of the random forest approach. Such approaches may have advantages over crop-growth-model-based approaches because they rely on fewer model assumptions.

If a yield loss or anomaly is considered an extreme event that occurs occasionally, the probability (risk) of yield loss can be estimated using a class of extreme value theory, which states that the upper tail of the unknown population distribution over a sufficiently large threshold can be approximated using a generalized Pareto (GP) distribution (Coles *et al* 2001). By reversing the direction of the yield distributions, GP distributions can be used to estimate crop yield losses below a given threshold. The GP distribution-based approaches have the following advantages. (1) Unlike crop growth models, these approaches do not rely on assumptions about crop physiology and do not require information about the climate and soil. This allows for data-driven estimation of yield loss risk, which is applicable when yield figures are available. (2) By approximating the tail of the population distribution, the risk of future extreme events can be predicted. To date, there are two published studies in which the GP distribution was applied to estimate the risk of low yield. Park *et al* (2019) applied GP distributions to approximate the lower tail of the annual yield records of maize and wheat in the United States and estimated the probability (risk) of low yield. Chou *et al* (2024) used GP distributions to approximate the lower tail of the annual yield distributions of rice, wheat, and maize in China, Korea, and Japan to estimate the risk of low yield. In both of these studies, the authors adopted stationary GP models, in which the tail distribution of crop yield was assumed to be constant across years. However, this assumption may be unrealistic if the yield variability is affected by global climate change.

In the present study, nonstationary GP models were developed to approximate the lower tails of crop yield distributions and estimate the low-yield risks that change across years. The proposed method was applied to yield data on staple crops (maize, wheat, rice, and soybeans) from global regions and Japanese prefectures. The risk of low yield for the five world regions and Japan were then estimated. Japanese yield data were added to the analyses to estimate the risk at finer scale than is the case with global yield data. The contributions of the present study are: (1) a proposed and validated methodology to apply non-stationary GP models for estimating low-crop yield risks, (2) evaluation of the non-stationary GP model-based approaches by comparing the results with previous studies and simulation analyses, and (3) a discussion of the limitations of the proposed method and future extensions. Although the proposed methodology is still at the proof-of-concept stage, the results demonstrate that extreme value theory-based approaches can complement other approaches, such as those based on crop growth models and machine learning, to estimate the risk of low crop yield.

## 2. Methods

### 2.1. Global yield data

Global yield data were downloaded from the FAOSTAT website (https://www.fao.org/faostat/en/#home) on June 12^th^, 2024. The data included yield (100 g/ha) of maize (item code 112), rice (113), soya beans (soybean, 141), and wheat (111) of 192 countries (Table S1). The period was generally 1961 to 2022, with some variations between countries. The fill rate of yield records was 0.595: the total number of yield records was 28,318 and the data consisted of 4 items (maize, rice, soybean, and wheat), 192 countries, and 62 years, resulting in 47,616 (4 × 192 × 62) combinations.

The 192 countries were grouped into five areas, and the risk of low crop yield was estimated for each area (Figure 1 and Table S1). This manipulation is required to obtain sufficient yield records for the approximation with GP distributions, given that the number of records of a country was only 62 at most, which was too few for the approximation. However, the yield records of adjacent countries can correlate and induce spatial dependency. This issue is discussed in “4.2. Spatial dependency between yield records” in the Discussion section. The grouping followed the M49 standard (Standard country or area codes for statistics use, revision 4, 1999); however, some modifications were made to arrange the number of countries of each area as much as close to each other. Because Oceania, according to the M49 standard, consists of only seven countries and so was grouped with Southeastern and Eastern Asia (referred to as ESeAsia-Oc). However, because Asia consists of 50 countries, Western, Central, and Southern Asia were separated from Southeastern and Eastern Asia and grouped together (WCS Asia). The resulting areas (number of member countries) were as follows: Africa (55), Americas (37), WCSAsia (31), ESeAsia-Oc (26), and Europe (43) (Table S1).

**Figure 1.**
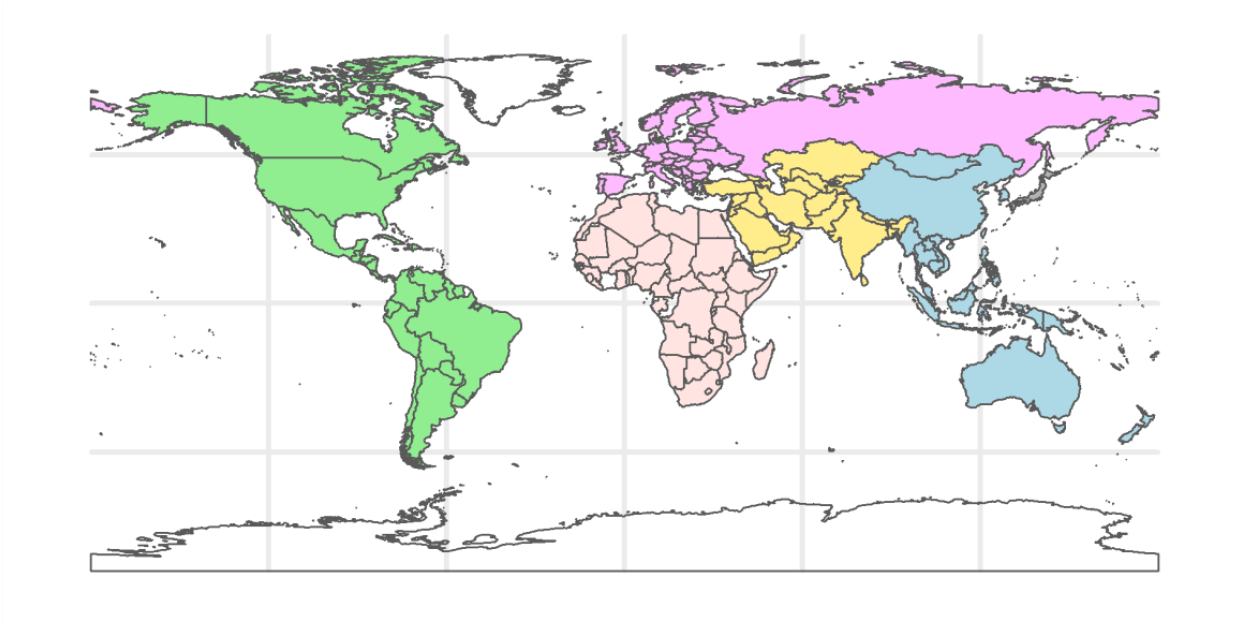
Areas where the risk of low crop yields was estimated. The global yield data consisted of data from five areas: Africa (red), Americas (green), Western, Central, and Southern Asia (WCSAsia, yellow), Eastern and Southern eastern Asia and Oceania (ESeAsia_Oc, blue), and Europe (purple). The local yield data consists of data from 47 prefectures in Japan (gray).

### 2.2. Local yield data

Yield data from Japan were downloaded from the Portal Site of Official Statistics of Japan, e-Stat (https://www.e-stat.go.jp/en) using “estatapi” R package (ver. 0.4.0) in April, 2024. The data included the yield (kg/10 a) of rice (paddy rice), soybeans, and wheat. Maize (corn) was not included because it is not a staple crop in Japan, and its yield records were much lower than those of other crops. The rice and wheat data included records from 1958 to 2020, whereas the soybean data included records from 1948 to 2020. Each crop dataset consisted of records from all the prefectures in Japan (*N* = 47) (Table S2). For each crop, the records in Okinawa Prefecture began in 1974; records before that year were missing. For rice and soybean, the records of the other 46 prefectures were available for the entire study period; in the case of wheat, 34 records were missing in six prefectures (Table S2). Altogether, data from 47 prefectures were used to estimate the risk of low yield of each of the crops in Japan.

### 2.3. Data processing

Both global and local yield data were processed before analyses in similar manners. First, yield records at each country (prefecture) were detrended by removing 5-year moving averages. Next, each country (prefecture) record was scaled such that the variances of each country (prefecture) were 1.0. The standard deviations (SDs) used for scaling are presented in Tables S1 for the global yield data and Table S2 for the local yield data. Because of scaling, Cuba (area: Americas) and French Guiana (Americas) were excluded from the analyses of soybean because the records had no variances: Cuba had two-year records (2021 and 2022) with the same yield (2,500 kg/ha) and French Guiana had only one-year record (1986, 2,300 kg/ha). The detrended and scaled records were then multiplying with −1, because the GP distributions are defined for positive real numbers. Stationary and non-stationary GP models were applied to these detrended, scaled, and sign-inverted records (hereafter referred to as transformed records). To be exact, the GP models were applied to values of the transformed records that exceeded thresholds.

### 2.4. GP distribution

The density function of GP distributions is

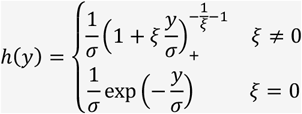

where *y* is the random variable, and *ξ* and σ are the shape and scale parameters (Coles *et al* 2001). The stationary GP models in the present study applied this GP distribution to values of the transformed records that exceeded certain thresholds. To estimate the parameters efficiently, it is essential to determine the range of the shape parameter (*ξ*), *ξ* > 0, *ξ* < 0, or *ξ* = 0, before estimation. These ranges indicate that the distribution of the variables of interest belongs to the domain of attraction of the Fréchet, Weibull, and Gumbel distributions (Coles *et al* 2001). The ranges of *ξ* were determined by creating the mean excess plots using “ismev” R package (ver. 1.42). As a result, because the crop-yield data was suggested to belong to the domain of attraction of the Weibull distribution (see the Result section), *ξ* was assumed to be negative. Thus, a positive parameter *nξ* which is defined as *nξ* = −*ξ* was introduced.

### 2.5. Stationary generalized Pareto model

The stationary GP model applies the GP distribution to the values of the transformed records that exceed the threshold *T*_*h*_. The likelihood function is expressed as follows:

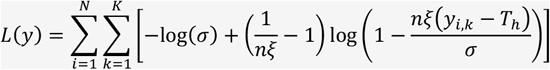

where *N* and *K* are the number of years and countries/prefectures, respectively, and *y*_*i,k*_ is the transformed record of country *k* in year *i*. Note that *N* differs between data types (global and local) and crops, and *K* differs between data types and areas. If *y*_*i,k*_ were missing, the corresponding likelihood was omitted. To evaluate the logarithm function, 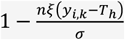 should be positive for all *y*_*i,k*_. Thus, the following constraints are imposed on each parameter:

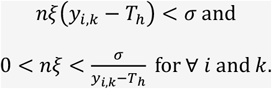

### 2.6. Non-stationary generalized Pareto model

The stationary GP model assumes that the distribution of values that exceed thresholds remains unchanged across years. In other words, low-yield risks were assumed to be constant across the years. A non-stationary GP model was developed to model changing risks. The model can be written as follows:

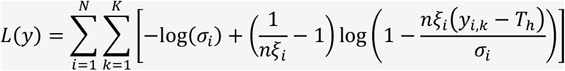

where

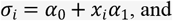

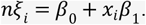

Herein, 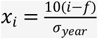 where *i* is the year, *f* denotes the year before the yield record started, and 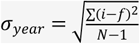. For example, if the yield data consists of 62-year records (*i* = 1961, …, 2022), *f* = 1960 and 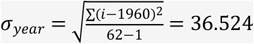. Multiplication with 10 in *x*_*i*_ was introduced to stabilize the computation using Markov chain Monte Carlo (MCMC) algorithms. In the non-stationary GP model, the shape (*nξ*_*i*_) and scale (*σ*_*i*_) parameters can be changed linearly across years (Coles *et al* 2001). The threshold *T*_*h*_ is assumed to be constant across years. This assumption stems from the assumption that even if the yield distributions change over the years, the values over the threshold *T*_*h*_ of each year can be approximated with the GP distributions (i.e., *T*_*h*_ is sufficiently large for each year). In this model, constraints were imposed on these intercepts (*α*_0_ and *β*_0_) and slopes (*α*_1_ and *β*_1_) as

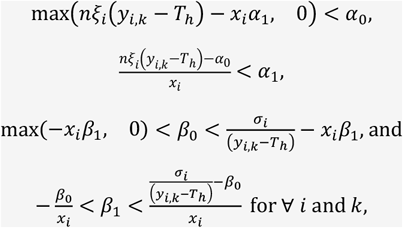

where max indicates the larger value between the two arguments. These constraints were derived from the requirements, 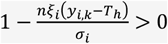, and *σ*_*i*_ > 0 and *nξ* > 0 at year *f*.

### 2.7. Parameter estimation

The parameters were estimated using Markov chain Monte Carlo (MCMC) algorithms. For the stationary GP models, the prior distributions of the parameters (*σ* and *nξ*) were

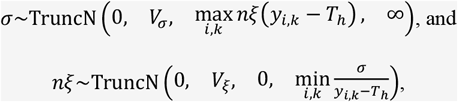

respectively. Herein, TruncN(*a, b, c, d*) indicate a normal distribution with mean *a* and variance *b* truncated between *c* and *d*. These truncated normal distributions are defined based on the constraints mentioned in the previous section. The variances (*V*_*σ*_ and *V*_*ξ*_) were fixed to a large value (10,000) to make the prior distributions flat.

The prior distributions of the non-stationary GP models (*α*_0_, *α*_1_, *β*_0_, and *β*_1_) were

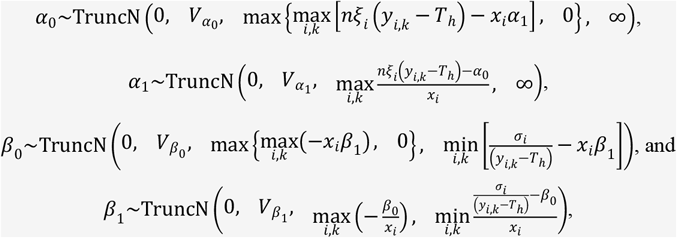

respectively. The variances (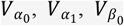, and 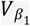) were fixed to a large value (10,000).

The parameters were updated using the random-walk algorithm. The proposed values of parameter *θ* were generated as

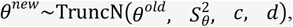

where *S*_*θ*_ was the standard deviation of the proposal distribution. The parameter *θ* was either of *σ* or *nξ* in the stationary GP models and either of *α*_0_, *α*_1_, *β*_0_, or *β*_1_ in the non-stationary GP models. The standard deviations *S*_*θ*_ was determined such that the acceptance rate of the new values largely ranged from 0.1 to 0.6. The range (*c* and *d*) was determined based on the constraints imposed on each parameter. The acceptance probability was defined using the likelihood (*L*(*y*)), the probability densities of the prior truncated normal distributions denoted by *Prior*(*θ*), and the probability densities of the proposal truncated normal distributions denoted by *Proposal*(*θ*) as

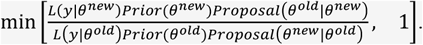

For the stationary GP models, the number of iterations, burn-in length, and sampling interval were 100,000, 80,000, and 10, respectively. The nonstationary GP models were 500,000, 400,000, and 50 for local yield data and 1,000,000, 800,000, and 100 for global yield data. Each condition yields

2,000 MCMC samples. The MCMC was repeated five times with different initial values, resulting in 10,000 samples. Low-yield risks were estimated using 10,000 samples. Convergence was verified using 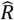 statistics (Gelman *et al* 2014). The MCMC computations were done using the house-developed C++ programs run under “Rcpp” R package (ver. 1.0.13-1) (Eddelbuettel and Balamuta 2018). These programs are available upon request.

### 2.8. Model comparison

Stationary and nonstationary GP models were compared using the widely applicable information criterion (WAIC, Watanabe 2010). WAIC was calculated from MCMC samples using “WAIC” function of “LaplacesDemon” R package (ver. 16.1.6).

### 2.9. *M*-year return level

The risk of low yield was estimated as *M*-year return levels. Herein, *M* indicates a period when the low yield worse than the level is expected to occur once. For example, when the 10-year return level is −2, yield of 2 SD or much lower than the average can occur once in 10 years. This SD is the SD of detrended yield which was used for scaling yield data (Tables S1 and S2), and the average is the average of the 5-year moving window. The *M*-year return level was reported as a continuous curve by taking *M* and standardized yield as *x* and *y* axes, respectively. As *M*, 20 values from 10, 20, …, 190, and 200 were chosen.

The *M*-year return level was estimated as follows: When the transformed yield record is represented as *X*, the GP distributions approximate the distribution of *X* − *T*_*h*_|*X* > *T*_*h*_ (the sign is inverted from the original yield record). Letting *y*_*m*_ denote the *M*-year return level, the probability that *X* exceeds *y*_*m*_ is 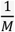 assuming that a yield record is obtained once per year. That is,

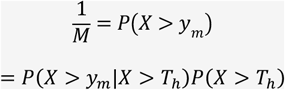

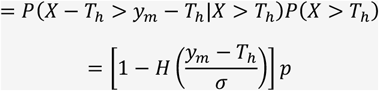

where *p* is *P*(*X* > *T*_*h*_) and *H* denotes the cumulative distribution function of the GP distribution, which is expressed as

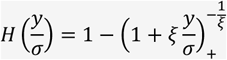

when *ξ* ≠ 0. Thus, we obtain

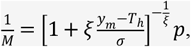

which yields

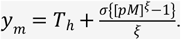

By replacing *ξ* with *nξ*, we obtain

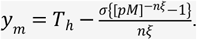

In the stationary GP models, *σ* and *nξ* were estimated with the MCMC algorithm, and *p* was empirically estimated as 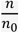 where *n* is the number of records that exceeded the threshold (*T*_*h*_) and *n*_*0*_ is the total number of records. In the non-stationary GP models, *σ*_*i*_ and *nξ*_*i*_ were calculated from the intercepts (*α*_0_ and *β*_0_) and slopes (*α*_1_ and *β*_1_) estimated with the MCMC algorithm. Because *σ*_*i*_ and *nξ*_*i*_ can change across years, the *M*-year return levels also can change across years. In the nonstationary GP models, *p* is also expected to change across years, whereas *T*_*h*_ is assumed to be constant across years. Thus, *p*_*i*_, *p* in year *i*, was estimated as follows assuming that *p*_*i*_ changes linearly across years and that the average of *p*_*i*_ across years is 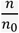. Let *z*_*i,k*_ = 1 if *y*_*i,k*_ > *T*_*h*_ and *z*_*i,k*_ = 0 otherwise, and *p*_*i*_ = *a* + *x*_*i*_*b* such that 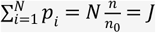. Then, the likelihood of the transformed yield records can be written as:

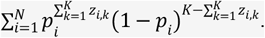

Because 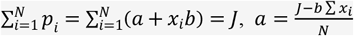. Thus,

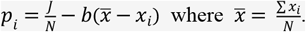

The slope (*b*) was estimated by maximizing the log likelihood

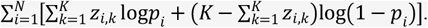

Because 0 < *p*_*i*_ < 1 for ∀*i*, the upper and lower bounds of *b* are constrained as follows. When *b* > 0, because 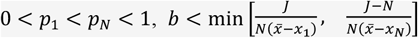. When *b* < 0, because 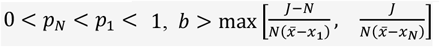. Thus, the bounds of *b* are

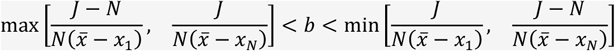

The value of *b* that maximized the log-likelihood was searched from 1,000 grids between these bounds, and *p*_*i*_ was estimated using the slope.

### 2.10. Simulation

Simulation analyses were conducted to investigate how accurately the risk of low yield (i.e., *M*-year return levels) could be estimated using real data analyses. The simulation schemes are as follows:

1. Chose the original data to be mimicked and calculate Pearson’s product-moment correlations of the transformed yield records among *K* countries/prefectures to create the *K*-by-*K* correlation matrix **C**.
2. Simulated *N*-year yield records for *K* countries/prefectures using *K*-dimensional multivariate normal distributions. The covariance matrix **Σ**_*i*_ of the multivariate normal distribution for year *i* was 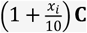. In other words, extreme events became more plausible as the years passed. The mean of the multivariate normal distribution is zero. If the original data contained missing records, the corresponding records were missing.
3. Apply the stationary and non-stationary GP models to the dataset created in (2) and obtain the *M*-year return level estimates. For *M*, 20 values of 10, 20, …, 190, and 200 were chosen.
4. Calculate the true *M*-year return levels from the normal distributions used to simulate the datasets in (2) and compare them with the estimates in (3).
5. Repeat (2) to (4) 100 times.

From the global yield data, wheat in ESeAsia_Oc, soybean in WCSAsia, and maize in Africa were chosen because the first two datasets were small and inference was expected to be difficult, whereas the last dataset was the largest. Specifically, for the wheat data in ESeAsia_Oc, *K* = 12 and the total number of records was 680, for the soybean data in WCSAsia, *K* = 17 and the total number of records was 679, and for the maize data in Africa, *K* = 53 and the tonal number was 3,043. All crops (wheat, rice, and soybeans) were selected from the local yield data. In the simulations of the local yield data, *K* = 47 for each crop, and the total number of records was 2,911 (wheat), 2,945 (rice), and 3,389 (soybean). The MCMC conditions were the same as those used for the real data in each simulation scenario.

### 2.11. Threshold

The accuracy of the approximation by the GP distributions and, consequently, the accuracy of estimating the *M*-year return levels, were expected to be influenced by the choice of thresholds (*T*_*h*_). However, it is difficult to select a single threshold value. Instead, in this study, a sequence of *T*_*h*_ from 0.5 to 2 were applied and the results were compared. In the real data analyses, the unit of these thresholds was the SD of the detrended yield, which was used to scale the yield data. In the simulation analyses, the units were the same as the simulated values.

## 3. Results

### 3.1. Yield data visualization

In Figure 2 are shown the histograms of the transformed (i.e., detrended, scaled, and sign-inverted) yield records pooled across years. The GP distributions approximate the upper tails of these distributions to estimate the risk of low yield. The Pearson correlation coefficients between countries/prefectures are shown in Figure 3. With respect to global yield data, the coefficients were almost zero on average, except in Europe, where weak mean correlations were observed. As for the local yield data, the correlation coefficients were moderate on average.

**Figure 2.**
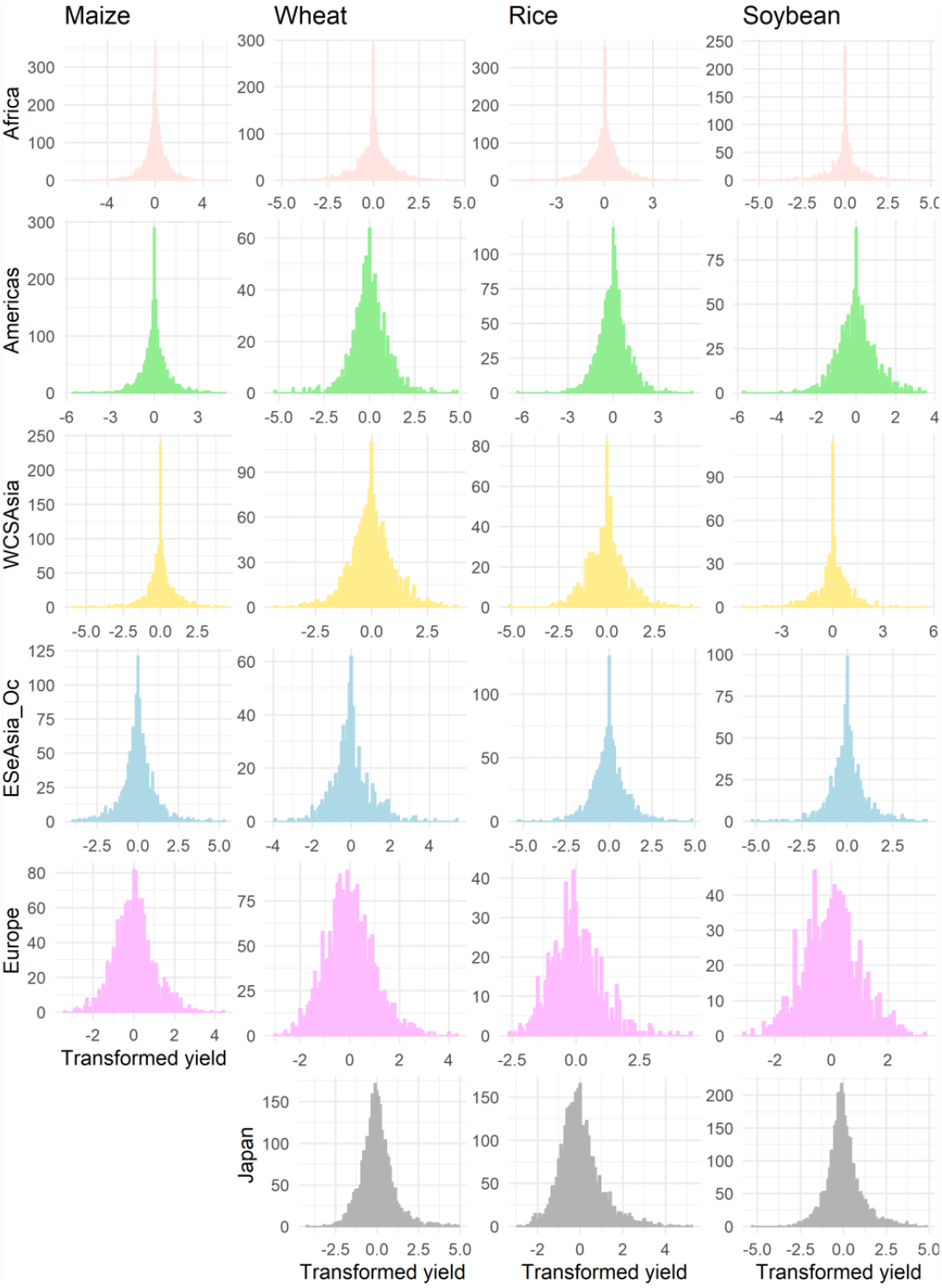
Histograms of transformed yield data. The units are the standard deviations of detrended yield records for each country/prefecture. The signs of original yield records were inverted after detrending and scaling. Thus, the low yield events are represented in the upper tails of the histograms. Africa, Americas, WCSAsia, ESeAsia_Oc, and Europe constitute the global yield data whereas Japan constitutes the local yield data. Data on maize yields in Japan is lacking as it is not a stable crop in Japan and the records were few. WCSAsia: Western, Central, and Southern Asia, ESeAsia_Oce: Eastern and Southern eastern Asia and Oceania.

**Figure 3.**
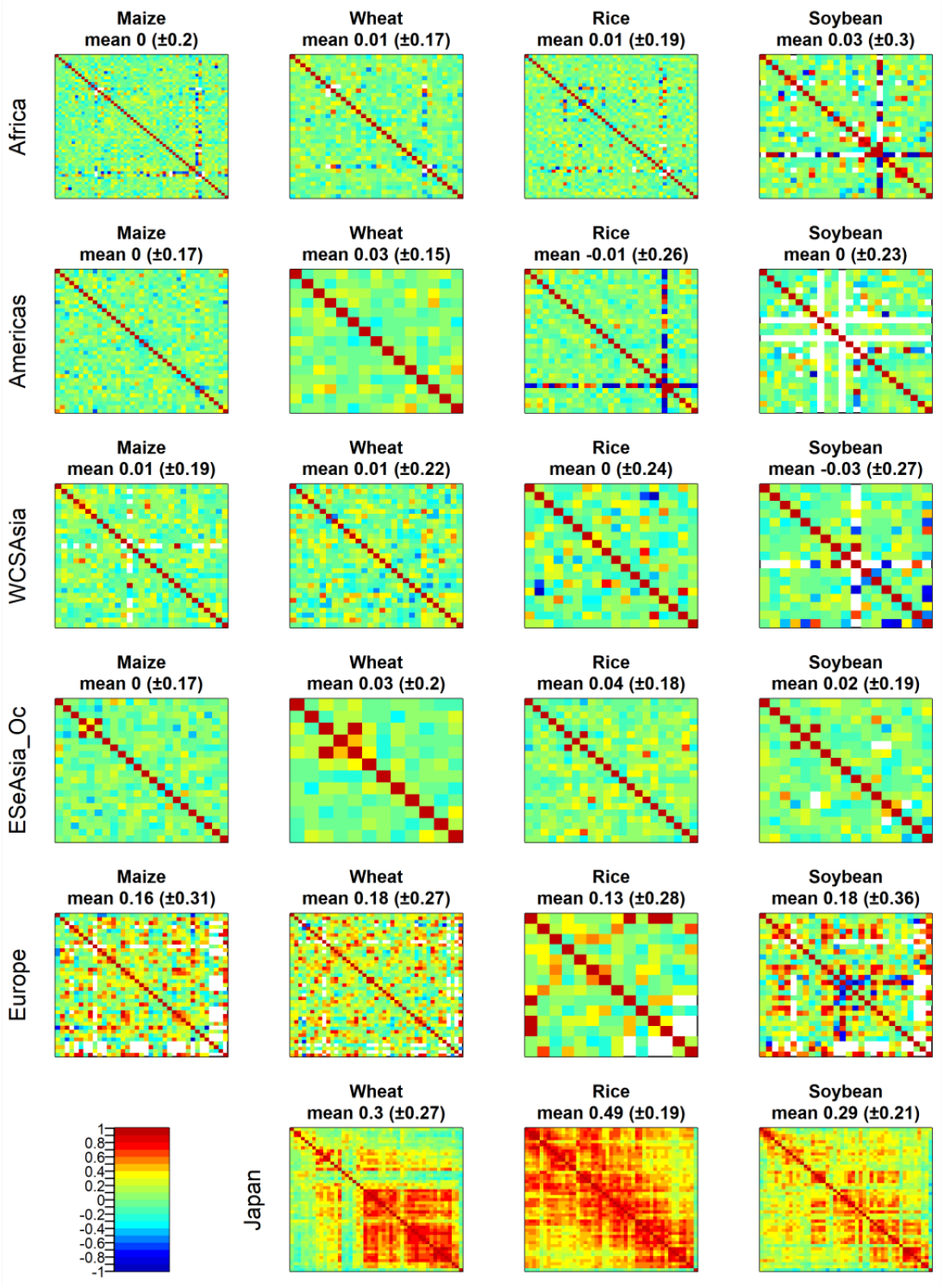
Heatmaps of the Pearson correlation coefficients of transformed yield data between areas (global yield data) or prefectures (local yield data of Japan). Titles of each panel include the mean correlation coefficients and the standard deviations. WCSAsia: Western, Central, and Southern Asia, ESeAsia_Oce: Eastern and Southern eastern Asia and Oceania.

### 3.2. Number of records used for approximation

Stationary and non-stationary GP models were applied to the transformed yield data that exceeded the threshold *T*_*h*_ to approximate the tail distributions. The number of records that exceeded the threshold decreased as the threshold increased. When *T*_*h*_ = 0.5, the average numbers of records were 336.1 (±142.6) for the global data and 730.7 (±32.2) for the local data (Tables S3 and S4). When *T*_*h*_ = 2.0, the numbers reduced to 42.8 (±20.4) and 119.3 (±13.0), respectively. Note that the unit of the thresholds is the SDs that were used for scaling the detrended data (see “2.3. Data processing”). As the threshold decreases, the approximation by the GP distribution can be inaccurate; however, as the threshold increases, the parameter estimation can be inaccurate because of the smaller sample size. To take balance in this trade-off relationship, multiple threshold values ranging from 0.5 to 2.0 with 0.1 interval were used and compared.

The chronological trends of the proportion of records that exceeded the thresholds are shown in Figure 4. The trends of low yield risk can be roughly understood from these plots. Interesting trends were observed for wheat and rice yield in Japan: for wheat, the frequencies of severe yield loss, such as 2 SD lower than averages, are decreasing, whereas those of loss with 0.5 SD below are constant or slightly increasing. In the case of rice, the frequency of severe yield losses increased until the 1990s, whereas the frequency of losses with 0.5 SD below average decreased. These issues and their influence on the results are discussed in Section 4.3. Interpreting the results of local yield data” in the Discussion section.

**Figure 4.**
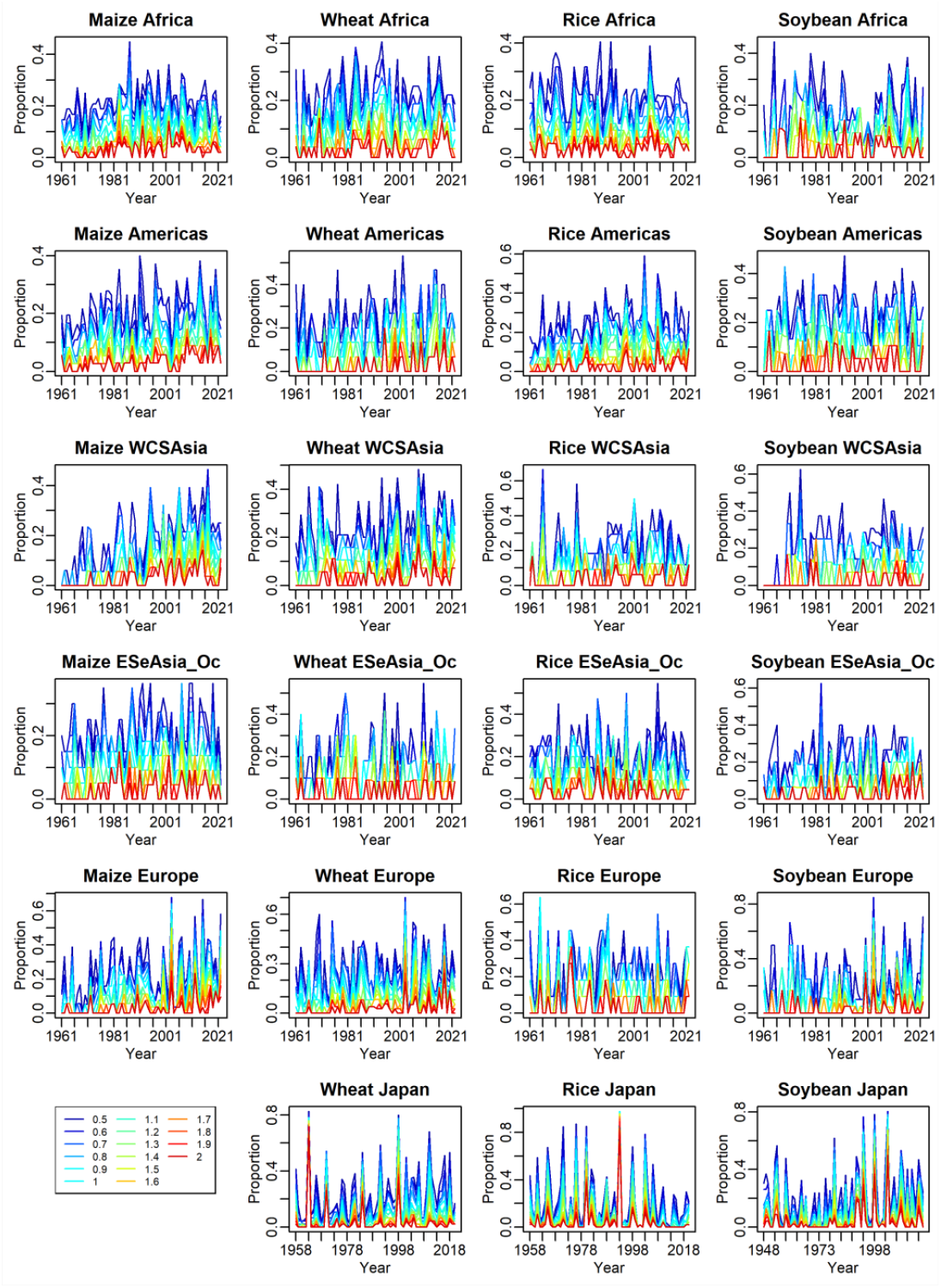
Proportions of yield records that exceeded the threshold values. The proportions were calculated for each year as the proportions of the transformed (sign-inverted) yield record that exceeded the threshold to the total number of records at the year. The proportions were colored according to the threshold values as shown in the legends at the left bottom panel. The y and x axes indicate the proportion of records and years of harvest, respectively. WCSAsia: Western, Central, and Southern Asia, ESeAsia_Oce: Eastern and Southern eastern Asia and Oceania. Maize of Japan

### 3.3. Parameter estimation via MCMC

Mean excess plots are presented in Figures S1–S4 (global yield data) and S5 (local yield data). All the plots showed decreasing trends, suggesting that the population distributions belonged to the domain of attractive of the Weibull distribution, that is, *ξ* < 0. Thus, the parameters of the stationary and nonstationary GP models were estimated based on this assumption.

The MCMC chains of stationary and non-stationary GP models showed good convergence. The 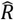 values are presented in Tables S5 and S6 (global yield data) and S7 and S8 (local yield data). The values were generally lower than 1.1 which was suggested as the convergence criterion by Gelman *et al* (2014). Several crop-area/prefecture combinations showed the values > 1.1 when the threshold *T*_*h*_ was high (Tables S5–S7), which was mainly due to the small sample sizes.

In the stationary GP models, the accuracy of the approximation can be assessed by quantile-quantile (QQ) plots between the empirical (observed) and approximated distributions (Figures S6–9 for the global yield data and Figures S10–12 for the local yield data). Although deviations from the 1:1 line were occasionally observed, particularly for extremely large values, the empirical and approximated distributions were generally concordant regardless of the threshold values. The QQ plots for the nonstationary GP models were not drawn because the approximated distributions change across years; thus, the empirical distribution to be compared consists of only one-year observations, and the number of records is small.

### 3.4. Model comparison

The WAIC values of the stationary and non-stationary GP models were compared (Figure 5). The WAIC tended to support stationary GP models as *T*_*h*_ increased (e.g., wheat in Americas and maize in Europe). This phenomenon probably occurred because higher *T*_*h*_ values masked chronologically increasing or decreasing trends by eliminating records of lower yields. This phenomenon was also observed in the simulations, as described later. When the non-stationary GP models were better to approximate the data, the slope parameters (*α*_1_ and *β*_1_) were expected to be significantly deviated from 0. Therefore, I judged that the non-stationary GP models were better than the stationary GP models when the WAIC values under *T*_*h*_ = 0.5 were smaller and the slope parameters deviated significantly from zero. As a result, the non-stationary GP models were judged as the best models in 13 combinations: maize in Africa; maize, wheat, and rice in Americas; maize, wheat, and rice in WCSAsia; soybean in ESeAsia_Oc; maize and wheat in Europe; and all crops in Japan (Figure 5). The signs of the slope parameter *α*_1_ were generally plus, which means that the tails of the GP distributions become thicker, that is, low yield risks increased. Exceptions were observed for rice in the WCS Asia, wheat in Japan, and rice in Japan. For wheat in Japan, *α*_1_ was significantly negative under any *T*_*h*_, suggesting a clear trend of mitigating risk.

**Figure 5.**
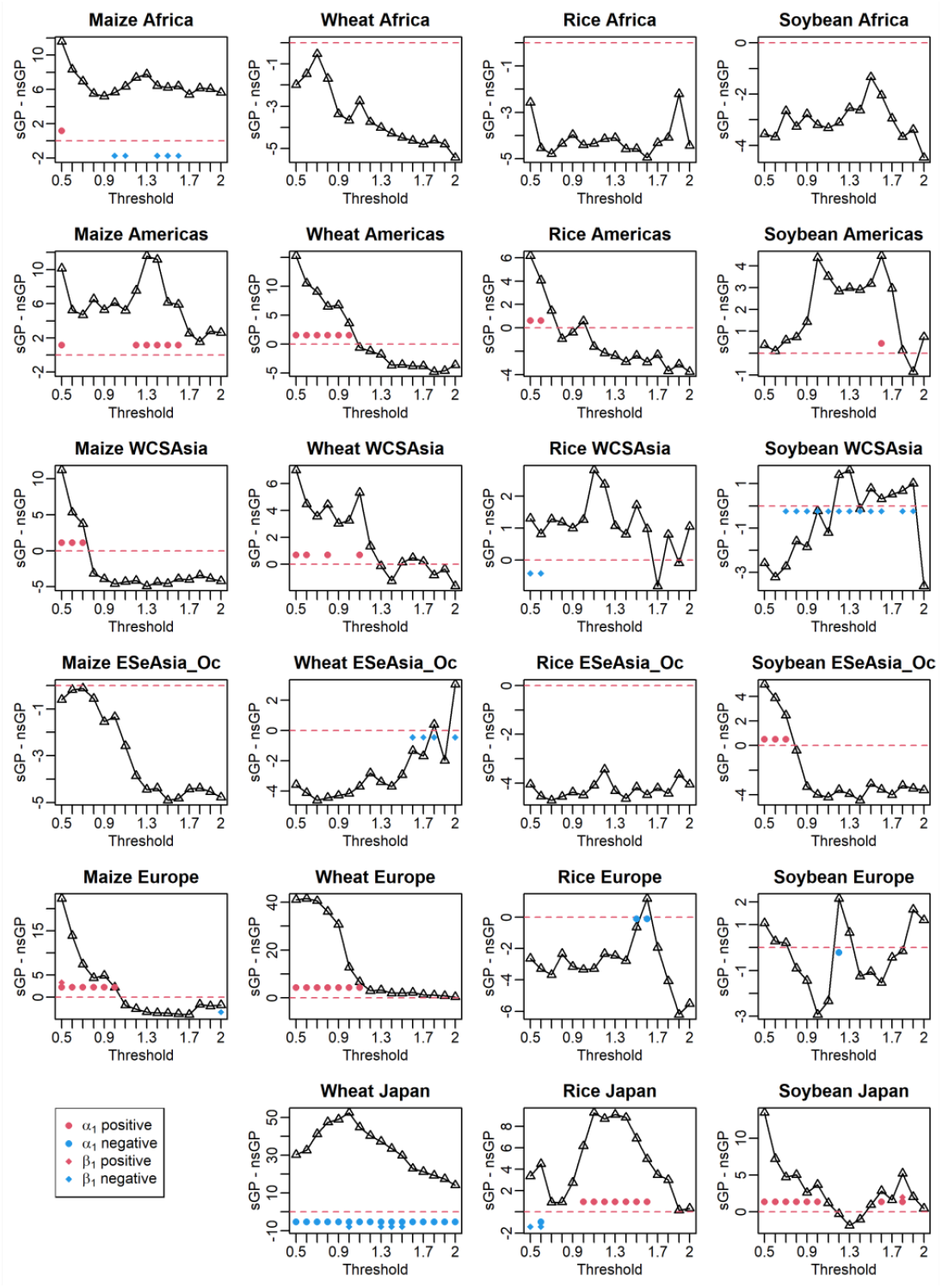
Differences of widely applicable information criteria (WAIC) between stationary generalized Pareto (sGP) models and non-stationary generalized Pareto (nsGP) models. The y and x axes indicate the differences in WAIC values and the threshold values, respectively. The red broken lines indicate zero. Values greater than zero suggest that the nsGP models were preferable to approximate the distributions of low yield records. Dots and diamonds indicate that the *α*_1_ and *β*_1_ parameters significantly deviated from zero with 95% highest density intervals, respectively, and red and blue colors indicate that the parameters significantly deviated from zero to positive and negative directions, respectively. WCSAsia: Western, Central, and Southern Asia, ESeAsia_Oce: Eastern and Southern eastern Asia and Oceania. Maize of Japan

### 3.5. *M*-year return level

In both the global and local yield data, the *M*-year return levels estimated using different *T*_*h*_ values were largely consistent among the values (Figures S 13–17). However, estimates occasionally showed unrealistic curves when *T*_*h*_ was high, which was probably due to estimation failure caused by small sample sizes (e.g., wheat in Americas and rice in Europe). In addition, as shown later, simulation analyses suggest that high *T*_*h*_ values can mask chronological trends when the risk of low yield changes over time. Therefore, a moderate value of 1.0 was adopted as *T*_*h*_ in order to assess the chronological trends of risk. In the case of maize in WCSAsia, the probability of exceeding *T*_*h*_ = 1.0 was lower than 1/200 where 200 is the maximum *M* value; as for soybean in this region, the *M*-return levels became unrealistic when the threshold was 1.0 (Figure S16). Therefore, *T*_*h*_ = 0.5 was adopted for this area. The *M*-year return levels estimated at *T*_*h*_ = 1.0 or 0.5 with non-stationary GP models are presented in Figure 6. The return levels at specific *M* values are listed in Tables S9 and S10. The return levels estimated for the oldest and latest years in the data are superimposed. These return levels are reported in the original signs, meaning that lower values suggest a more severe risk of low crop yields. Among the 13 crop-area/prefecture combinations where non-stationary GP models were judged as the best models, nine combinations (maize in Africa; maize, wheat, and rice in Americas; maize and wheat in WCSAsia; maize and wheat in Europe; and soybean in Japan) showed that the return levels worsened (decreased) as years passed, as represented by the non-overlapping highest density intervals between the oldest and most recent years (Figure 6). Such a clear trend was not observed for rice in WCSAsia or soybean in ESeAsia_Oc because of the large uncertainty. With respect to rice in Japan, the return levels of the oldest and latest years crossed. As expected from the slope parameter estimates, return levels for wheat in Japan were mitigated (increased) over time. The return levels of rice and wheat in Japan are discussed in Section 4.3. These results of the local yield data are interpreted using the unique trends presented in Figure 4.

**Figure 6.**
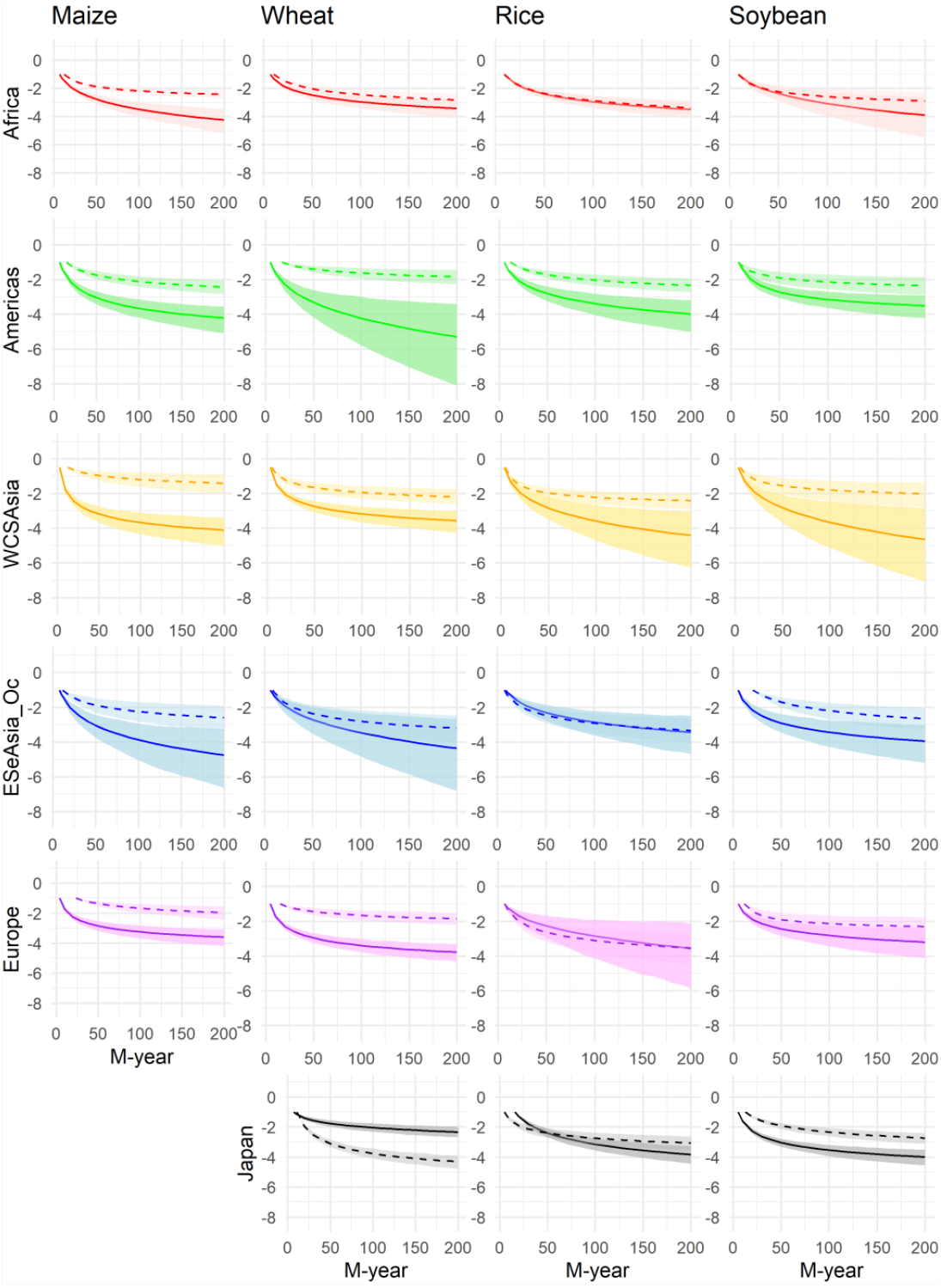
The *M*-year return levels of low yield risks estimated with the non-stationary generalized Pareto models. The threshold was 1.0 except for WCSAsia which adopted 0.5. The unit of y axis is the standard deviations of detrended yield records for each country/prefecture. The return levels are presented in original signs meaning that lower values suggest severer low yield risks. The broken and solid lines indicate the posterior means of *M*-year return levels estimated at the oldest and latest years included in each data, respectively. The oldest and latest years of the global yield data were 1961 and 2022, respectively, whereas these of the local yield data (Japan) were 1958 and 2020 for wheat and rice, and 1948 and 2020 for soybean. The shades indicate the 95% highest density intervals. WCSAsia: Western, Central, and Southern Asia, ESeAsia_Oce: Eastern and Southern eastern Asia and Oceania. Maize of Japan

Although the estimated risk is represented by the scale of the standard deviation (Figure 6), the estimates can be converted into units of observations. For example, the 30-year return level of maize in Europe in 2022 was estimated to be – 2.53 SD (Table S9). By multiplying the standard deviations of the detrended yield records (Table S1), the estimates at the original scale (100 g/ha) can be obtained for each country, such as – 15,177.5 in France and – 12,948.1 in Italy. This information will be beneficial for practical applications.

### 3.6. Simulations

Among the six simulation scenarios analyzed, three mimicked the global yield data (wheat in ESeAsia_Oc, soybean in WCSAsia, and maize in Africa), and three mimicked all crops of the local yield data in Japan. In each simulation scenario, the risk of low yield was assumed to worsen over time; thus, the non-stationary GP models were expected to explain the data better than the stationary GP models. The simulated yield data are summarized in Figure S18, which shows gradually increasing trends in the yield variability over time. The WAIC generally supported the non-stationary GP models when the *T*_*h*_ values were low (Figure S19). The differences in WAIC between the stationary and nonstationary GP models decreased as *T*_*h*_ increased, probably because the chronological trends were masked by high *T*_*h*_ values, as observed in the real data analyses. The means of *M*-year return levels estimated with the non-stationary GP models were consistent with the true values (Figures S20–S25), although a slight overestimation was observed in the older and later years. Overestimation became visible as the *T*_*h*_ values increased (Figures S20–25). Overestimation is reasonable because higher thresholds mean the elimination of smaller yield records and an increase in the overall risk level. By comparing the results between the simulated maize in Africa and the other two scenarios for the global yield data (wheat in ESeAsia_Oc and soybean in WSCAsia), the estimation became more precise as the number of records increased (Figure S22 vs. Figures S20 and S21).

## 4. Discussion

### 4.1. Estimated trends of low yield risk

The analysis of the global yield data revealed that the yield of maize and wheat crops showed worsening trends in multiple areas (Africa, Americas, WCSAsia, and Europe for maize, and Americas, WCSAsia, and Europe for wheat). The worsening trends in maize and wheat production have been also reported in previous studies. For example, Iizumi and Ramankutty (2016) examined the yield variability of maize, wheat, rice, and soybean in 1981–2010 using standard deviations of yield and found that the increasing variability of maize and wheat yield in Europe. Liu *et al* (2021) simulated future wheat yield using crop model emulators and found that the coefficients of variation (CV) in Europe and South America increase. Haqiqi (2024) estimated the low yield risk of maize based on statistical models using temperature and precipitation as explanatory variables and found that the risk is worsened in almost all regions in the case of rainfed crops; even in the case of irrigation, the risk is worsened in most areas other than major breadbaskets such as France, Germany, and Canada. Vogel *et al* (2019) reported that 18–43% of the variance in yield anomalies was explained by climate extremes, and North America for maize, spring wheat, and soybean; Asia for maize and rice; and Europe for spring wheat production were particularly susceptible to climate extremes. These consistent results support the validity of the proposed model and GP distribution-based approaches.

However, not all the results of the present study are consistent with the abovementioned previous studies. For example, the risk of wheat in ESeAsia_Oc did not change significantly in the present study (Figure 6), whereas the yield variability of wheat in most regions of Australia increased significantly (Iizumi and Ramankutty, 2016). This inconsistency can be attributed to the difference in study area units; the present study targeted rough areas of global scales (e.g., Europe and Americas), whereas the previous study targeted finer regions of specific areas. Thus, trends observed locally may not be detected in the present study.

### 4.2. Spatial dependency between yield records

As described in “2.1. Global yield data” in the Methods section, the study areas (Africa, Americas, WCSAsia, ESeAsia_Oc, and Europe) were defined such that the number of included countries was sufficient for each area, which was necessary for a stable statistical inference. The simulation results also show that the number of records is a key factor for precise estimation (Figures S20–S22). However, including many countries/regions will result in wide study area units, which will dismiss the realistic usefulness of the estimated risks. It is possible to narrow the area unit if longitudinal yield records from finer regions (e.g., prefectures/counties) are included, as is done for the local yield records of Japan, while keeping the number of records large enough. In this case, the spatial dependencies of the yield records between regions emerge (Figure 3). Spatial dependency causes an underestimation of risk uncertainty, while dependency does not seem to affect the estimation of means, as shown in the simulations (Figures S23–25). The estimation precision was also affected by spatial dependency owing to the decreasing effective sample size. This was confirmed by comparing the simulation results for maize in Africa (Figure S22) with those for all crops in Japan (Figures S23–25). Although the total number of yield records was comparable, the estimated precision of maize in Africa was narrower than that of all crops in Japan, which showed stronger spatial dependency (Figure 3). The application of GP models to spatially dependent yield records was also attempted by Mitchell *et al* (2020) who estimated the theoretical maximum wheat yield in England using GP distributions. Nevertheless, it must be acknowledged that increasing the number of records results in wider study areas, whereas narrowing study areas yields spatial dependency. This trade-off is a disadvantage of the current model; a possible remedy is to estimate the GP model parameters (*ξ* and *σ*) for each country/prefecture allowing spatial heterogeneity using hierarchical models (Park et al. 2019, Momoki and Yoshida 2024).

Another issue to be addressed is that the low-yield risk estimation with GP distributions does not consider the lowest value of yield, that is, 0. Therefore, the estimated yield can be below this limit. For example, the 200-year returns were estimated as approximately −2 to −5 SDs, depending on the combinations of crop and country/prefecture. The estimated return levels were below the limit in 26 (4.7%) crop-country combinations in the global yield data and two (1.4%) crop-prefecture combinations in the local yield data (Tables S11 and S12). Because the global yield data is more spatially heterogeneous, such invasion would be more frequently observed in the global yield data. Further improvements to the model to allow for estimation of the GP model parameters unique to each member of the study unit (country/prefecture) would mitigate this problem.

### 4.3. Interpreting the results of local yield data

As mentioned in the Results section, the trends in the risk of low yield in wheat and rice crops in Japan is noteworthy. In the case of wheat, the frequency of severe yield decreased in the study period, whereas the frequency of losses lower than 0.5 SD were constant or slightly increased; as for rice, the frequency of severe yield loss increased until the 1990s, whereas frequency of losses lower than 0.5 SD decreased (Figure 4). These trends affected the estimated GP distributions, which approximated the lower tails of yield (Figure 7).

**Figure 7.**
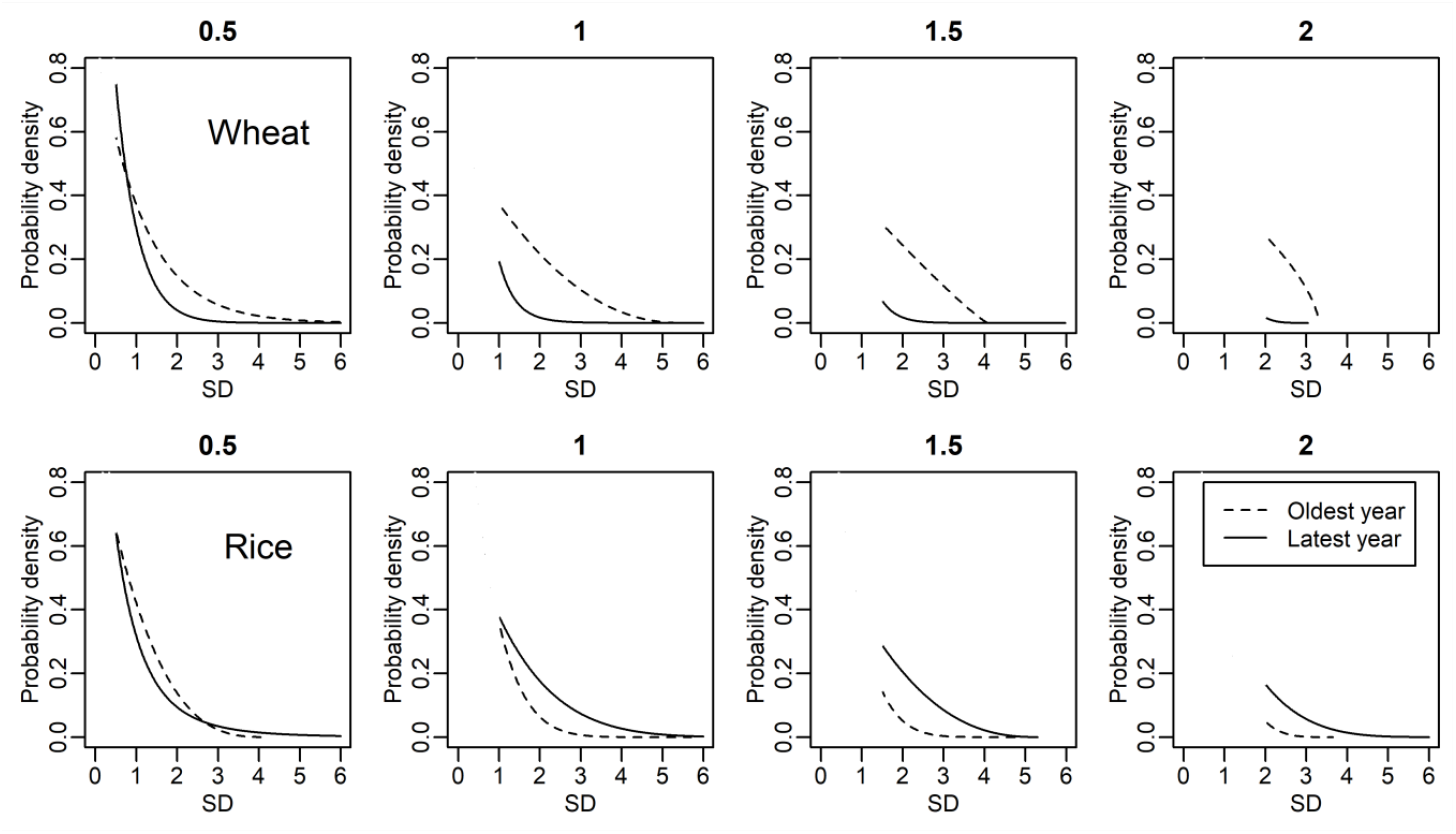
The probability densities of the generalized Pareto distributions estimated for wheat and rice of the local yield data in Japan. The upper and lower panels are for wheat and rice, respectively. The broken and solid lines indicate the density curves estimated at the oldest (1958) and latest (2020) years, respectively. The plot titles represent the threshold (*T*_*h*_) values.

In the case of wheat in Japan, the GP distributions that were estimated with *T*_*h*_ = 0.5 were close between the oldest (1958) and latest (2020) years, with a slightly ticker tail of the oldest year, reflecting the decreasing trend of severe yield loss (Figure 4). The differences increased as *T*_*h*_ increased (Figure 7) because the records used for estimation were dominated by severe cases that decreased over time (Figure 4). Consequently, the differences in the return levels between the oldest and latest years also increased as *T*_*h*_ increased (Figure S17).

For rice in Japan, the GP distributions estimated with *T*_*h*_ = 0.5 were also close between the oldest and latest years (Figure 7); however, the latest one showed a slightly ticker tail reflecting the increasing trend of severe yield loss (Figure 4), whereas the oldest one showed higher densities of less than 2 SD, reflecting the decreasing trend of slight-to-moderate yield loss (Figure 4). As *T*_*h*_ increased, the density of the latest year became higher than that of the oldest year in any range (Figure 7), which reflects the increasing trend of severe yield loss (Figure 4). Thus, for rice, the shapes of the density curves differed significantly, depending on the threshold (*T*_*h*_) (Figure 7). This was also reflected in the estimated parameter values; as presented in Figure 5, *α*_1_ and *β*_1_ were estimated as negative, but *α*_1_ became positive as *T*_*h*_ increased. The opposing trends in frequencies between slight-to-moderate and severe yield losses would make the return levels of the oldest and latest years cross; that is, short-term risks were higher for the oldest year, but the long-term risks were higher for the latest year. Crosses were also observed for *T*_*h*_ values other than 1.0 (Figure S17).

## 5. Conclusion

In the present study, non-stationary GP models were developed within a Bayesian framework to estimate the risk of low yields of staple crops, and the model was tested with both actual and simulated yield data. The proposed model reliably estimates changes in the risk of low yield over time. The MCMC chains generally showed good convergence unless the data size was small, owing to stringent threshold values (*T*_*h*_), suggesting that the proposed methods are computationally practical. Although the proposed model requires a combination of longitudinal yield data from multiple regions within an area of interest, the estimated risks can be interpreted on the scale of each individual region. As illustrated by the simulation analyses, estimation precision depends on the number of samples, which can be increased by including more regions. However, including more regions in an area of interest also results in stronger spatial dependencies, which may affect the estimation of uncertainty. This issue can be addressed by improving the model to account for spatial dependencies. Although there is still room for improvement, the present study demonstrates that it is possible to estimate low-yield risks that are chronologically changing using a data-driven approach without assumptions about climate and crop physiology.

## Acknowledgements

The authors would like to thank Ryukoku University for providing financial support.

## References

Ceglar A, Toreti A, Lecerf R, Van der Velde M and Dentener F 2016 Impact of meteorological drivers on regional inter-annual crop yield variability in France. Agric. For. Meteorol. 216 58–67. 10.1016/j.agrformet.2015.10.004

Chou J, Jin H, Xu Y, Zhao W, Li Y and Hao Y 2024 Impacts and risk assessments of climate change for the yields of the major grain crops in China, Japan, and Korea. Foods 13 966. 10.3390/foods13060966

Coles S, Bawa J, Trenner L and Dorazio P 2001 An introduction to statistical modeling of extreme values. London, Springer

Department of Economic and Social Affairs, Statistics Division, United Nations 1999 Standard country or area codes for statistics use (revision 4). https://unstats.un.org/unsd/publication/SeriesM/Series_M49_Rev4(1999)_en.pdf

Deressa T T, Hassan R M, Ringler C, Alemu T and Yesuf M 2009 Determinants of farmers’ choice of adaptation methods to climate change in the Nile Basin of Ethiopia. Glob. Environ. Change. 19 248–55. 10.1016/j.gloenvcha.2009.01.002

Eddelbuettel D and Balamuta J J 2018 Extending R with C++: a brief introduction to Rcpp. Am. Stat. 72 28–36. 10.1080/00031305.2017.1375990

Faye B, Webber H, Naab J B, MacCarthy D S, Adam M, Ewert F, Lamers J P A, Schleussner C, Ruane A and Gessner U 2018 Impacts of 1.5 versus 2.0 °C on cereal yields in the West African Sudan Savanna. Environ. Res. Lett. 13 034014. 10.1088/1748-9326/aaab40

Funk C, Dettinger M D, Michaelsen J C, Verdin J P, Brown M E, Barlow M and Hoell A 2008 Warming of the Indian Ocean threatens eastern and southern African food security but could be mitigated by agricultural development. Proc Natl Acad Sci U S A. 105 11081–6. 10.1073/pnas.0708196105

Gelman A, Carlin J B, Stern H S, Dunson D B, Vehtari A and Rubin D B 2014 Bayesian Data Analysis (Third Edition) CRC press, London

Haqiqi I 2024 Trade can buffer climate-induced risks and volatilities in crop supply. Environ. Res.: Food Syst. 1 021004 10.1088/2976-601X/ad7d12

Howden S M, Soussana J, Tubiello F N, Chhetri N, Dunlop M and Meinke H 2007 Adapting agriculture to climate change. Proc Natl Acad Sci U S A. 104 19691–6. 10.1073/pnas.07018901

Hsiao J, Kim S, Timlin D J, Mueller N D and Swann A L S 2024 Model-aided climate adaptation for future maize in the US. Environ. Res.: Food Syst. 1 015004. 10.1088/2976-601X/ad3085

Iizumi T and Ramankutty N 2016 Changes in yield variability of major crops for 1981–2010 explained by climate change. Environ. Res. Lett. 11 034003. 10.1088/2976-601X/ad3085

Jüttner U, Peck H and Christopher M 2003 Supply chain risk management: outlining an agenda for future research. Int. J. Logist. 6 197–210. 10.1080/13675560310001627016

Kamali B, Jahanbakhshi F, Dogaru D, Dietrich J, Nendel C and AghaKouchak A 2022 Probabilistic modeling of crop-yield loss risk under drought: A spatial showcase for sub-Saharan Africa. Environ. Res. Lett. 17 024028. 10.1088/1748-9326/ac4ec1

Leng G and Hall J W 2020 Predicting spatial and temporal variability in crop yields: an inter-comparison of machine learning, regression and process-based models. Environ. Res. Lett. 15 044027. 10.1088/1748-9326/ab7b24

Liu W, Ye T, Jägermeyr J, Müller C, Chen S, Liu X and Shi P 2021 Future climate change significantly alters interannual wheat yield variability over half of harvested areas. Environ. Res. Lett. 16 094045. 10.1088/1748-9326/ac1fbb

Mitchell E G, Crout N M J, Wilson P, Wood A T A and Stupfler G 2020 Operating at the extreme: estimating the upper yield boundary of winter wheat production in commercial practice. R. Soc. Open Sci. 7 191919. 10.1098/rsos.191919

Momoki K and Yoshida T 2024 Unit-level mixed effects models for conditional extremes. arXiv 2305.05106v3, 10.48550/arXiv.2305.05106

Müller C, Franke J, Jägermeyr J, Ruane A C, Elliott J, Moyer E, Heinke J, Falloon P D, Folberth C and Francois L 2021 Exploring uncertainties in global crop yield projections in a large ensemble of crop models and CMIP5 and CMIP6 climate scenarios. Environ. Res. Lett. 6 034040 10.1088/1748-9326/abd8fc

Park E, Brorsen B W and Harri A 2019 Using Bayesian Kriging for spatial smoothing in crop insurance rating. Am. J. Agric. Econ. 101 330–51. 10.1093/ajae/aay045

Rathore L S, Kumar M, Moftakhari H and Ganguli P 2024 Divergent changes in crop yield loss risk due to droughts over time in the US. Environ. Res. Lett. 19 114008. 10.1088/1748-9326/ad7618

Ray D K, Gerber J S, MacDonald G K and West P C 2015 Climate variation explains a third of global crop yield variability. Nat. Commun. 6 5989. 10.1038/ncomms6989

Sultan B, Guan K, Kouressy M, Biasutti M, Piani C, Hammer G L, McLean G and Lobell D B 2014 Robust features of future climate change impacts on sorghum yields in West Africa. Environ. Res. Lett. 9 104006 10.1088/1748-9326/9/10/104006

Tang C S 2006 Perspectives in supply chain risk management. Int. J. Prod. Econ. 103 451–88. 10.1016/j.ijpe.2005.12.006

Vogel E, Donat M G, Alexander L V, Meinshausen M, Ray D K, Karoly D, Meinshausen N and Frieler K 2019 The effects of climate extremes on global agricultural yields. Environ. Res. Lett. 14 054010. 10.1088/1748-9326/ab154b

Watanabe S 2010 Asymptotic equivalence of Bayes cross validation and widely applicable information criterion in singular learning theory. J. Mach. Learn. Res. 11 3571–94. http://jmlr.org/papers/v11/watanabe10a.html

Zampieri M, Ceglar A, Dentener F and Toreti A 2017 Wheat yield loss attributable to heat waves, drought and water excess at the global, national and subnational scales. Environ. Res. Lett. 12 064008. 10.1088/1748-9326/aa723b

