## Supplementary Figures for "Estimating the changing risks of low crop yield using non-stationary generalized Pareto distributions"

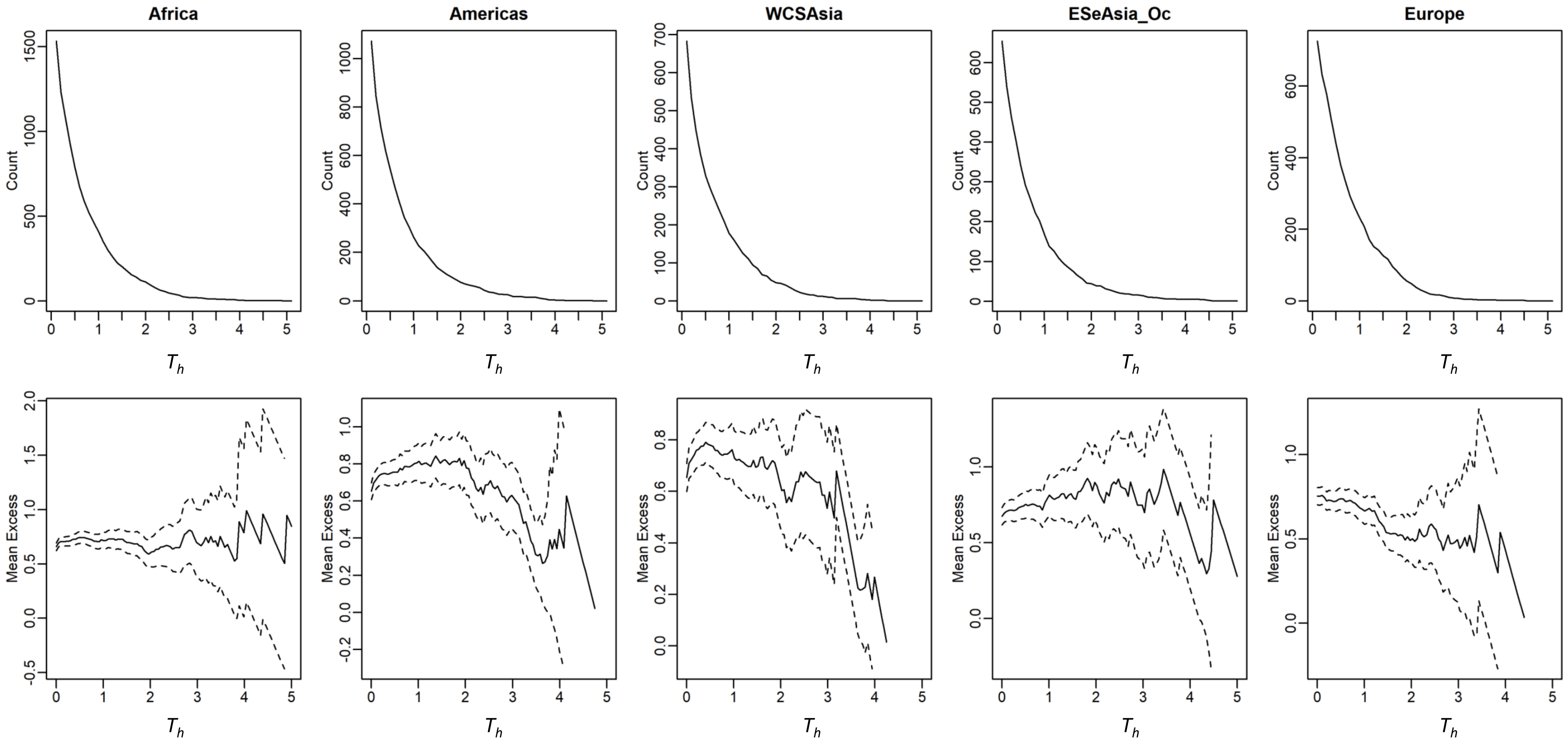


**Figure S1** Mean excess plots of maize in the global yield data. The top panels are the number of records (Count) that exceeded the thresholds ($T_h$). The bottom panels are the mean excess plots. The broken lines indicate the 95% confidence intervals. WCSAsia: Western, Central, and Southern Asia; ESeAsia_Oc: Eastern and Southeastern Asia and Oceania

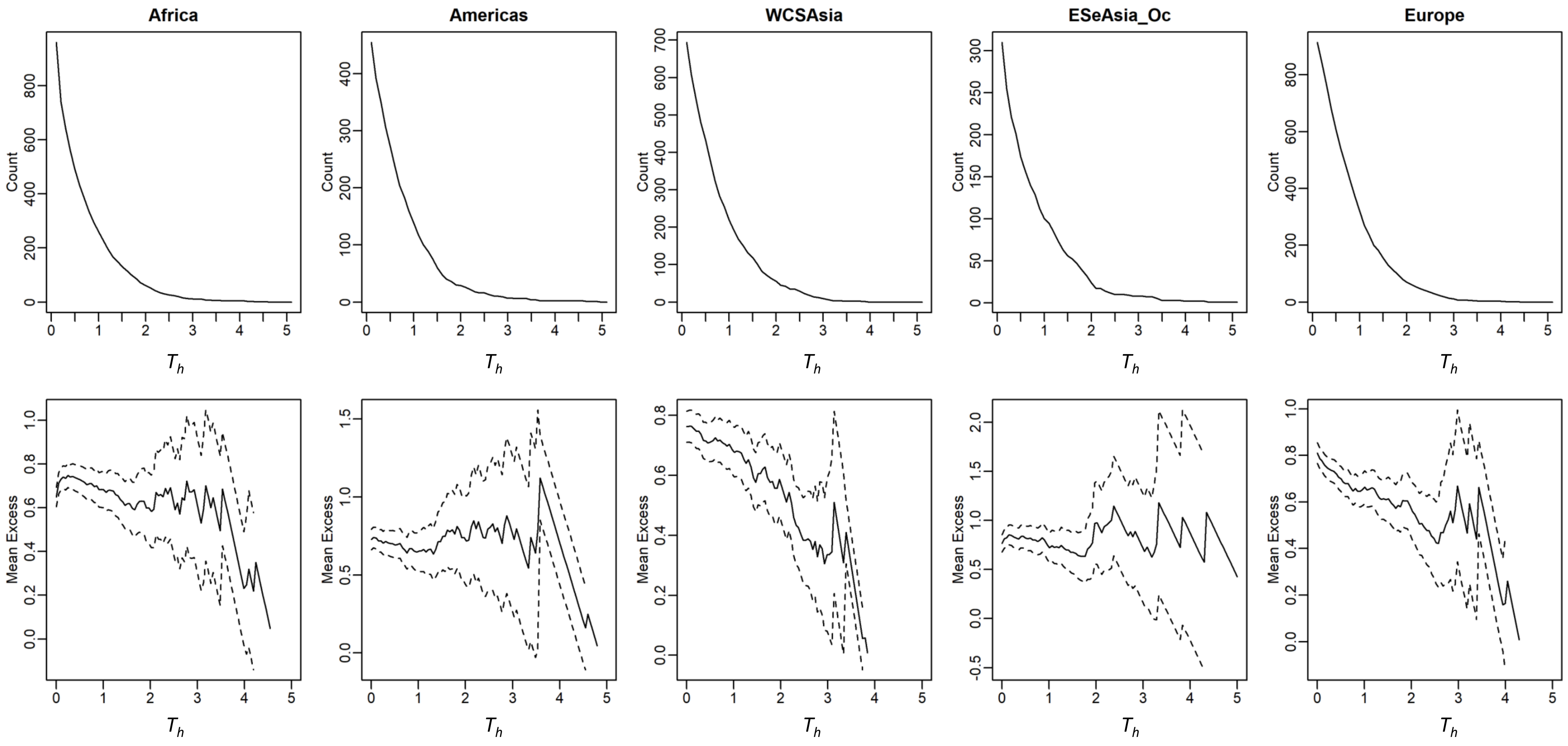


**Figure S2** Mean excess plots of wheat in the global yield data. The top panels are the number of records (Count) that exceeded the thresholds ($T_h$). The bottom panels are the mean excess plots. The broken lines indicate the 95% confidence intervals. WCSAsia: Western, Central, and Southern Asia; ESeAsia_Oc: Eastern and Southeastern Asia and Oceania

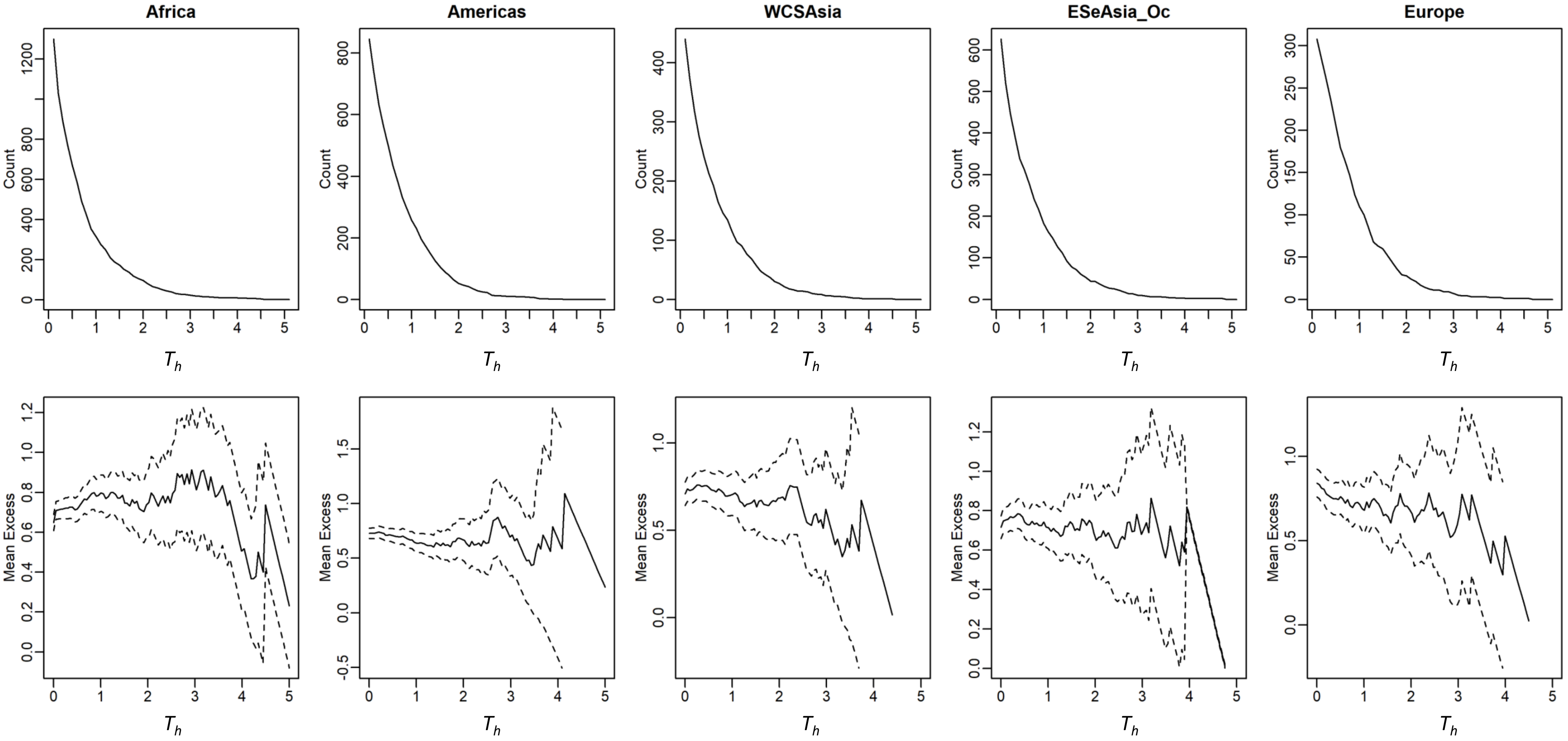


**Figure S3** Mean excess plots of rice in the global yield data. The top panels are the number of records (Count) that exceeded the thresholds ($T_h$). The bottom panels are the mean excess plots. The broken lines indicate the 95% confidence intervals. WCSAsia: Western, Central, and Southern Asia; ESeAsia_Oc: Eastern and Southeastern Asia and Oceania

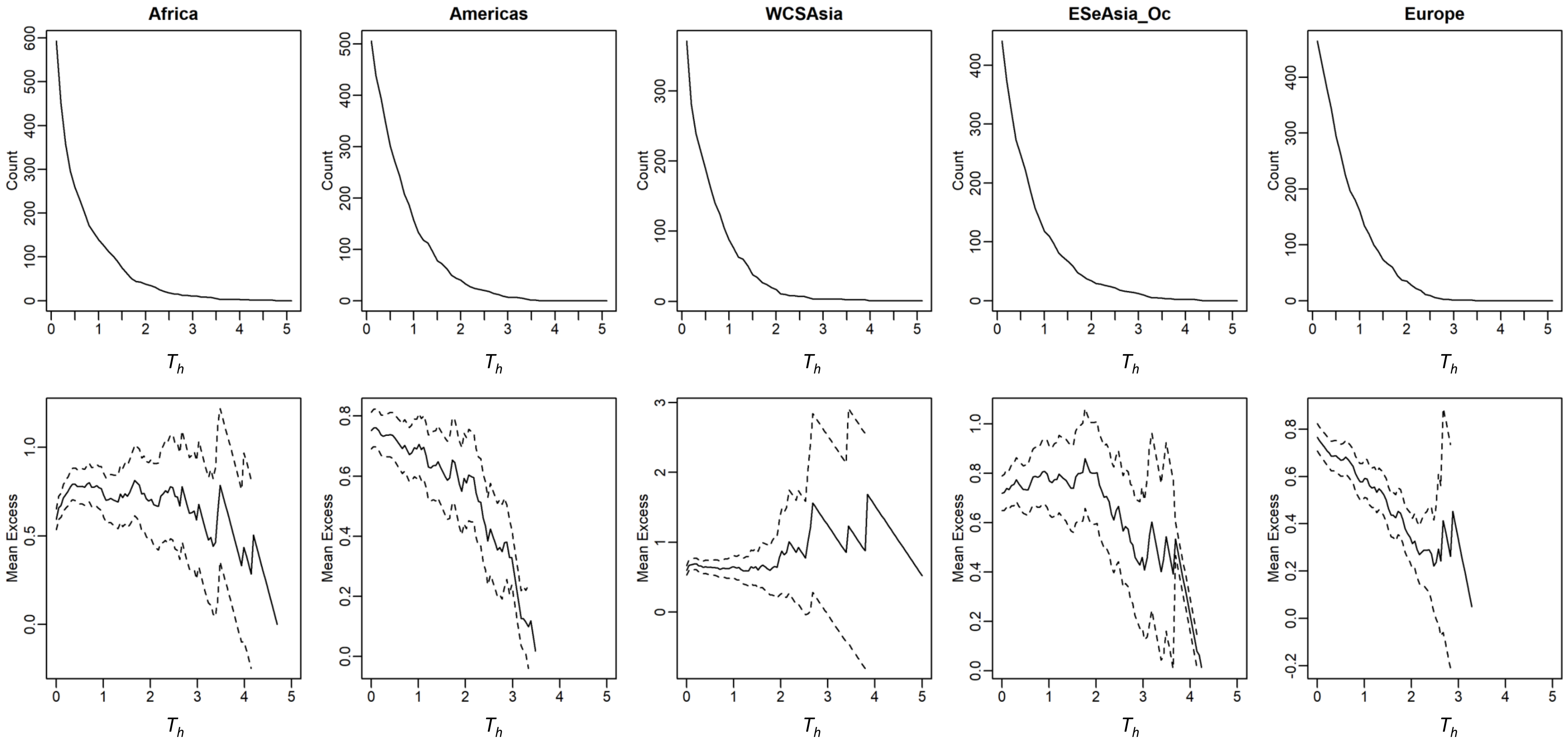


**Figure S4** Mean excess plots of soybean in the global yield data. The top panels are the number of records (Count) that exceeded the thresholds ($T_h$). The bottom panels are the mean excess plots. The broken lines indicate the 95% confidence intervals. WCSAsia: Western, Central, and Southern Asia; ESeAsia_Oc: Eastern and Southeastern Asia and Oceania

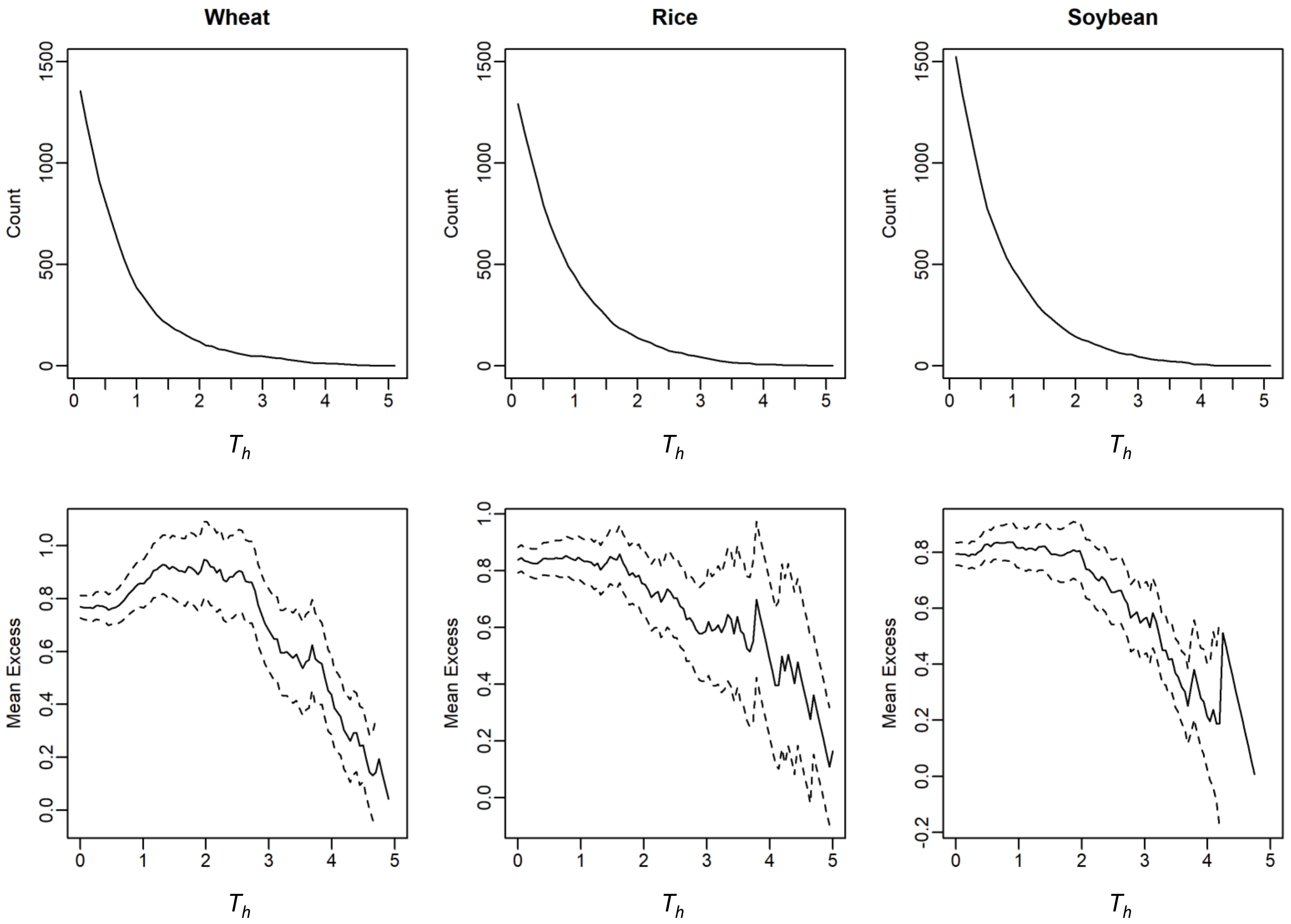


**Figure S5** Mean excess plots for the local yield data. The top panels are the number of records (Count) that exceeded the thresholds ($T_h$). The bottom panels are the mean excess plots. The broken lines indicate the 95% confidence intervals.

(A) Africa

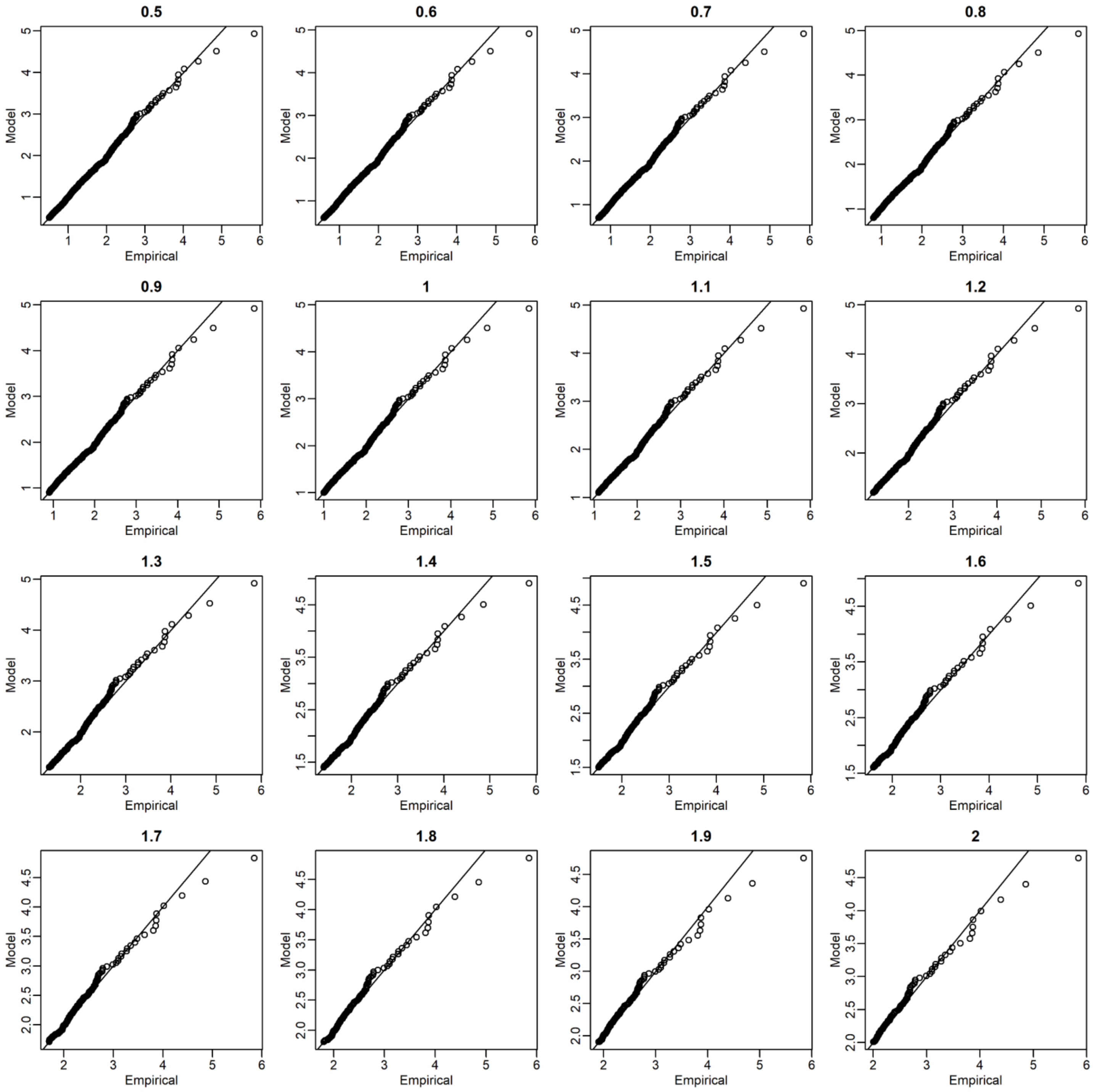


**Figure S6** Quantile-quantile (QQ) plots for maize of the global yield data between the empirical (observed) and approximated distributions. The empirical distributions were approximated with the stationary generalized Pareto models. The titles of plots denote the threshold values ($T_h$).

(B) Americas

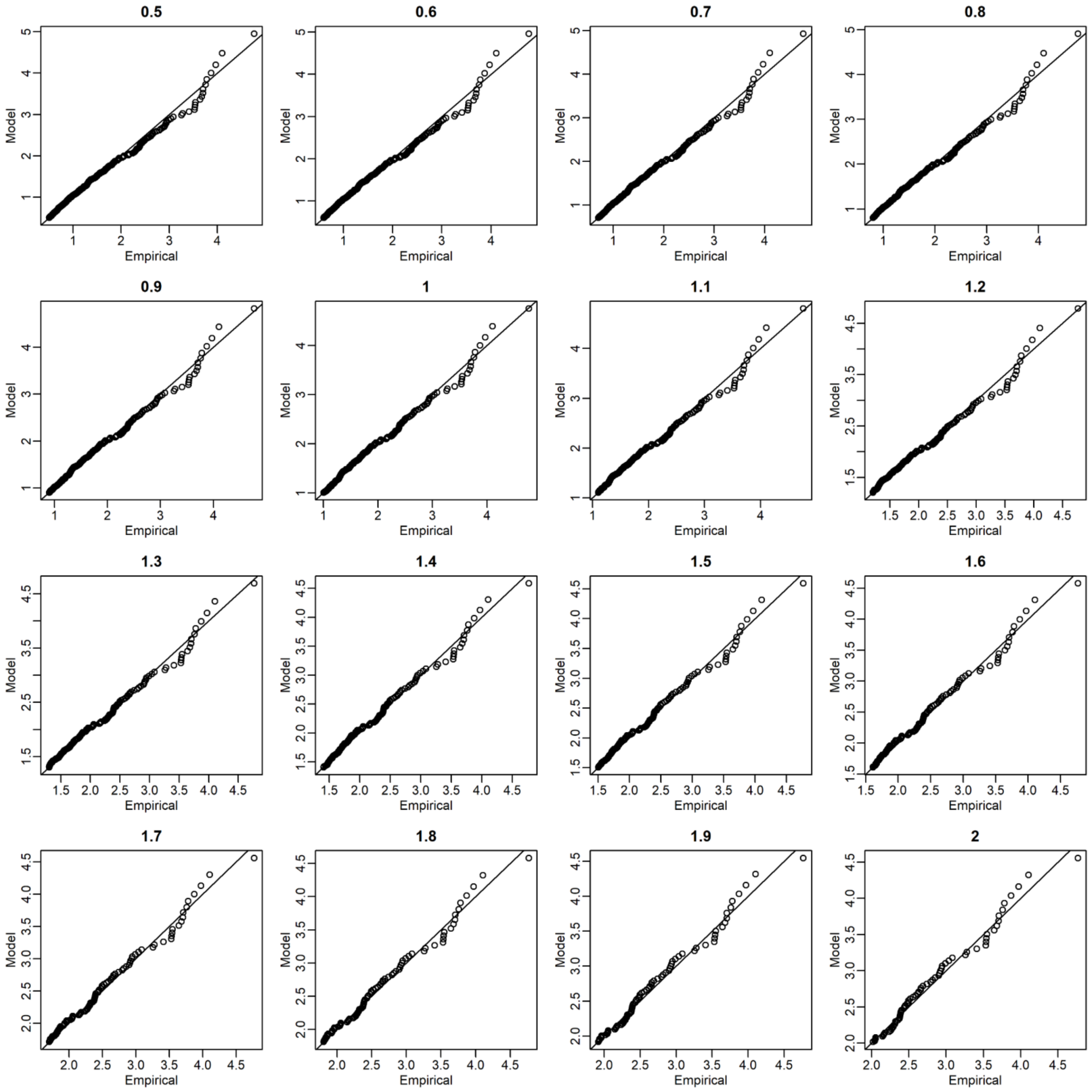


Figure S6 continued

(C) WCSAsia (Western, Central, and Southern Asia

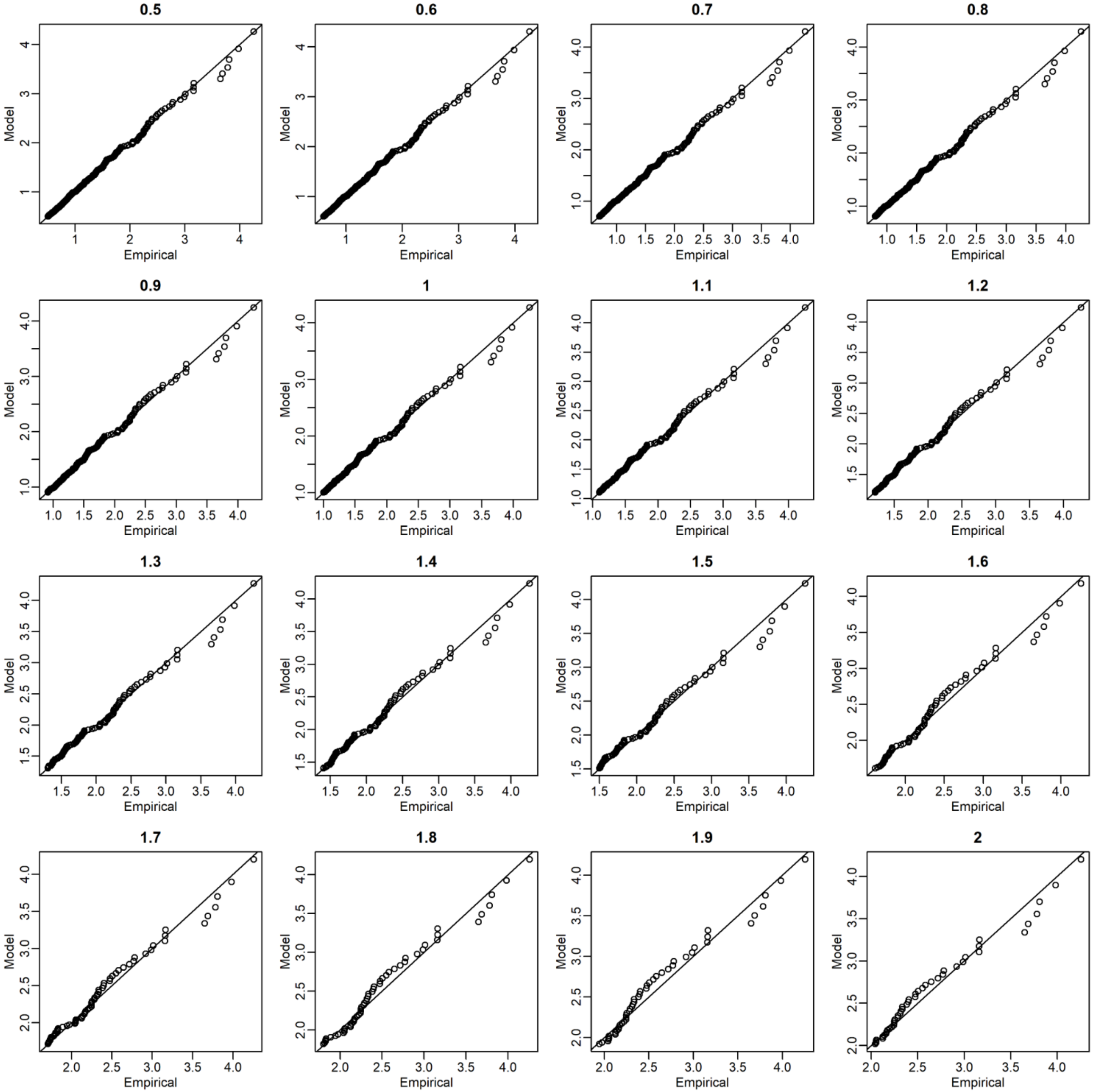


Figure S6 continued

(D)ESeAsia_Oc (Eastern and Southeastern Asia and Oceania)

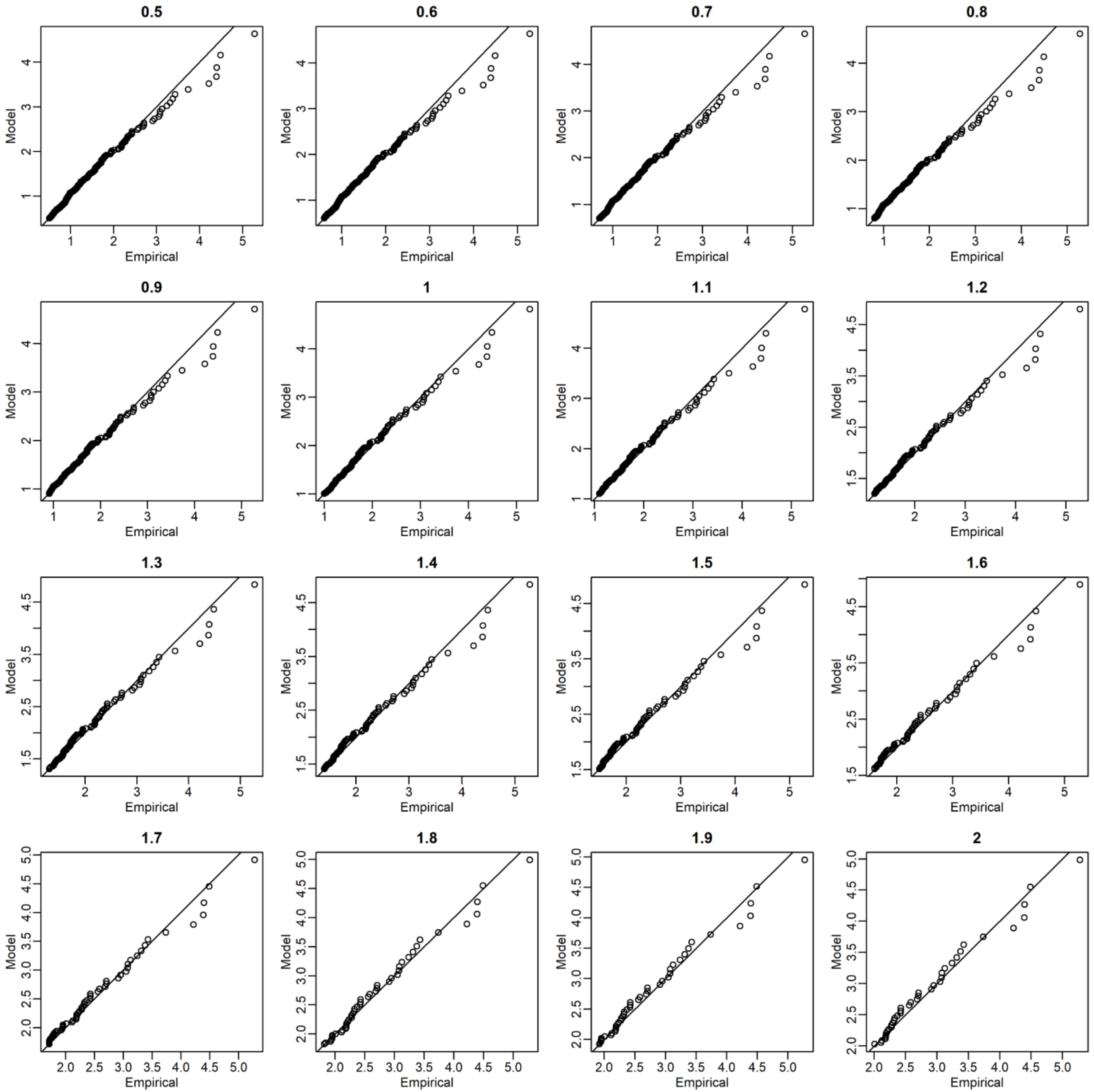


Figure S6 continued

(E) Europe

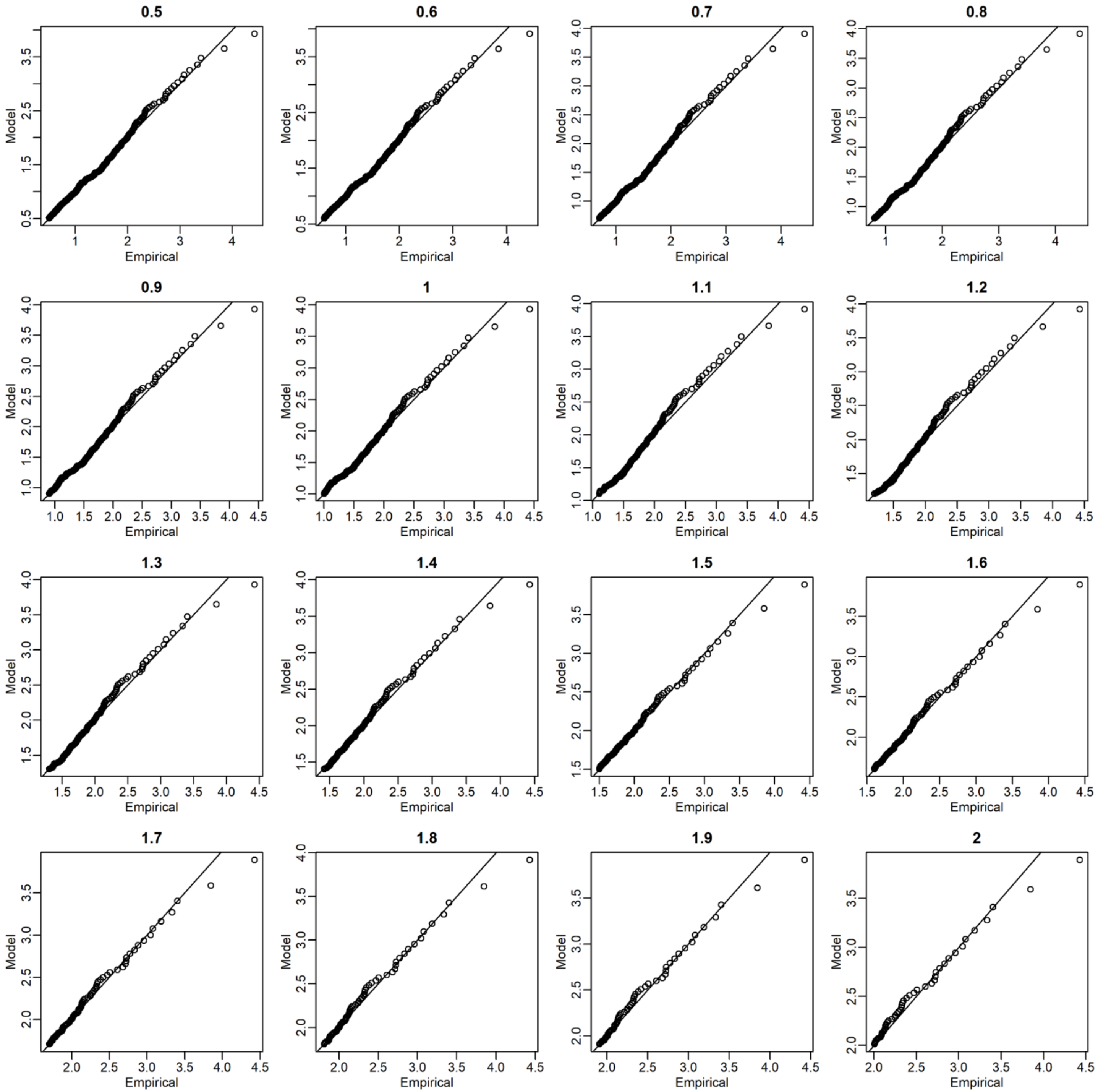


Figure S6 continued

(A) Africa

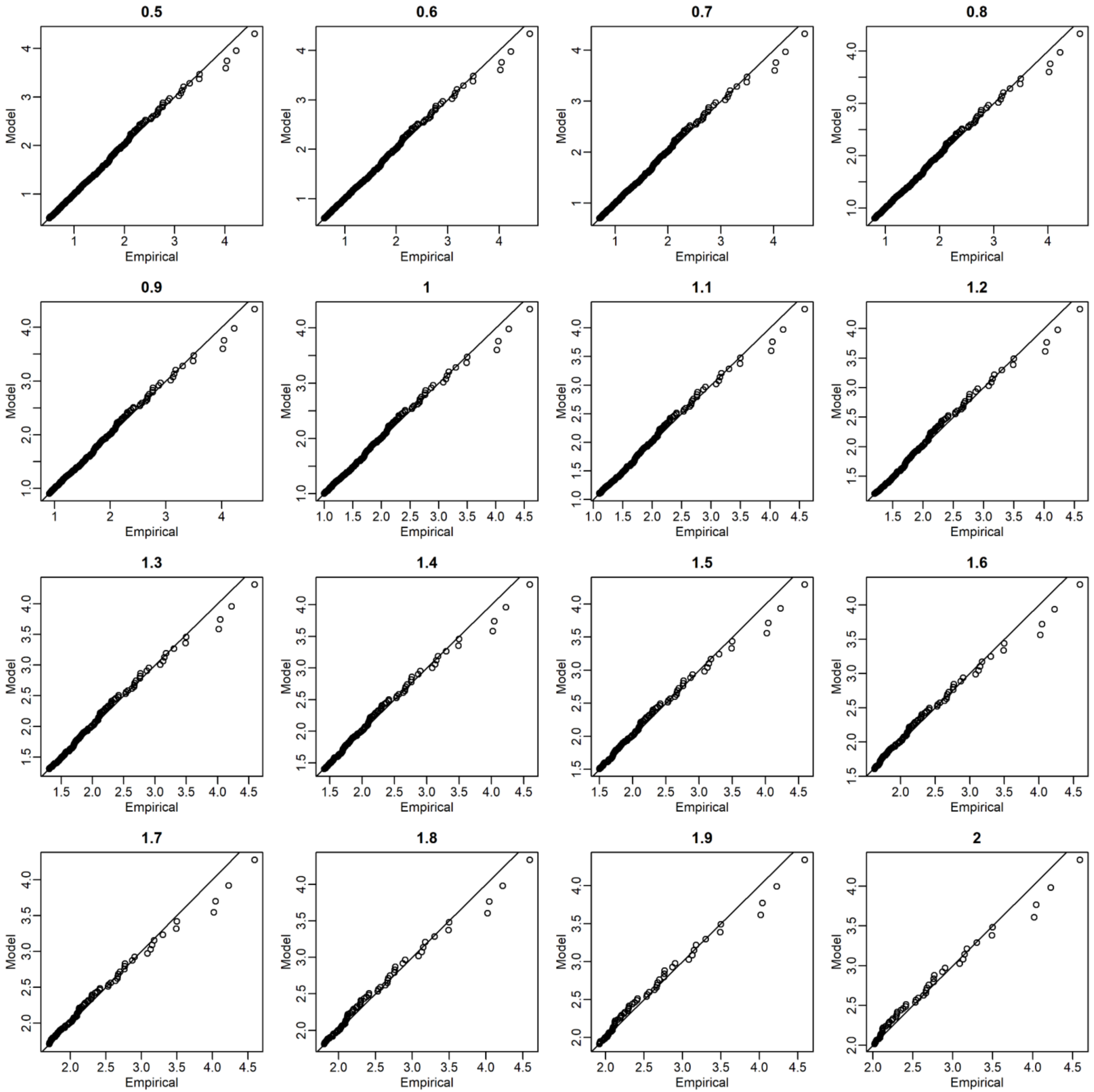


**Figure S7** Quantile-quantile (QQ) plots for wheat of the global yield data between the empirical (observed) and approximated distributions. The empirical distributions were approximated with the stationary generalized Pareto models. The titles of plots denote the threshold values ($T_h$).

### (B) Americas

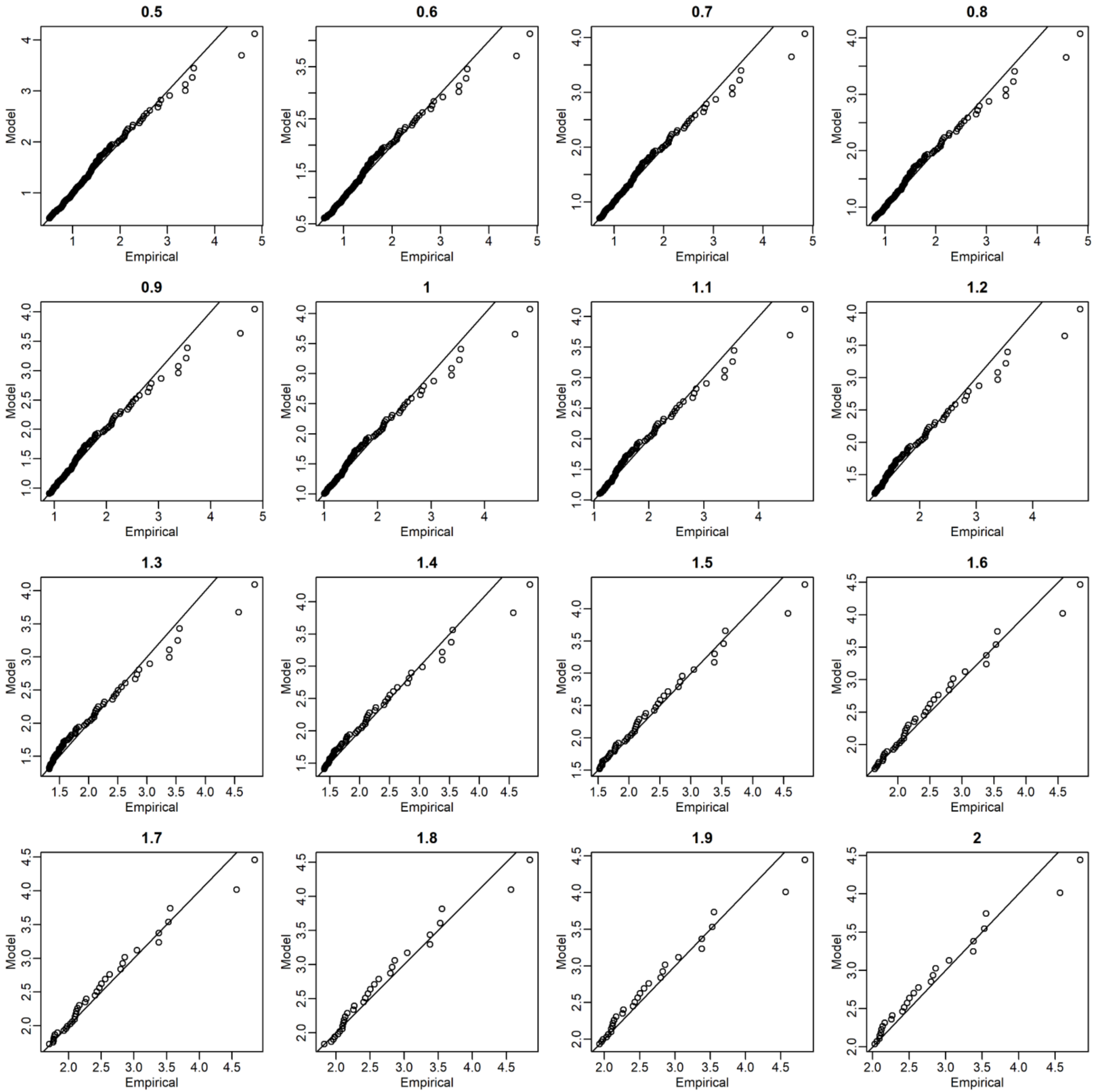


Figure S7 continued

(C) WCSAsia (Western, Central, and Southern Asia

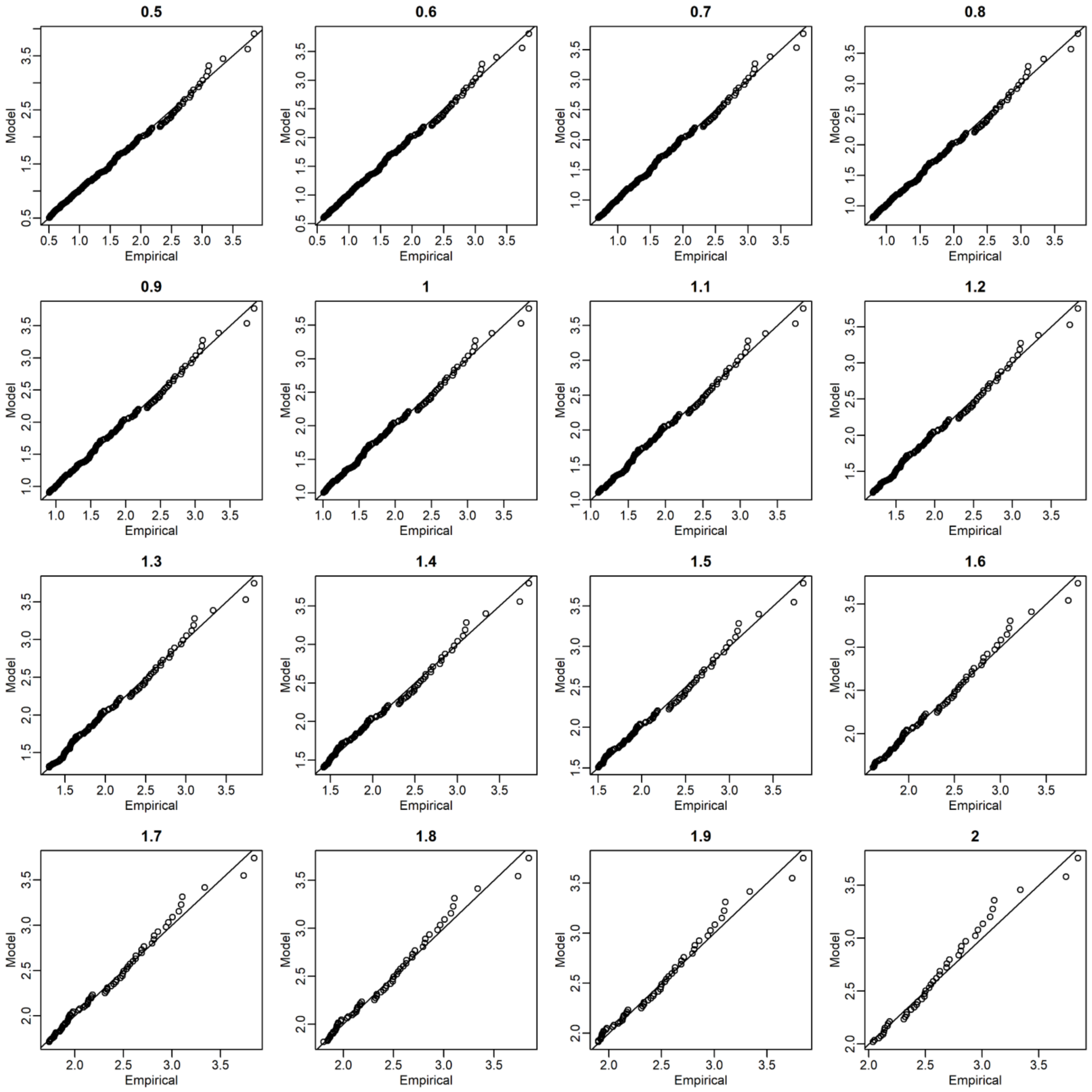


Figure S7 continued

(D)ESeAsia_Oc (Eastern and Southeastern Asia and Oceania)

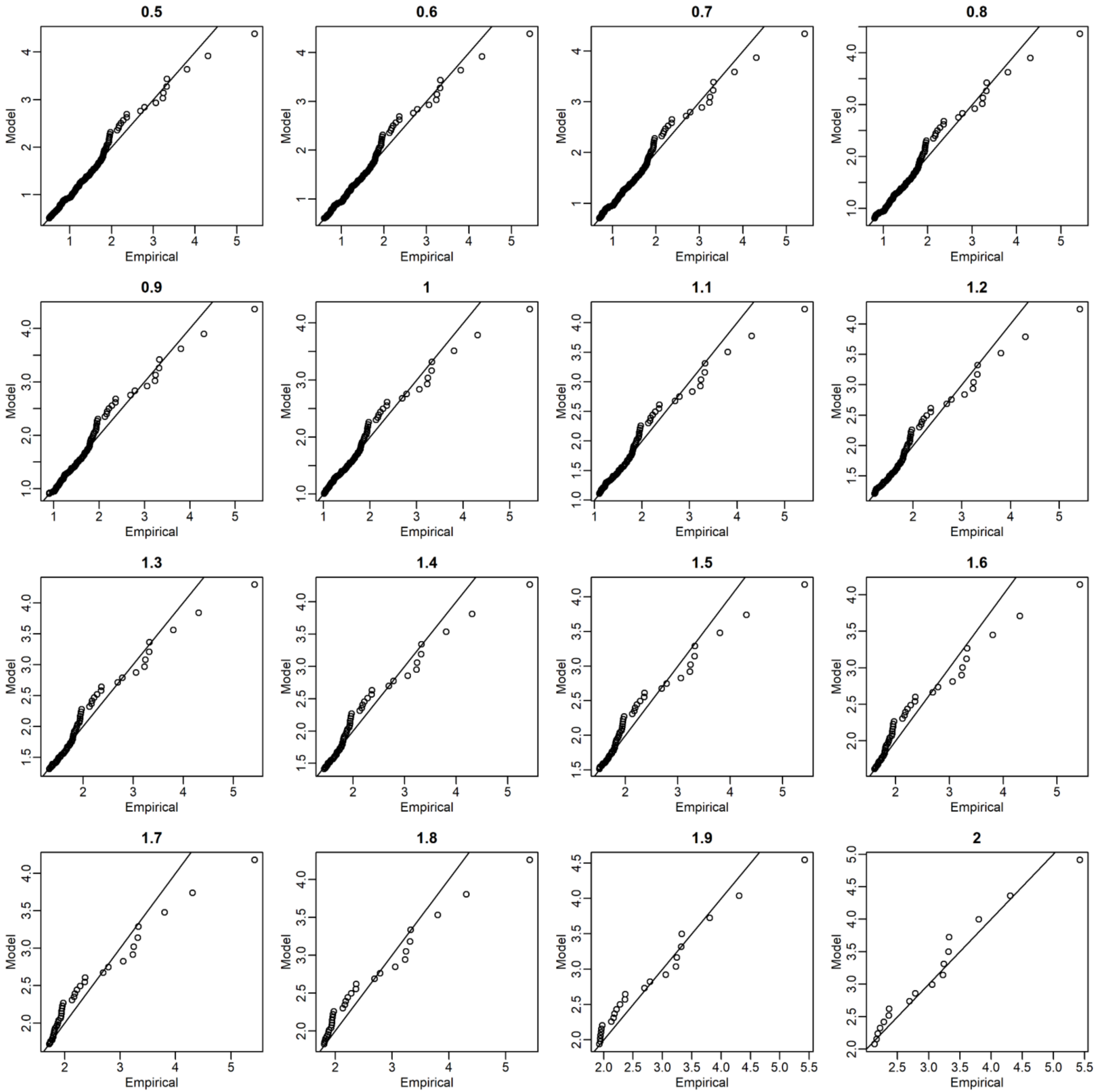


Figure S7 continued

(E) Europe

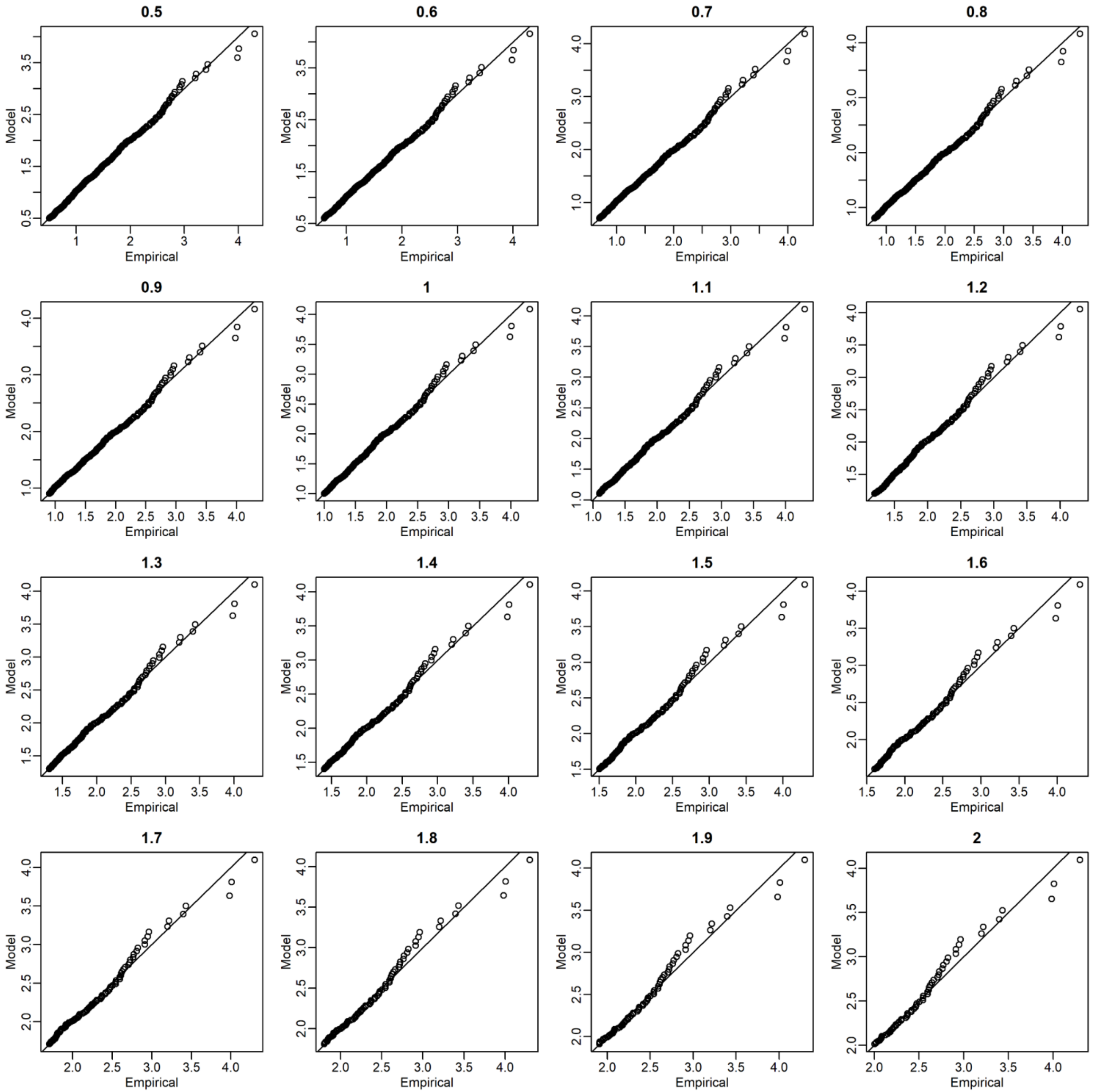


Figure S7 continued

(A) Africa

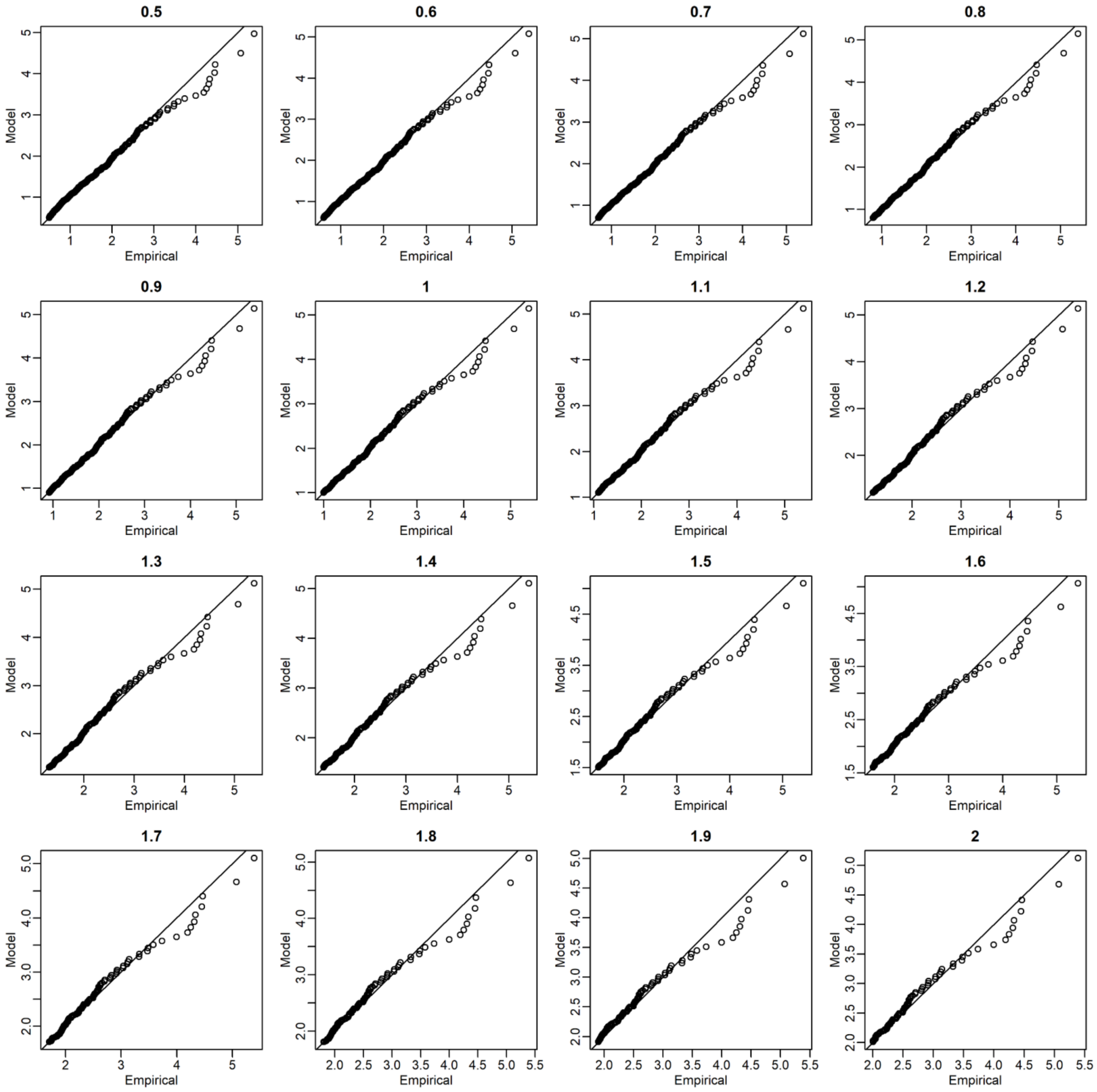


**Figure S8** Quantile-quantile (QQ) plots for rice of the global yield data between the empirical (observed) and approximated distributions. The empirical distributions were approximated with the stationary generalized Pareto models. The titles of plots denote the threshold values ($T_h$).

(B) Americas

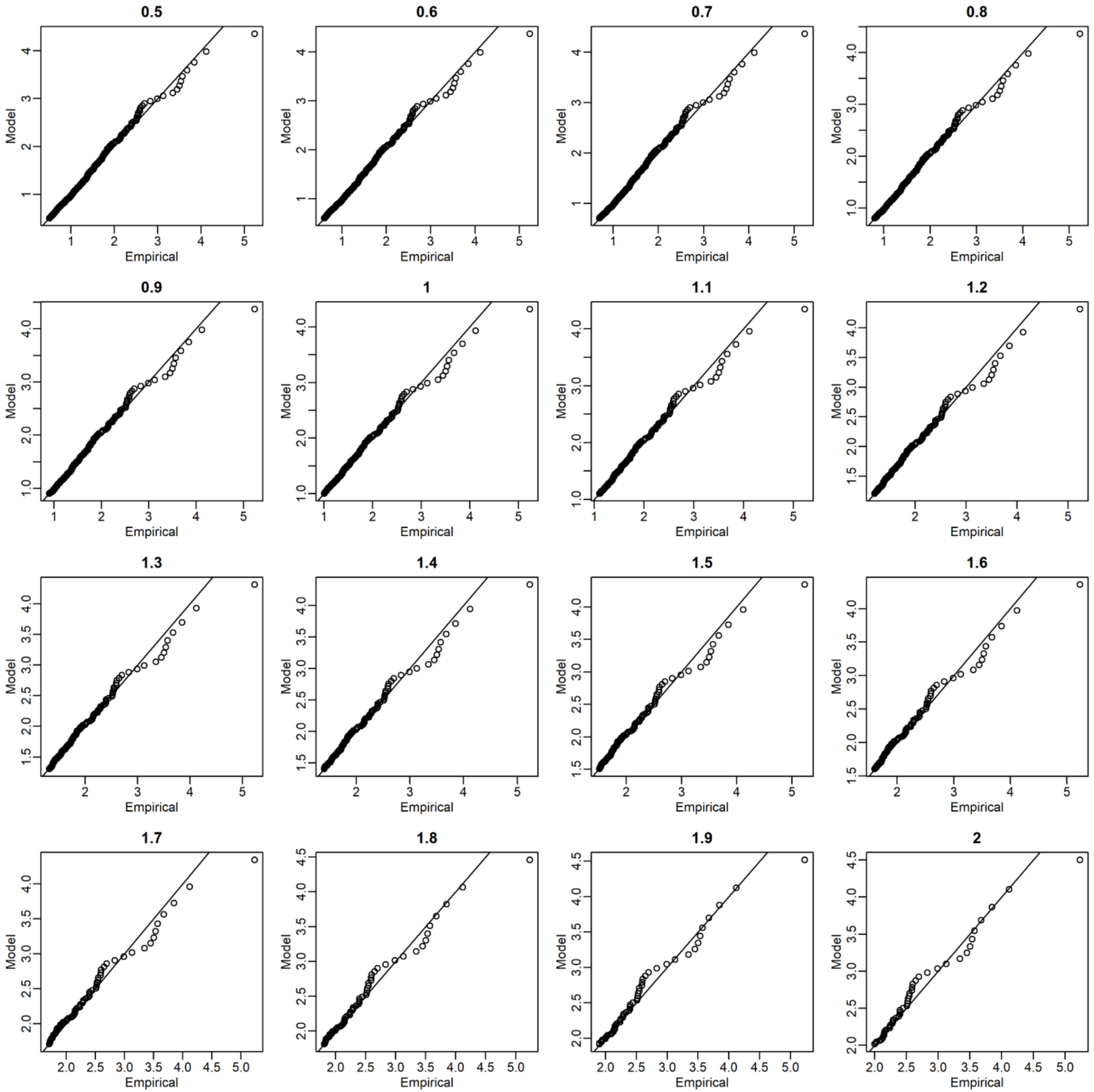


Figure S8 continued

(C) WCSAsia (Western, Central, and Southern Asia

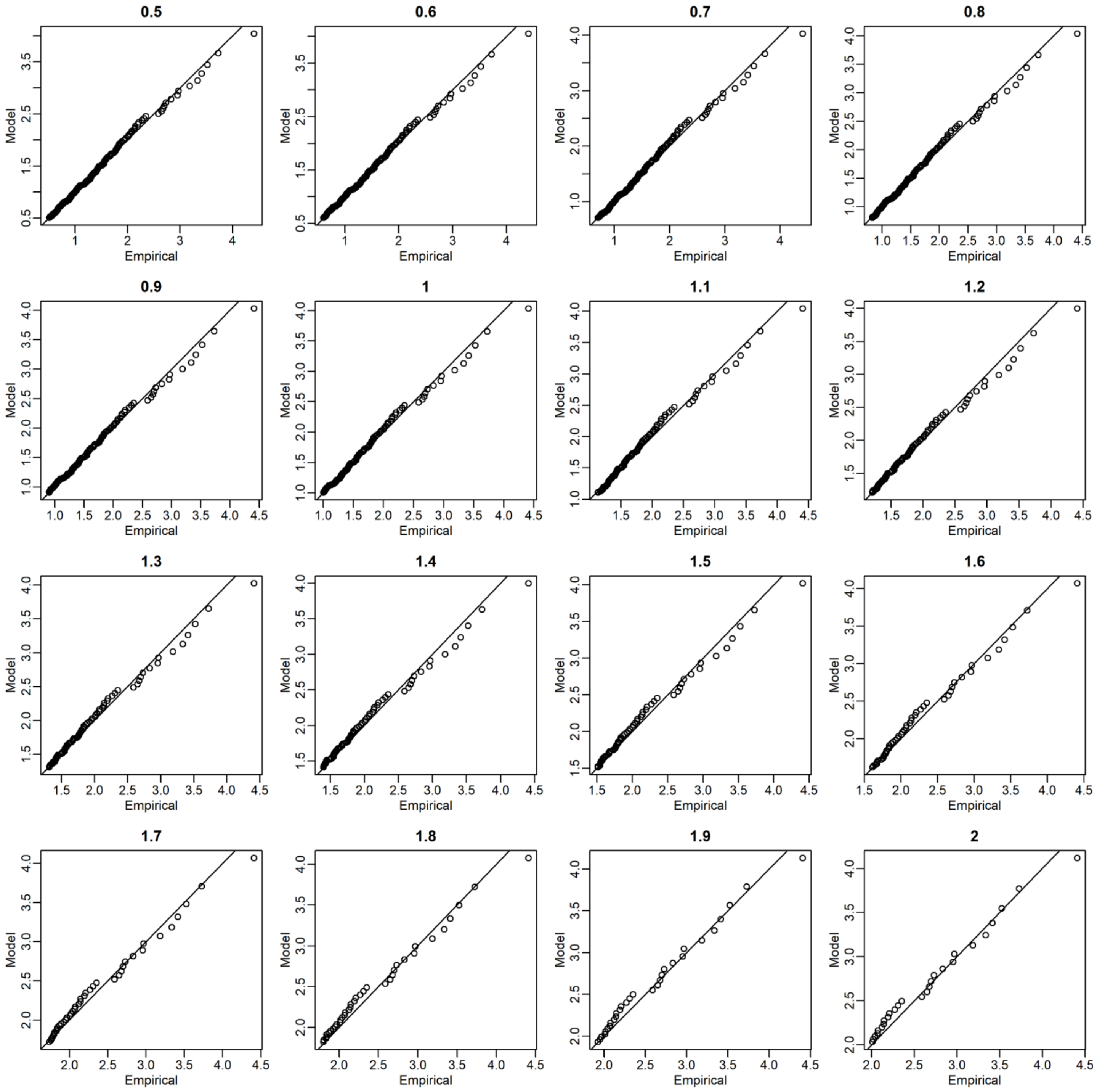


Figure S8 continued

(D)ESeAsia_Oc (Eastern and Southeastern Asia and Oceania)

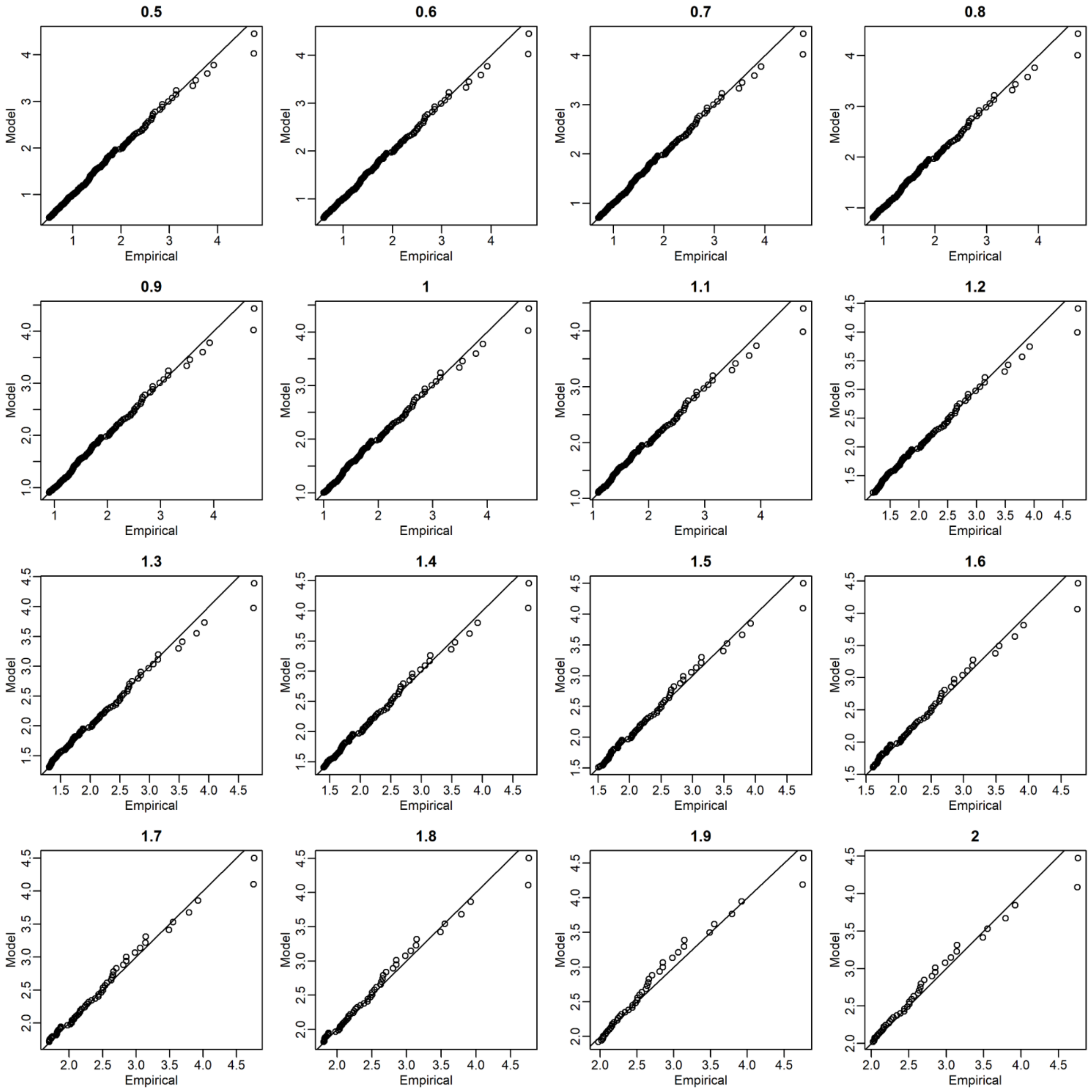


Figure S8 continued

(E) Europe

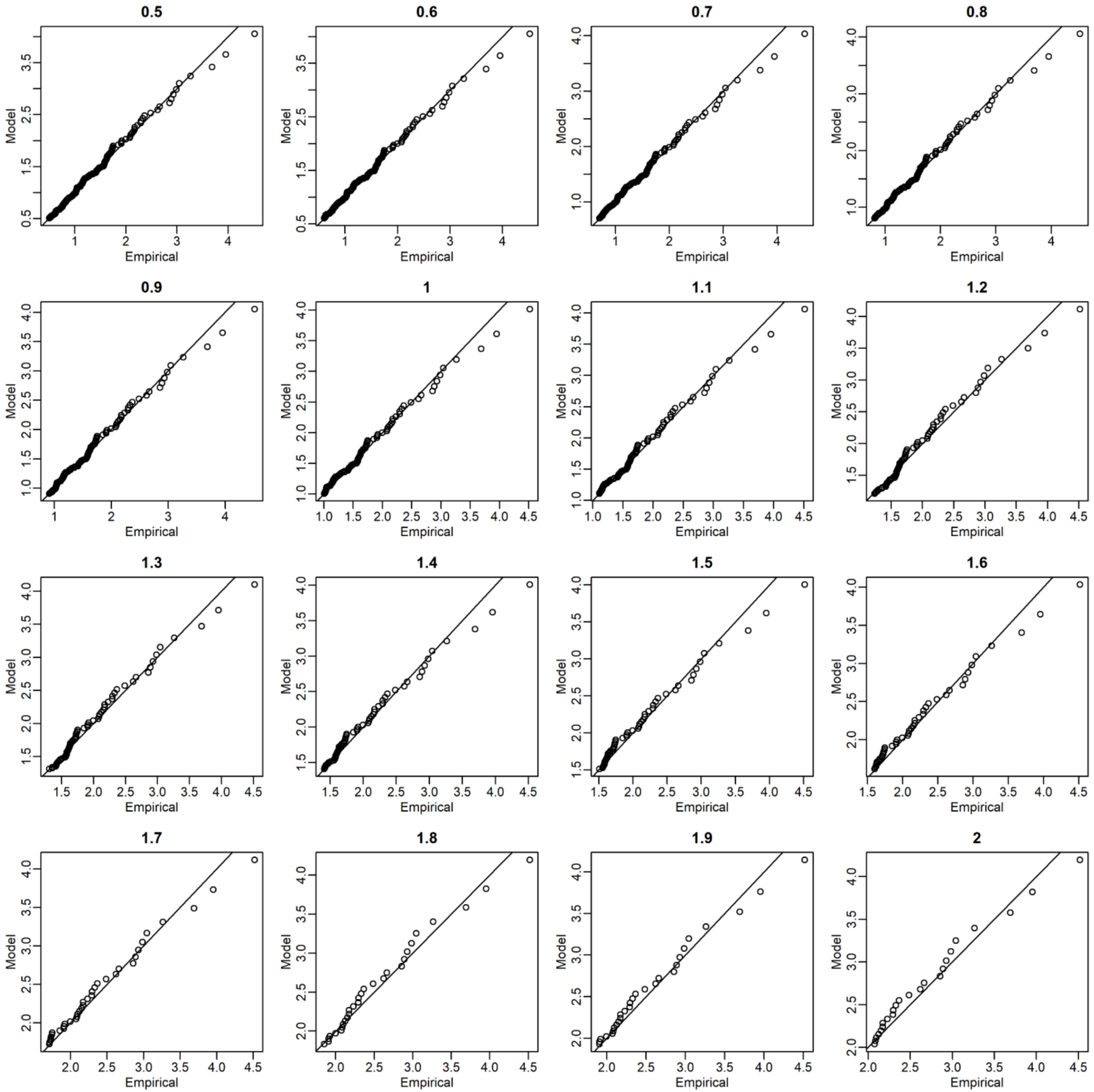


Figure S8 continued

(A) Africa

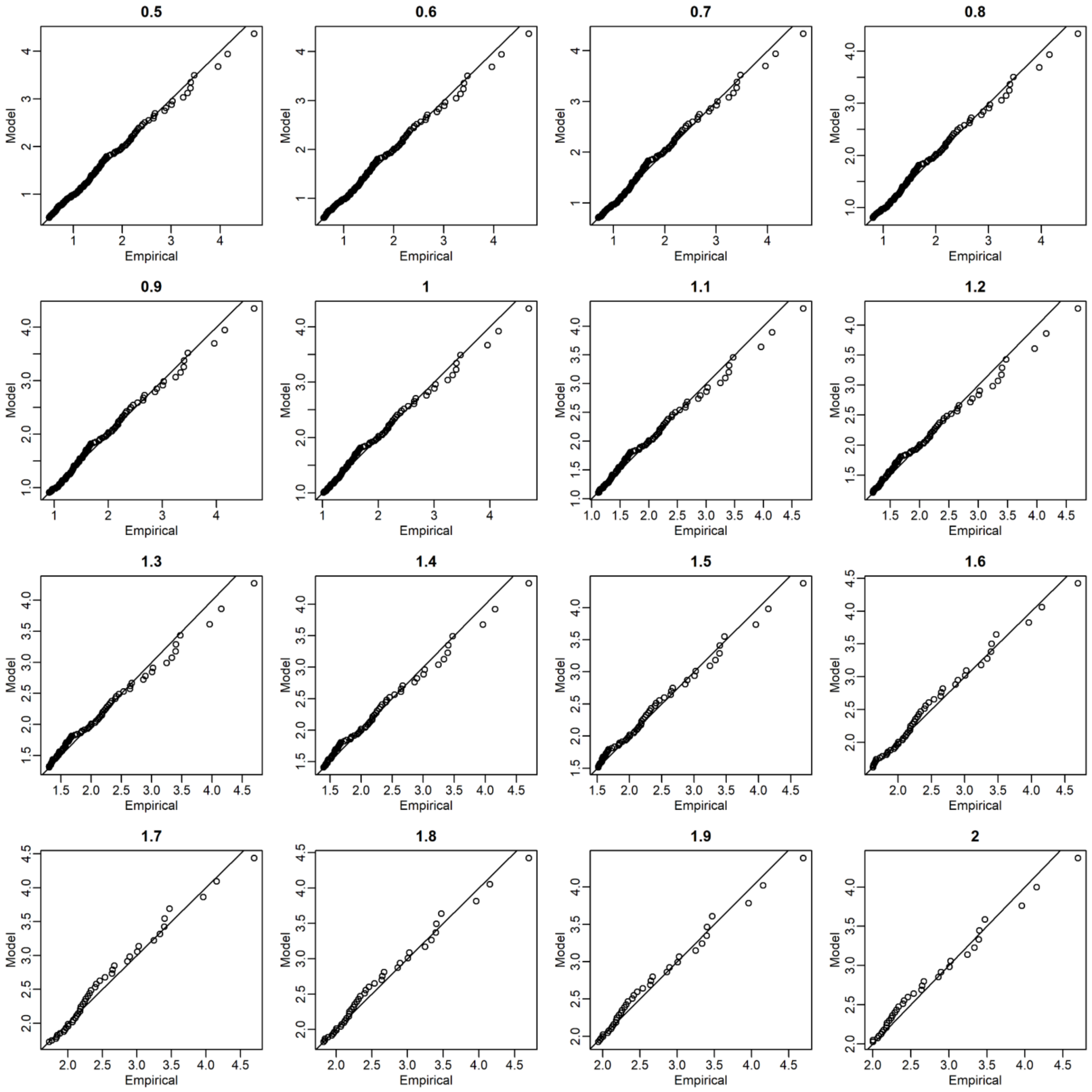


**Figure S9** Quantile-quantile (QQ) plots for soybean of the global yield data between the empirical (observed) and approximated distributions. The empirical distributions were approximated with the stationary generalized Pareto models. The titles of plots denote the threshold values ($T_h$).

### (B) Americas

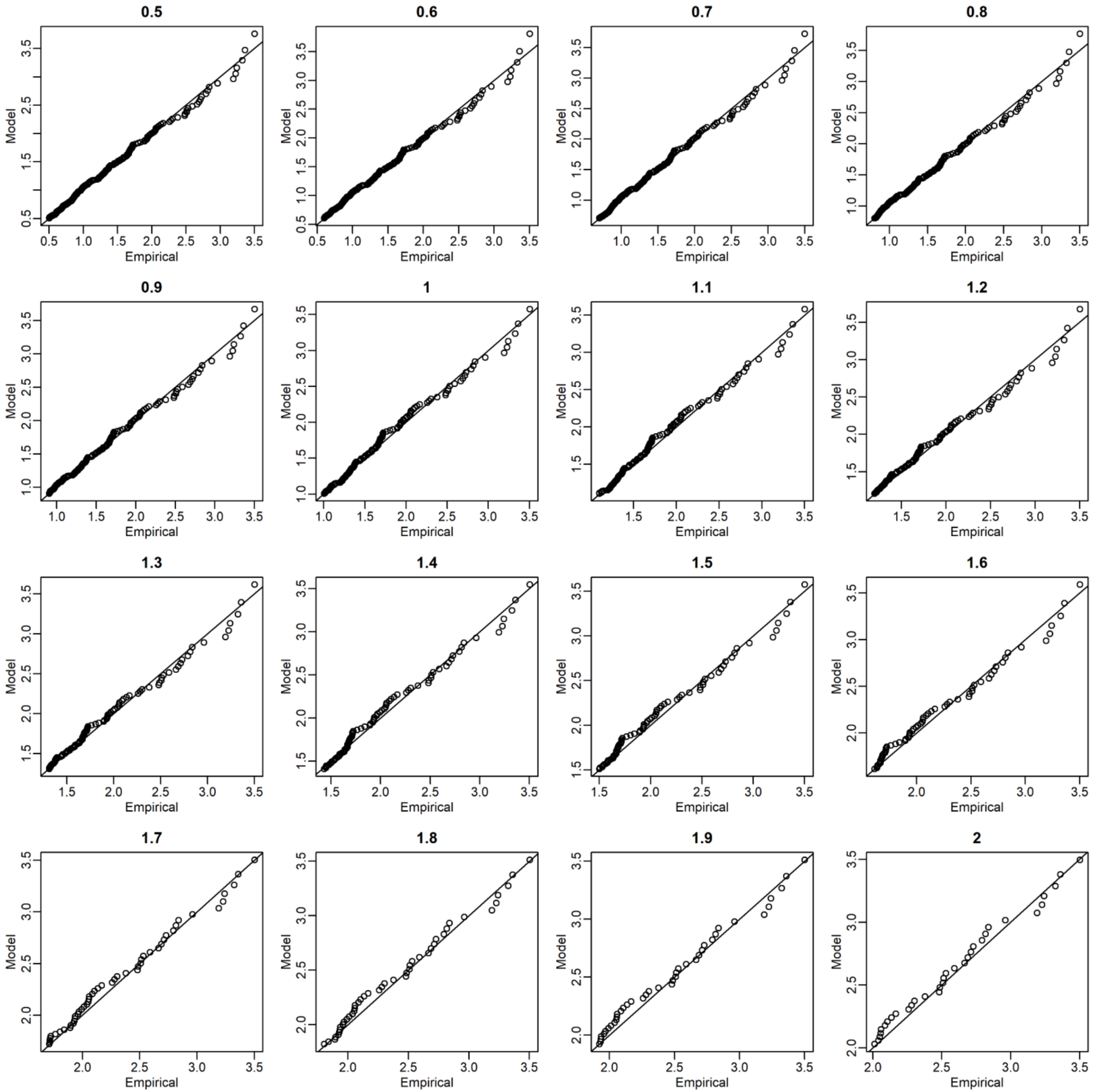


Figure S9 continued

(C) WCSAsia
(Western, Central,
and Southern Asia

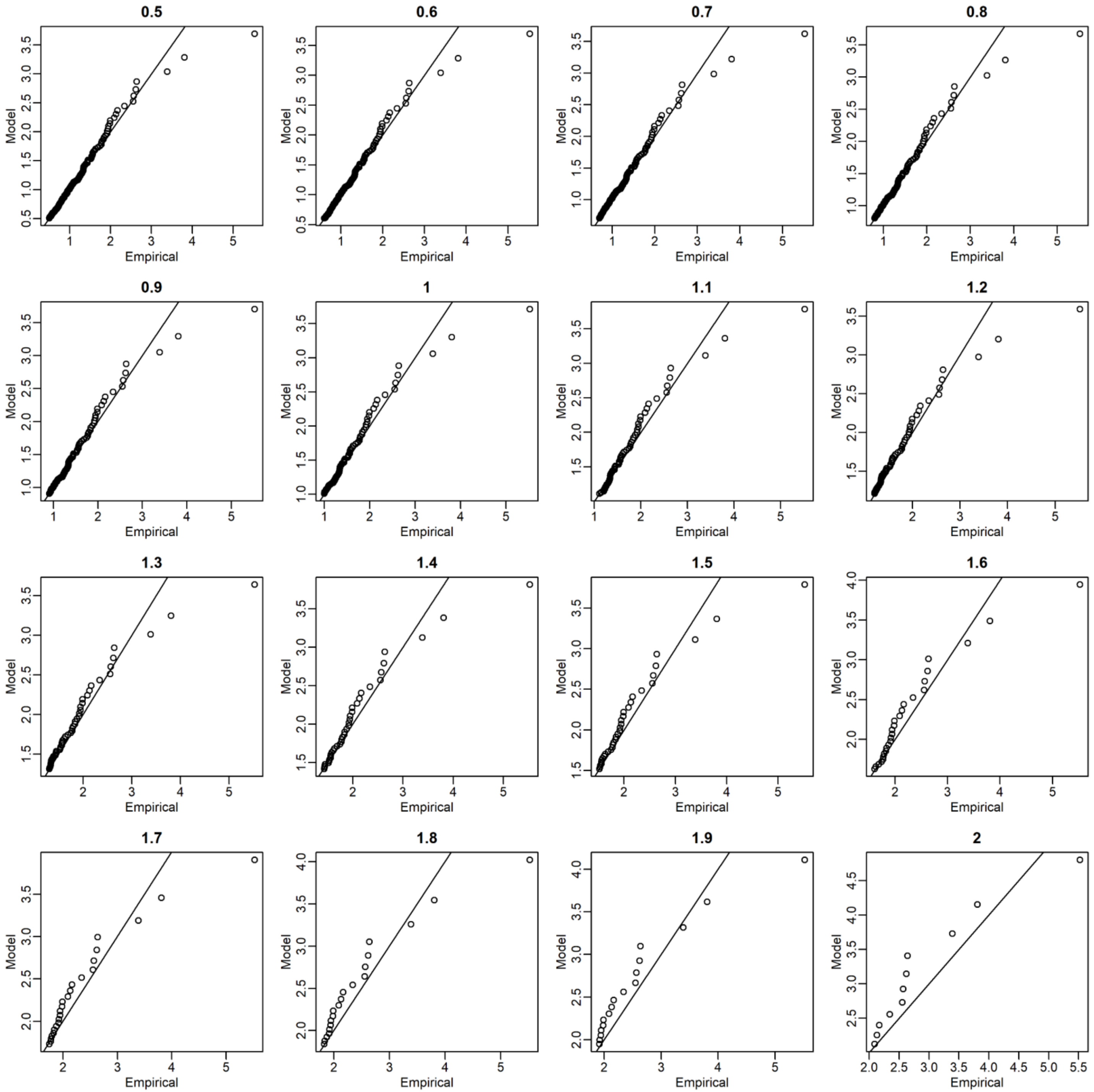


Figure S9 continued

(D)ESeAsia_Oc (Eastern and Southeastern Asia and Oceania)

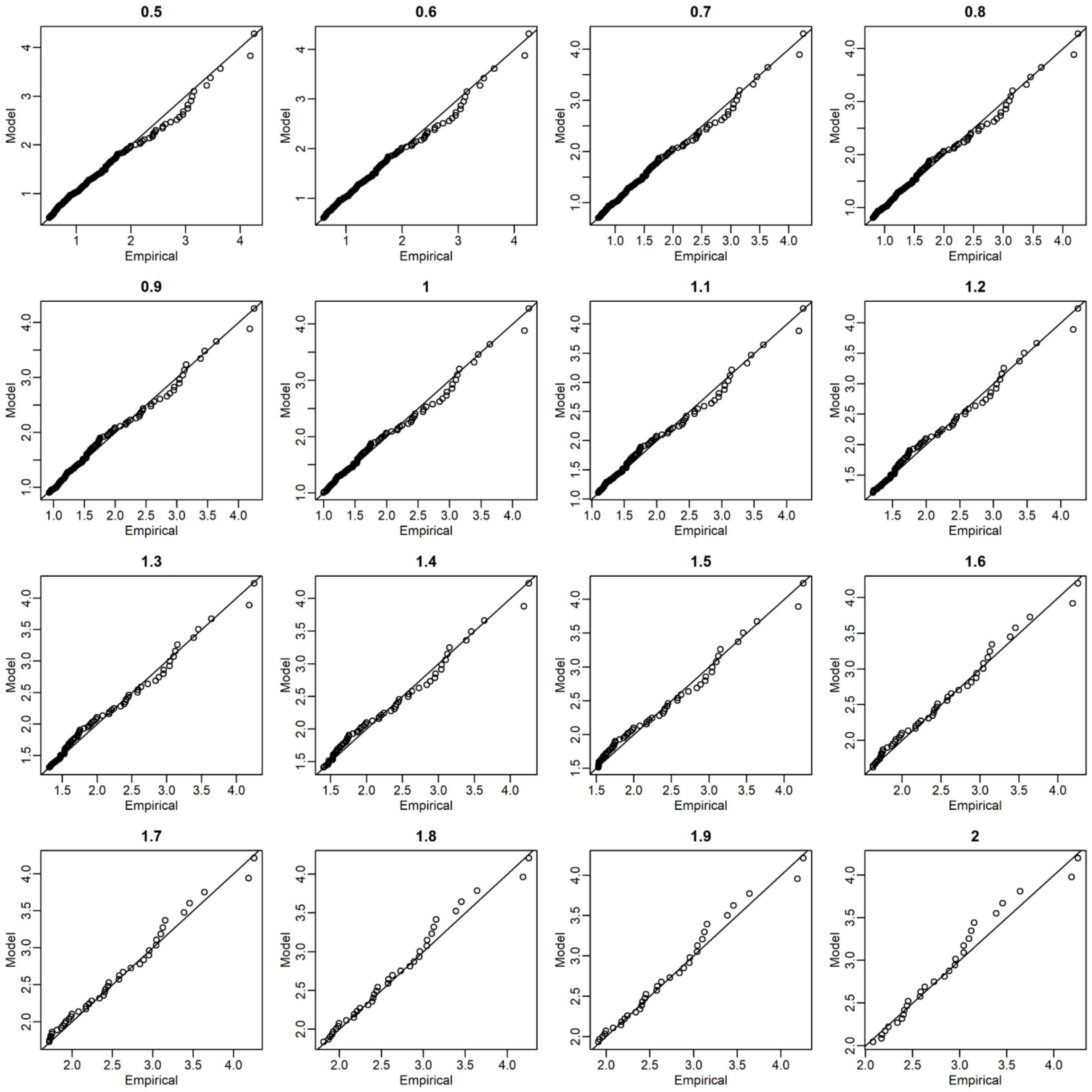


Figure S9 continued

(E) Europe

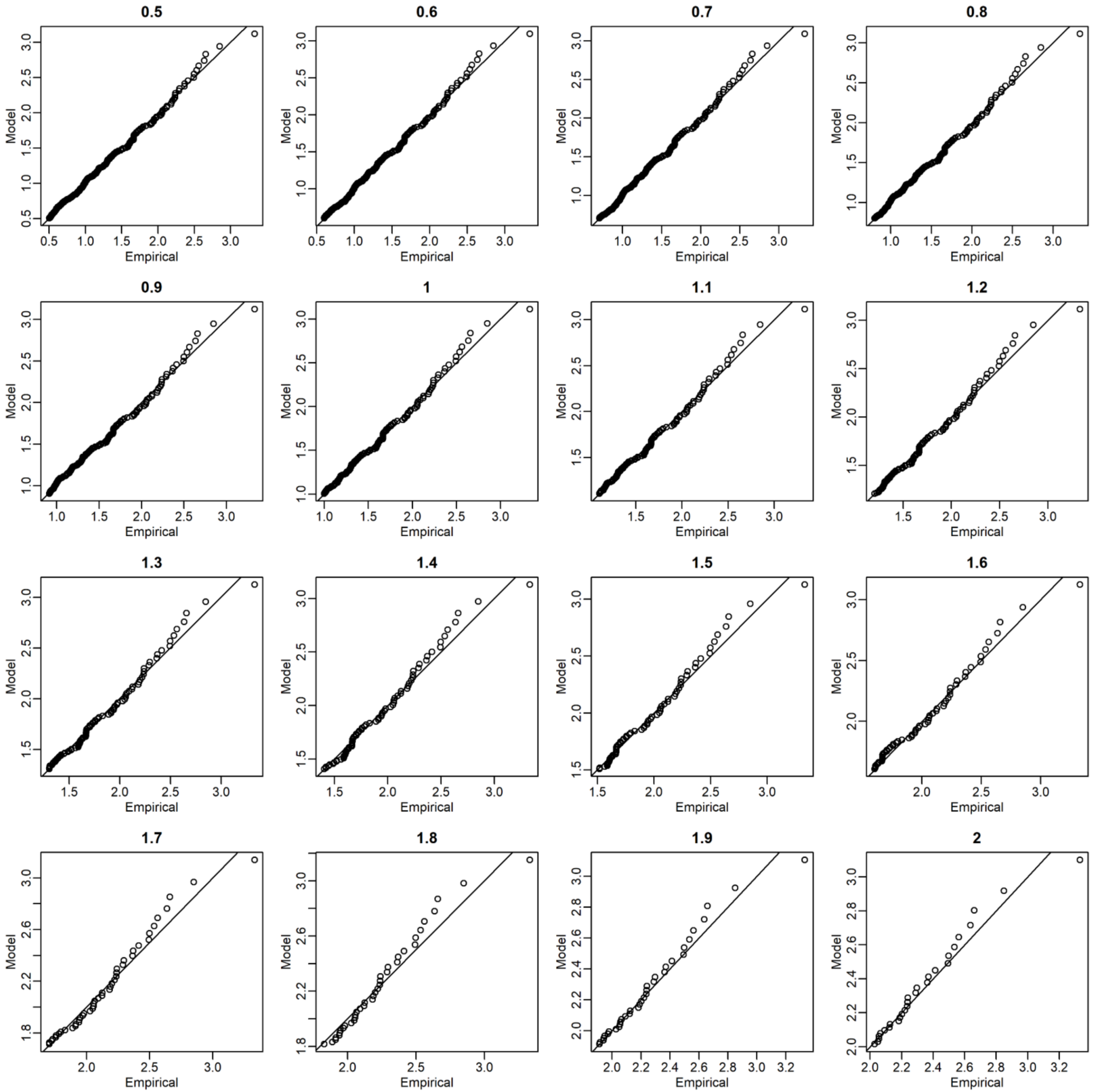


Figure S9 continued

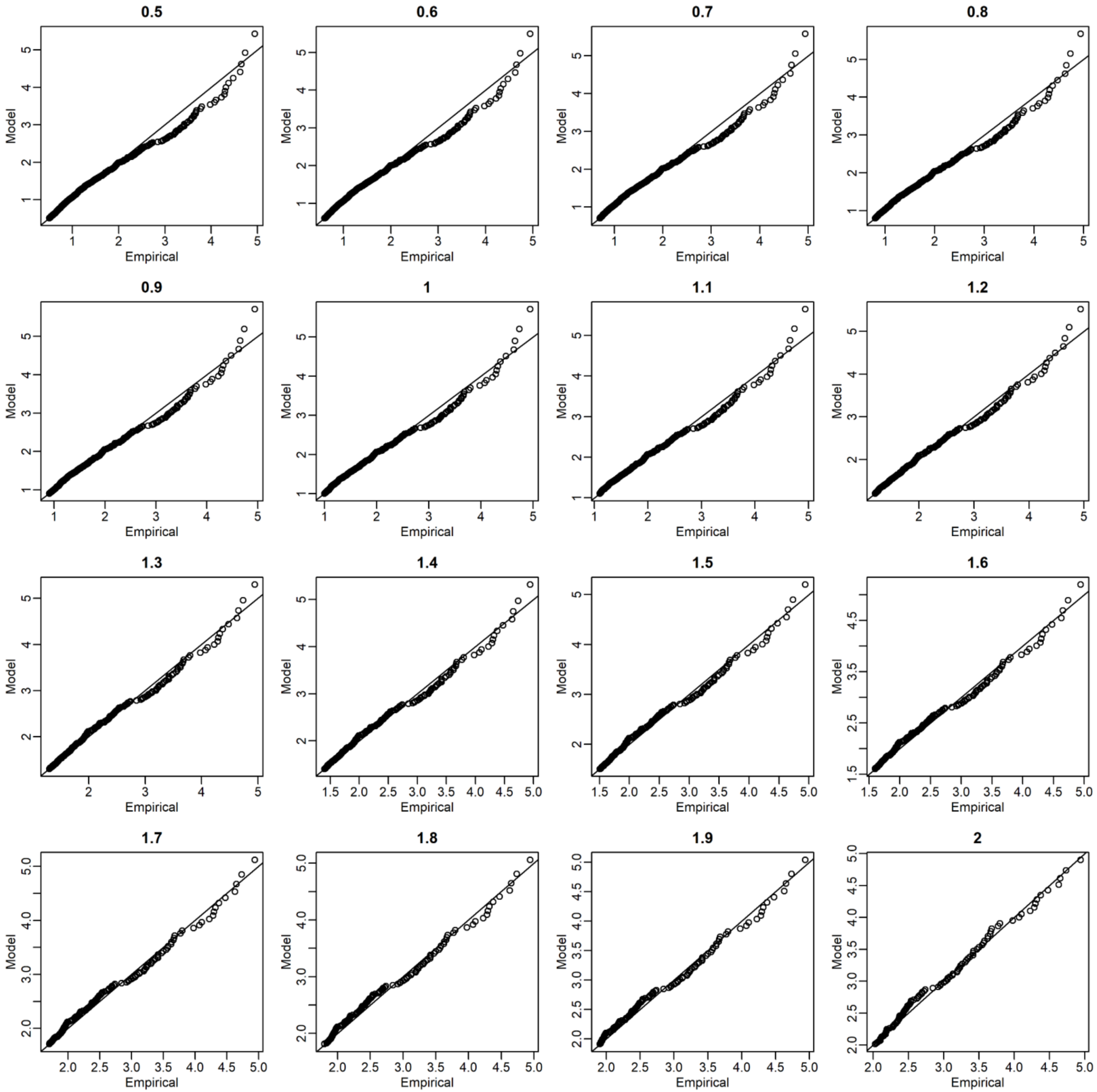


**Figure S10** Quantile-quantile (QQ) plots for wheat of the local yield data between the empirical (observed) and approximated distributions. The empirical distributions were approximated with the stationary generalized Pareto models. The titles of plots denote the threshold values ($T_h$).

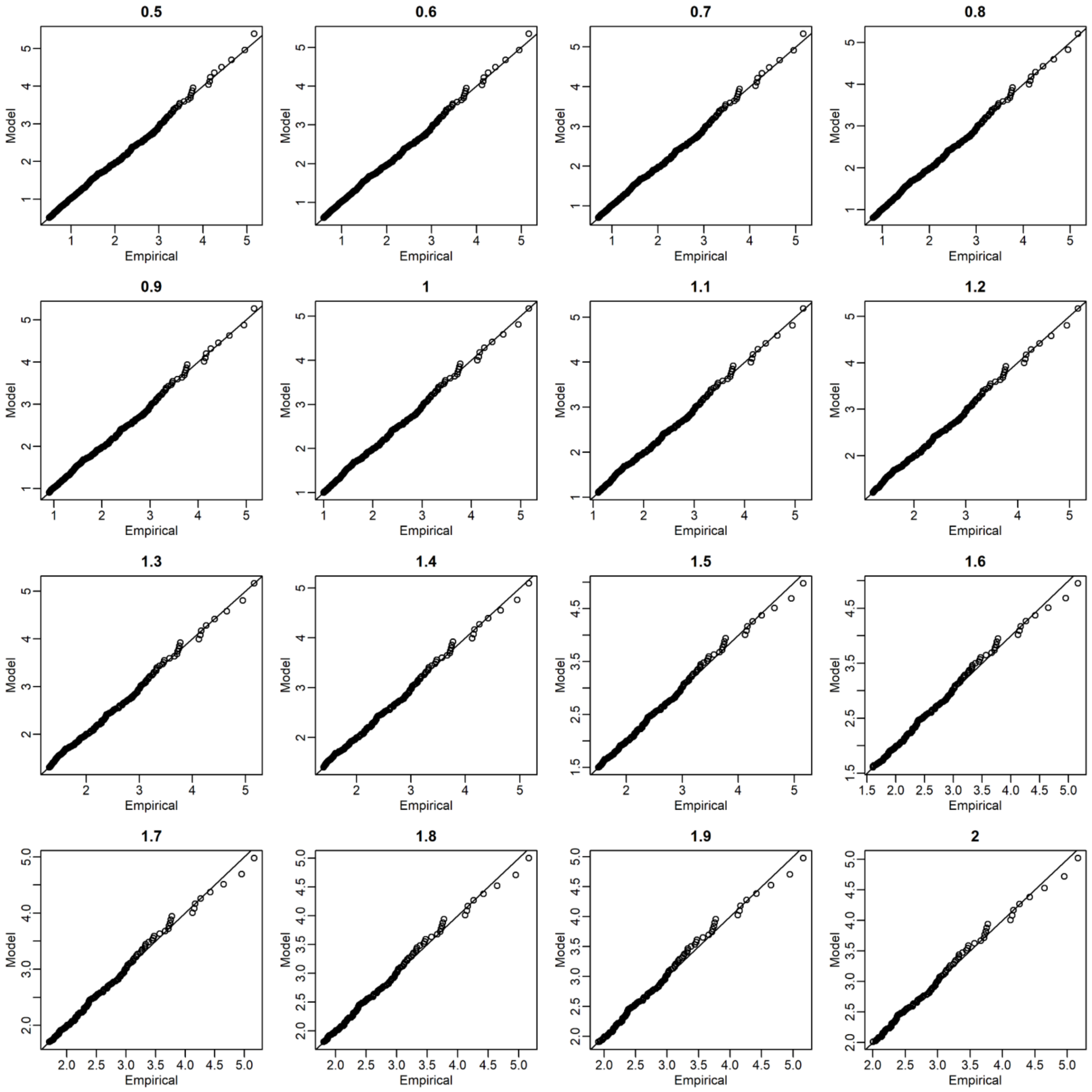


**Figure S11** Quantile-quantile (QQ) plots for rice of the local yield data between the empirical (observed) and approximated distributions. The empirical distributions were approximated with the stationary generalized Pareto models. The titles of plots denote the threshold values ($T_h$).

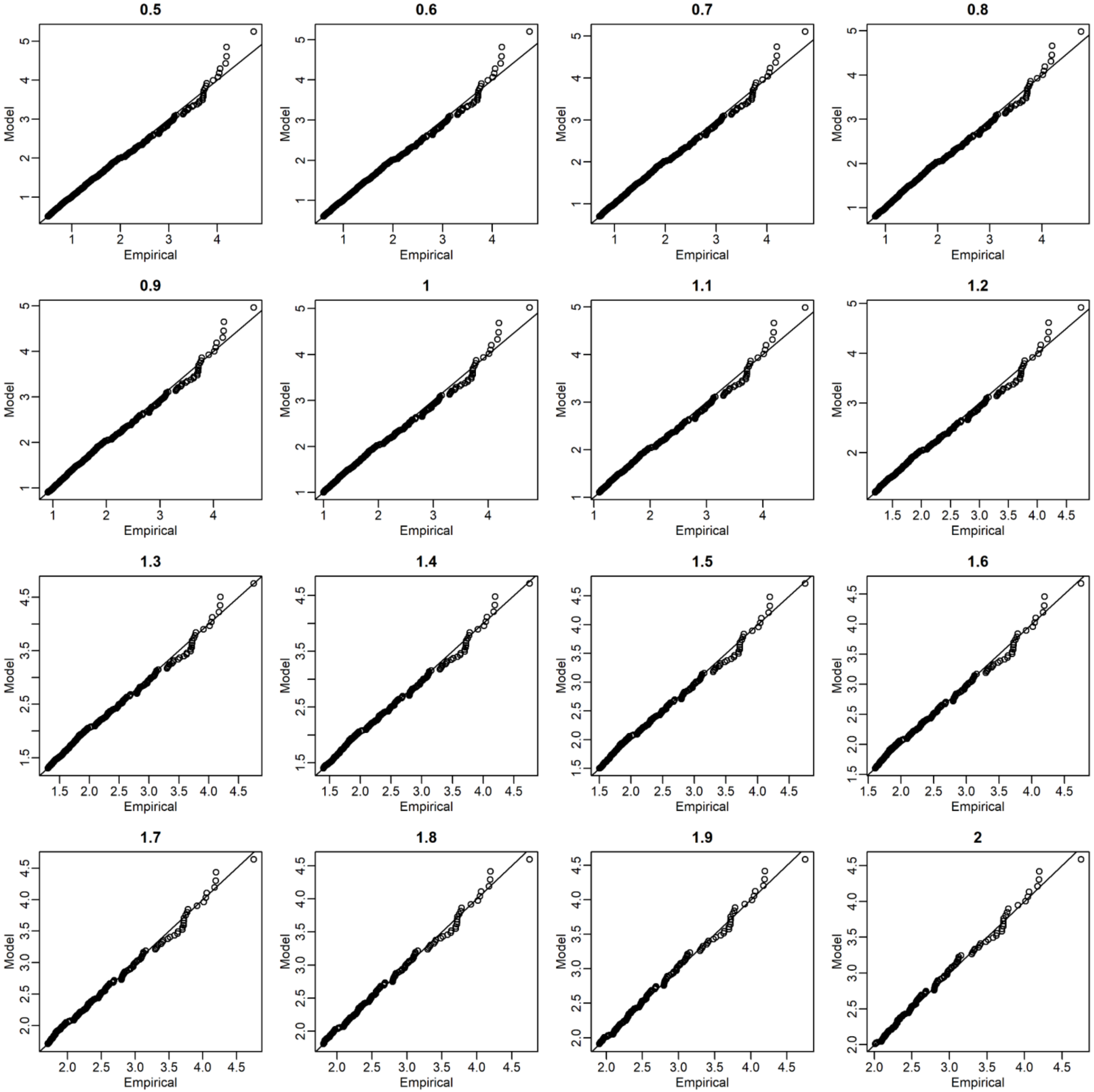


**Figure S12** Quantile-quantile (QQ) plots for soybean of the local yield data between the empirical (observed) and approximated distributions. The empirical distributions were approximated with the stationary generalized Pareto models. The titles of plots denote the threshold values ($T_h$).

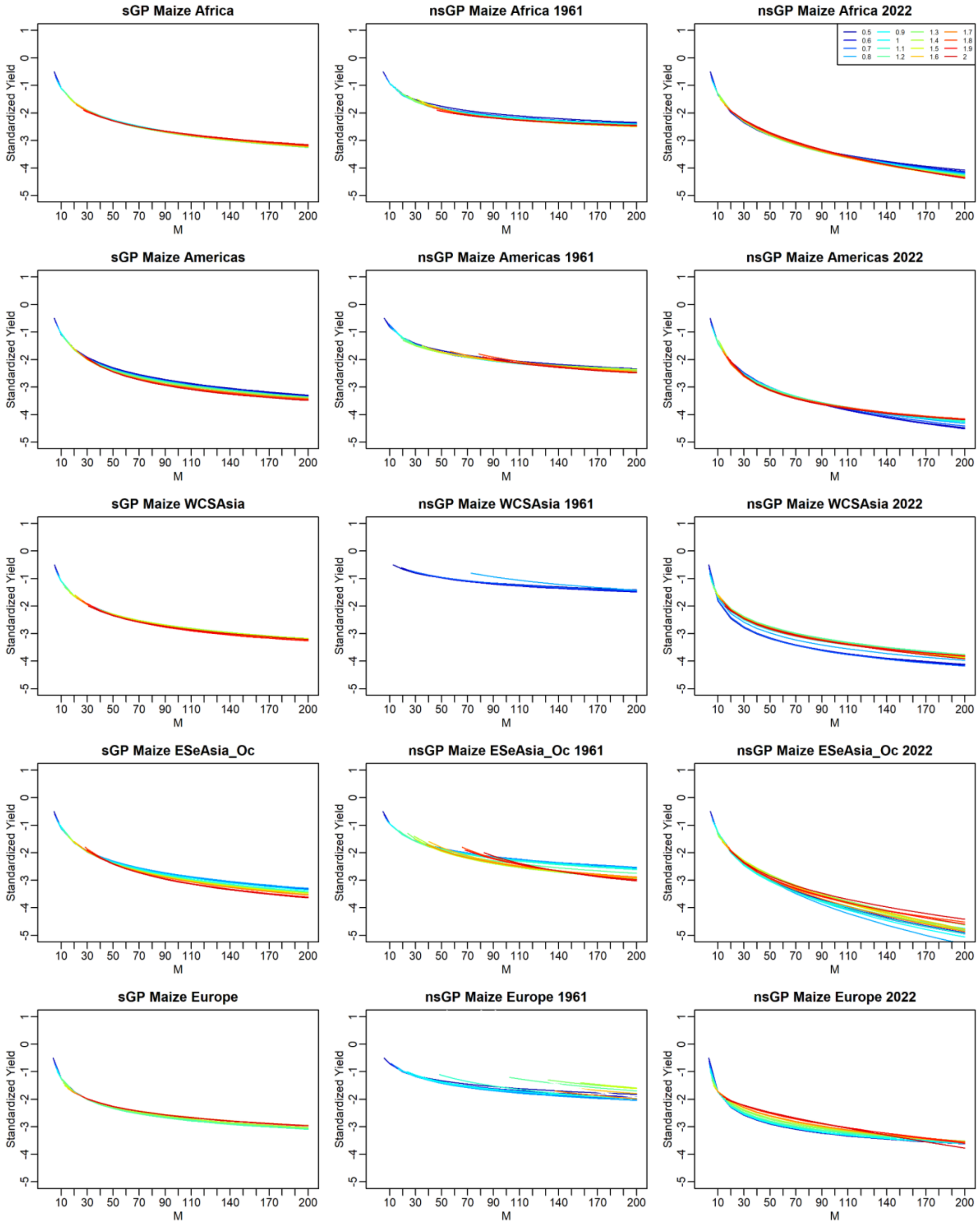


**Figure S13** *M*-year return levels estimated for maize in the global yield data. The left panels are the levels estimated with the stationary generalized Pareto (sGP) models and the middle and right panels were those estimated with the non-stationary GP (nsGP) models. For the nsGP models, the levels estimated at the oldest (1961) and latest (2022) years are presented. Different colors indicate the different threshold ($T_h$) values.

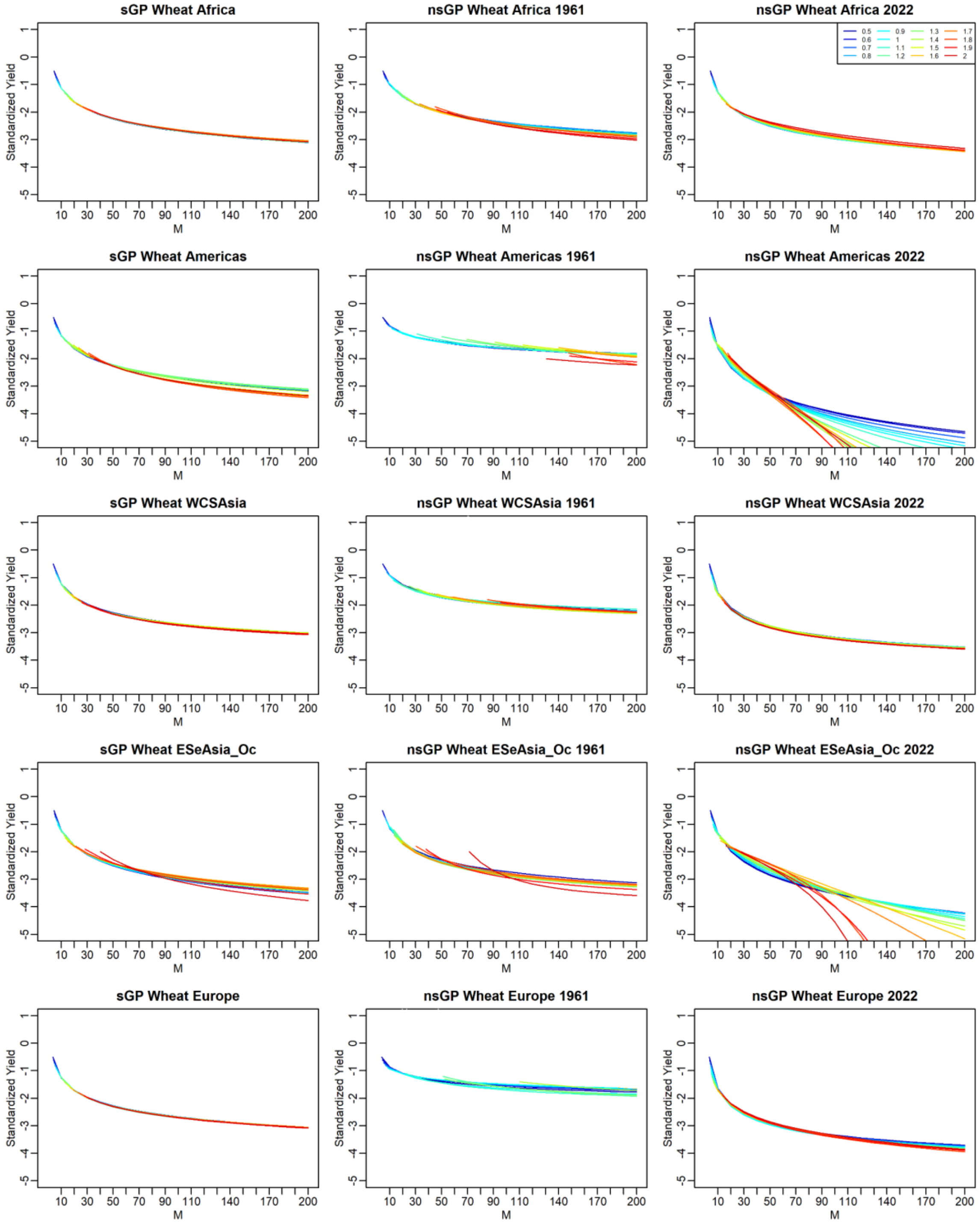


**Figure S14** *M*-year return levels estimated for wheat in the global yield data. The left panels are the levels estimated with the stationary generalized Pareto (sGP) models and the middle and right panels were those estimated with the non-stationary GP (nsGP) models. For the nsGP models, the levels estimated at the oldest (1961) and latest (2022) years are presented. Different colors indicate the different threshold ($T_h$) values.

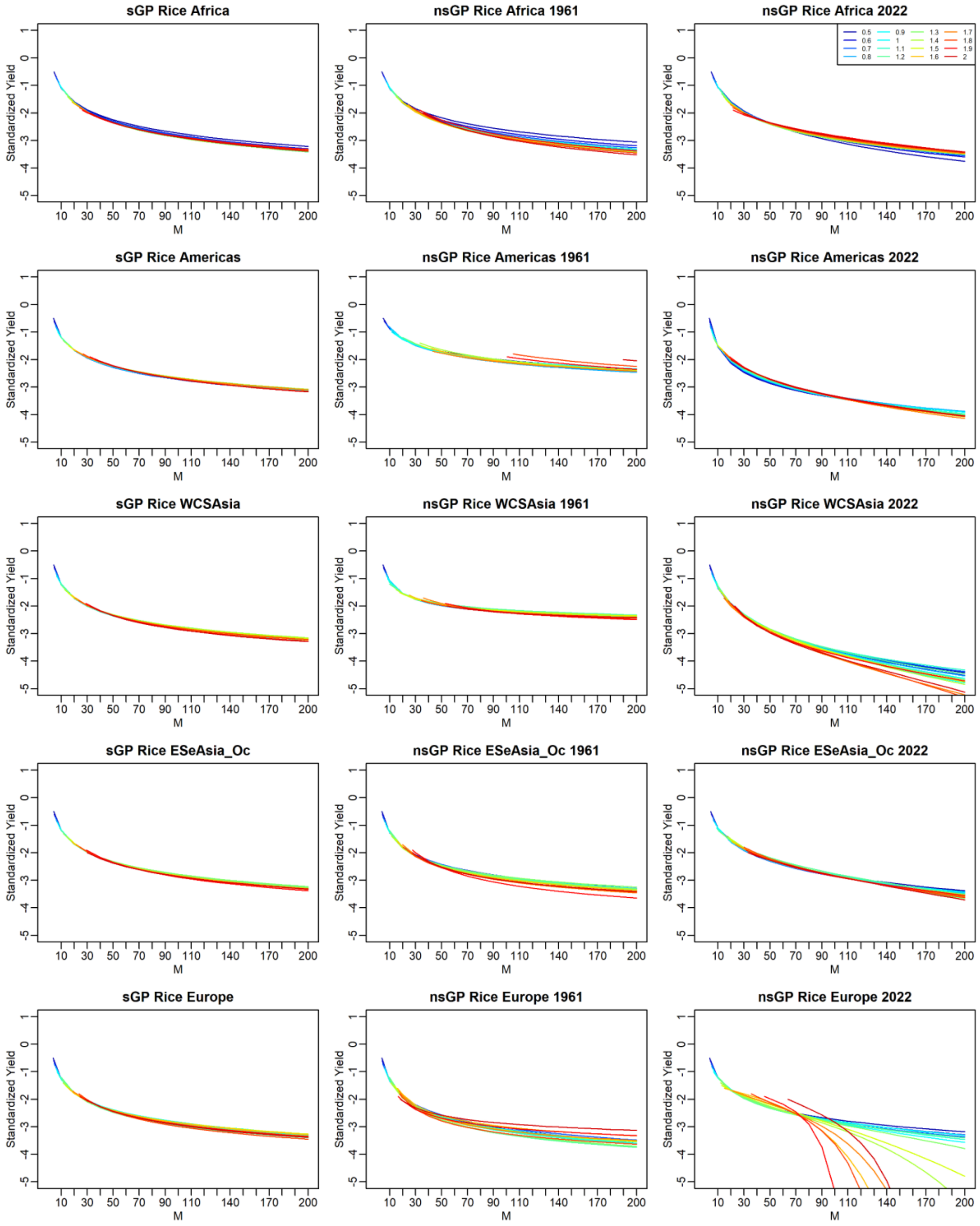


**Figure S15** *M*-year return levels estimated for rice in the global yield data. The left panels are the levels estimated with the stationary generalized Pareto (sGP) models and the middle and right panels were those estimated with the non-stationary GP (nsGP) models. For the nsGP models, the levels estimated at the oldest (1961) and latest (2022) years are presented. Different colors indicate the different threshold ($T_h$) values.

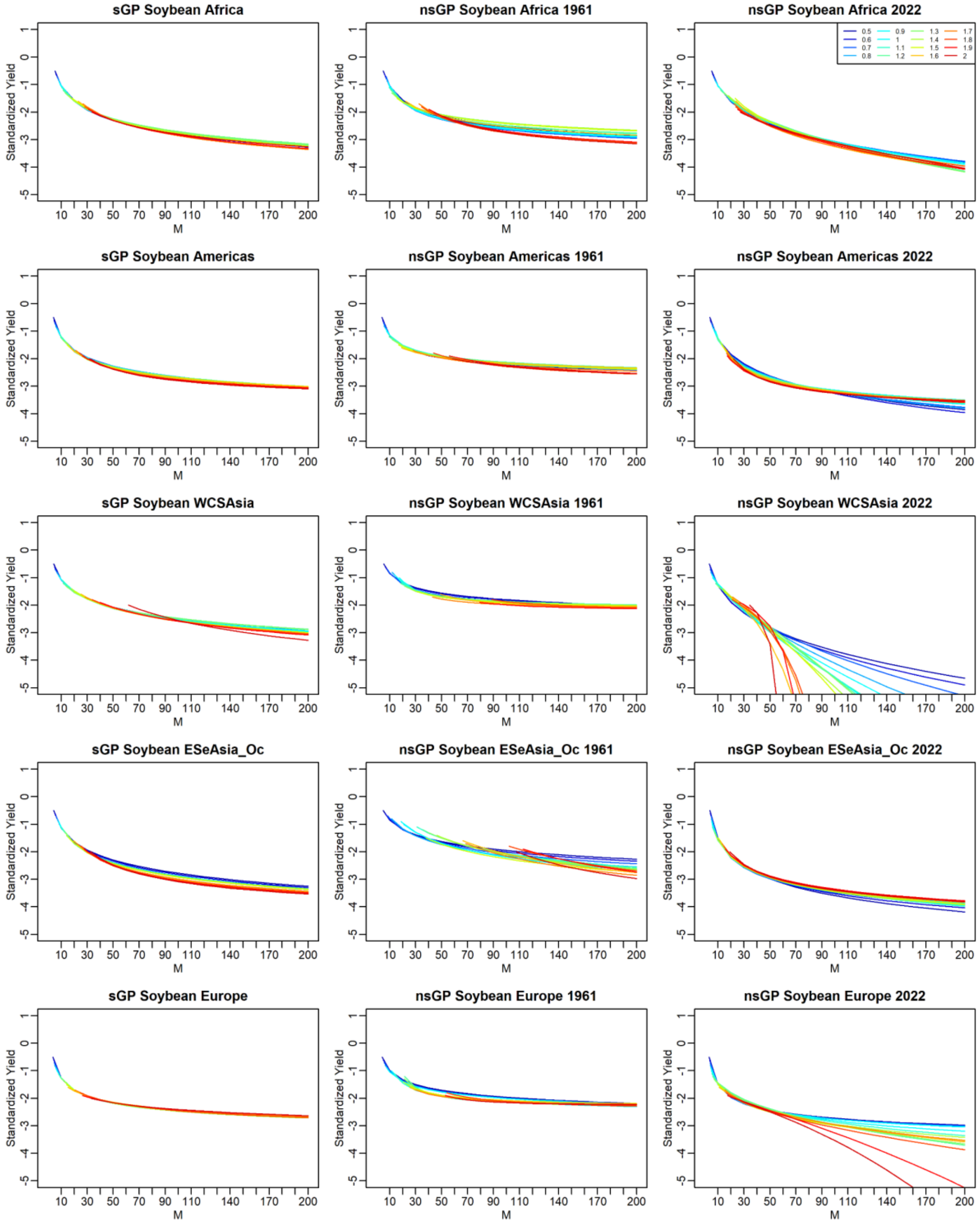


**Figure S16** *M*-year return levels estimated for soybean in the global yield data. The left panels are the levels estimated with the stationary generalized Pareto (sGP) models and the middle and right panels were those estimated with the non-stationary GP (nsGP) models. For the nsGP models, the levels estimated at the oldest (1961) and latest (2022) years are presented. Different colors indicate the different threshold ($T_h$) values.

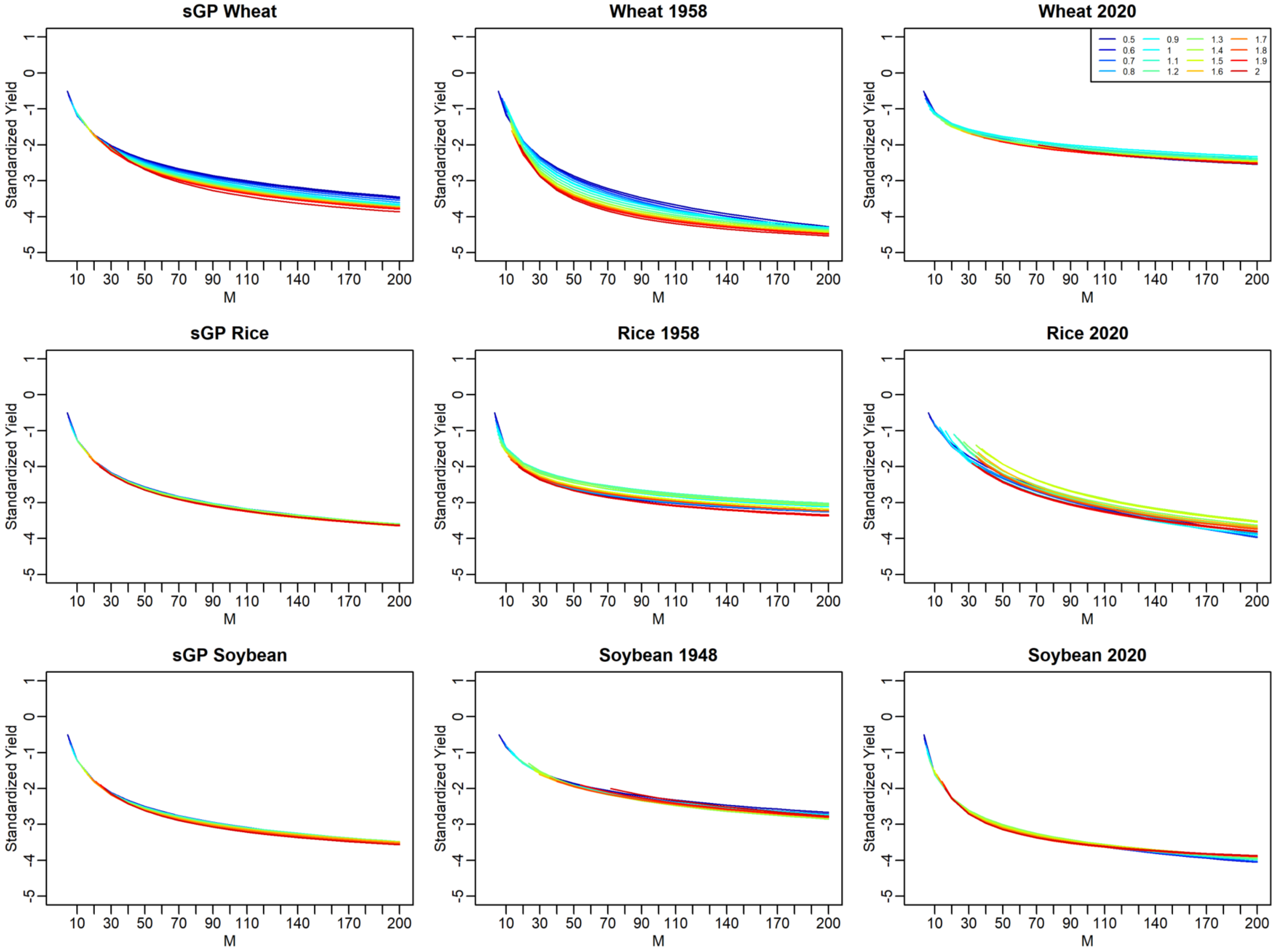


**Figure S17** *M*-year return levels estimated for the local yield data in Japan. The left panels are the levels estimated with the stationary generalized Pareto (sGP) models and the middle and right panels were those estimated with the non-stationary GP (nsGP) models. For the nsGP models, the levels estimated at the oldest (1958 or 1948) and latest (2020) years are presented. Different colors indicate the different threshold ($T_h$) values.

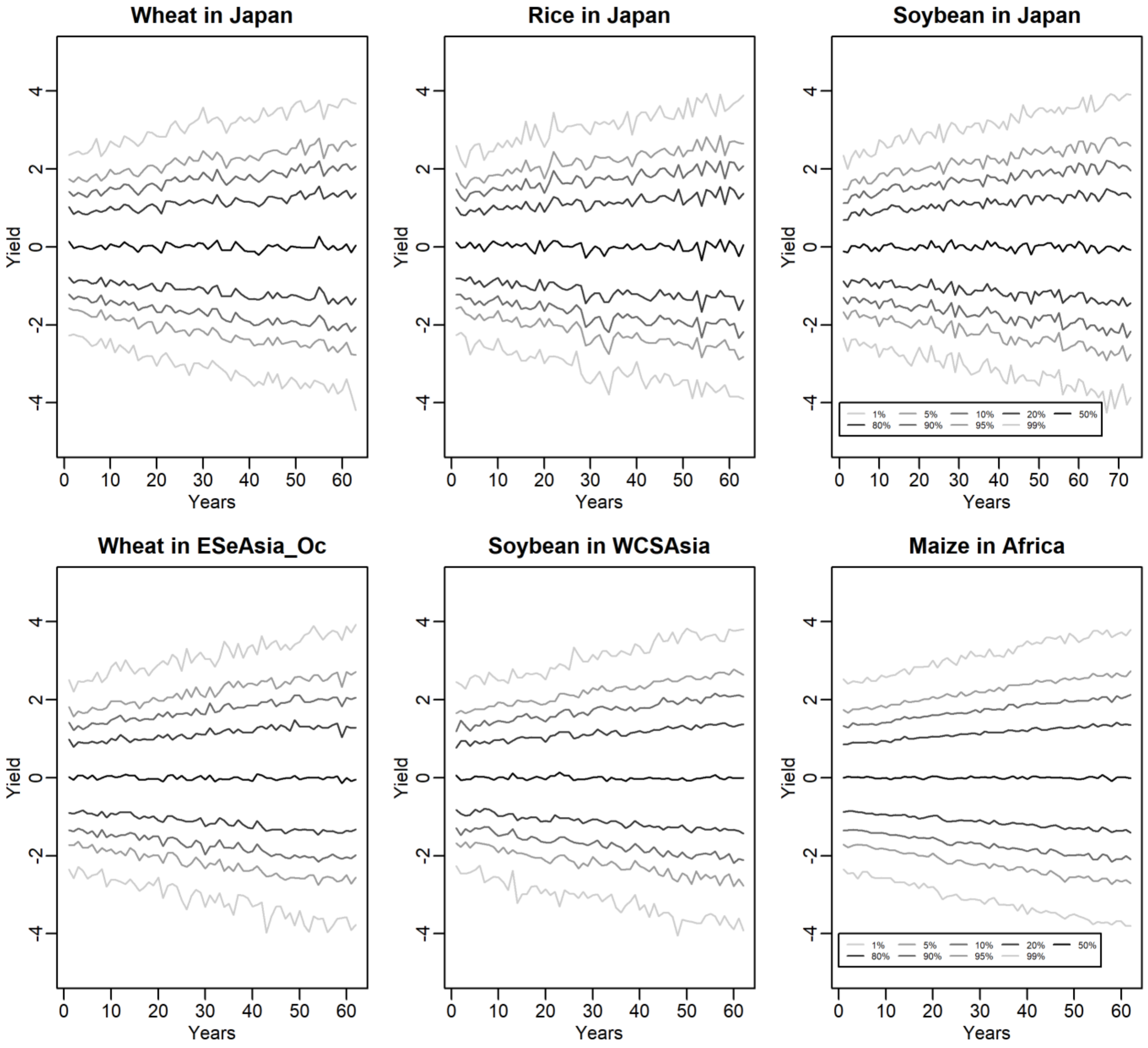


**Figure S18** Quantiles of simulated yield data. The y axis is the simulated yield of which scale corresponds with the transformed (i.e., detrended, scaled, and sign-inverted) scale of the real data analyses. The x axis indicate years. The quantiles were calculated by pooling 100 replications for each scenario. For each scenario, the variability of yield was designed to increase gradually. WCSAsia: Western, Central, and Southern Asia; ESeAsia_Oc: Eastern and Southeastern Asia and Oceania.

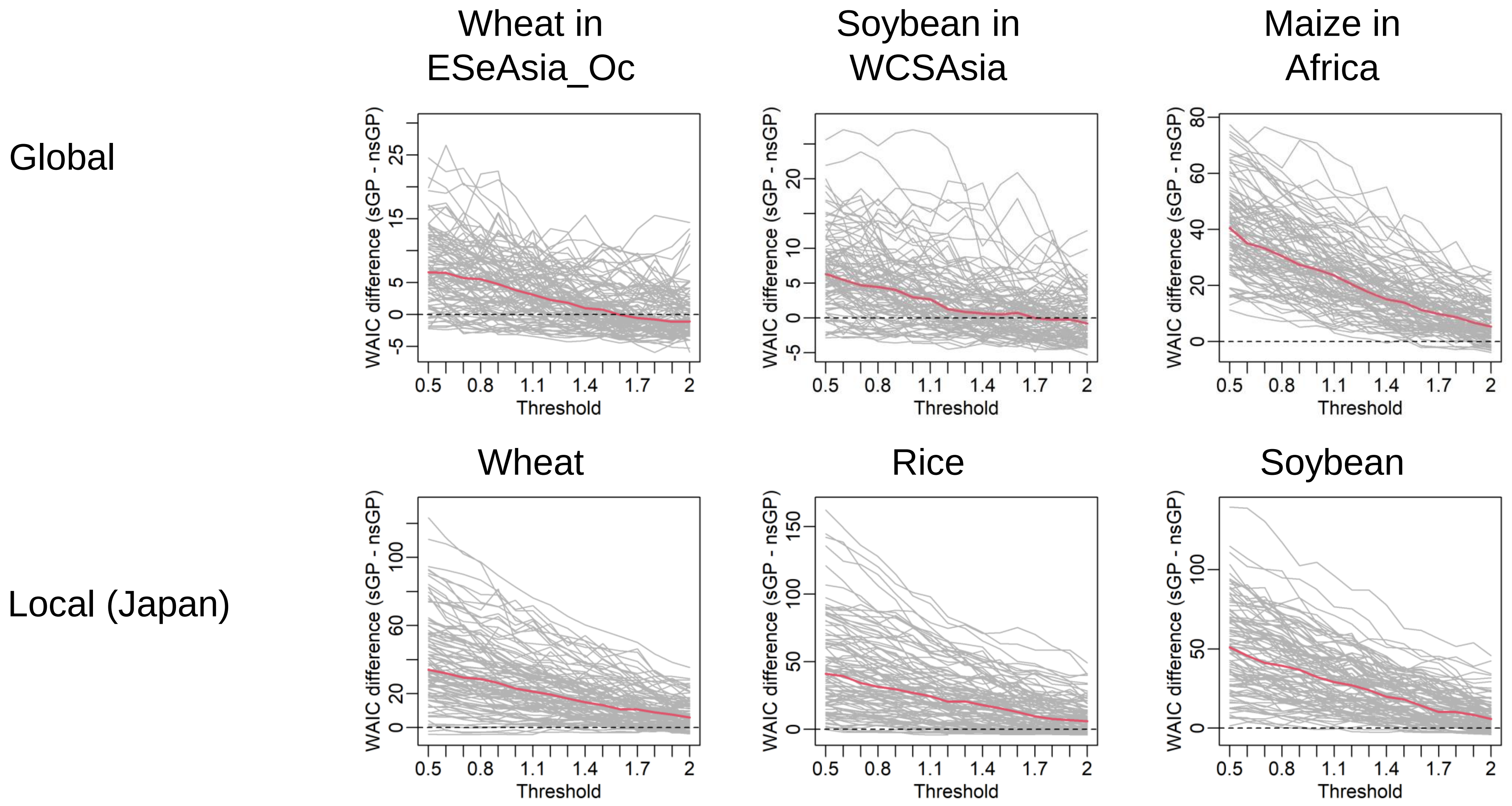


**Figure S19** Differences of WAIC values between the stationary generalized Pareto (sGP) and the non-stationary generalized Pareto (nsGP) models in five simulation scenarios. Positive values indicate that the nsGP models are supported. The gray lines are the trajectories of each replication, and the red lines are the averages.

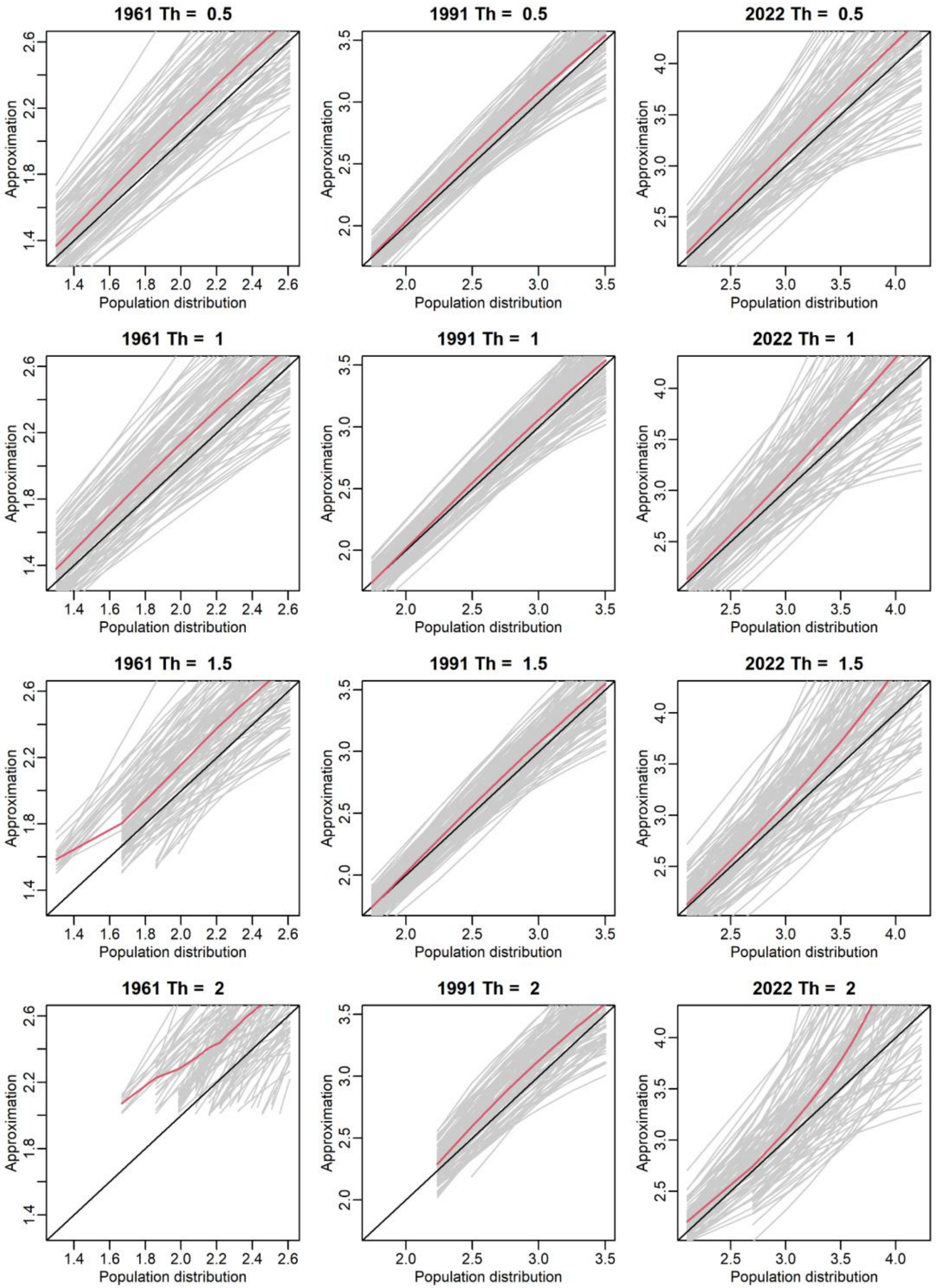


**Figure S20** Comparison of the *M*-year return levels in the simulations mimicking the wheat yield in ESeAsia_Oc. The y and x axes are the population (true) and approximated (estimated) return levels, respectively. The left, middle, and the right panels are for the oldest (1961), middle (1991), and latest (2022) years, respectively. The gray lines are the trajectories of each replication, and the red lines are the averages. The results obtained for four threshold values (0.5, 1, 1.5, and 2) are presented.

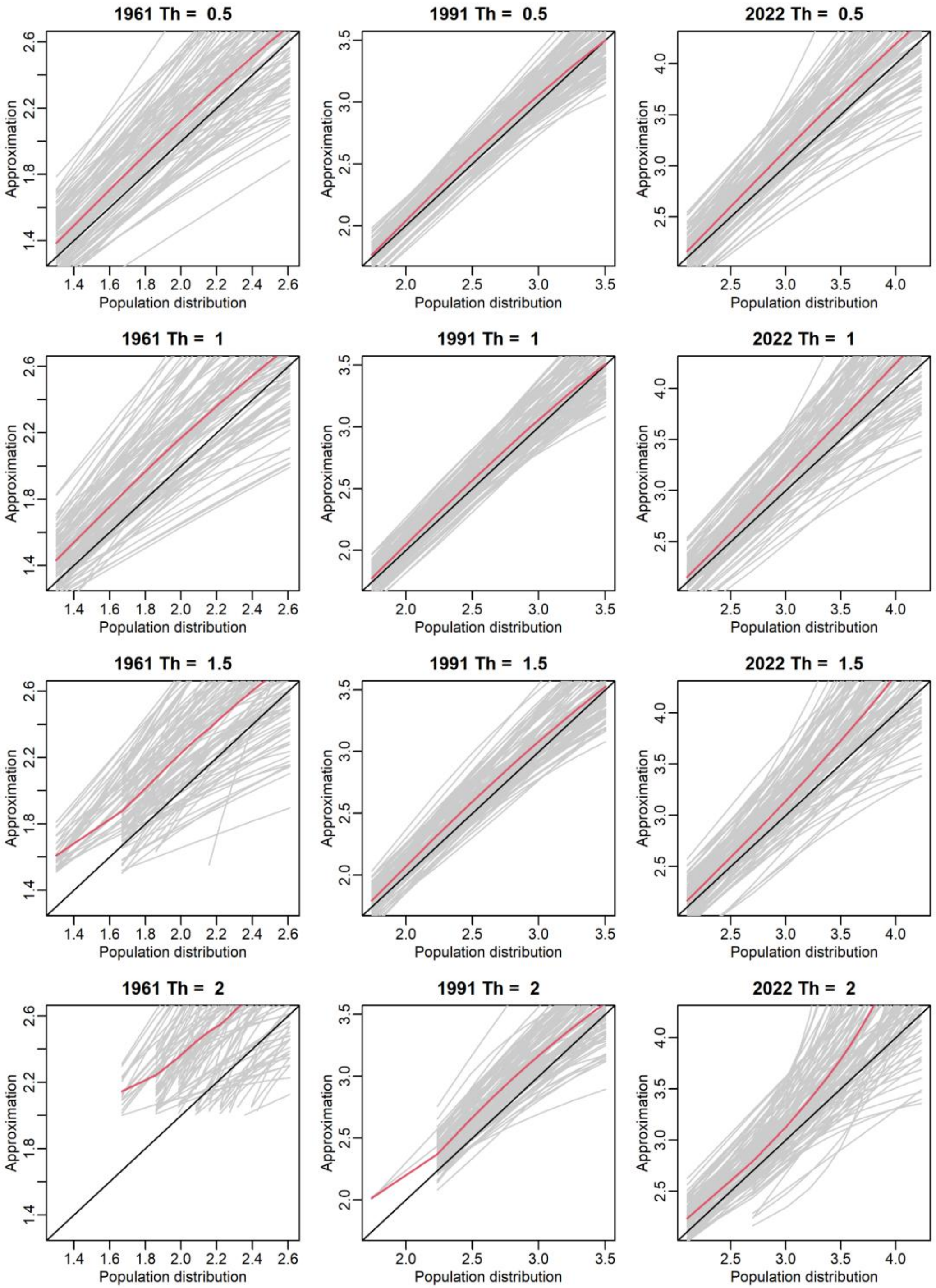


**Figure S21** Comparison of the *M*-year return levels in the simulations mimicking the soybean yield in WCSAsia. The y and x axes are the population (true) and approximated (estimated) return levels, respectively. The left, middle, and the right panels are for the oldest (1961), middle (1991), and latest (2022) years, respectively. The gray lines are the trajectories of each replication, and the red lines are the averages. The results obtained for four threshold values (0.5, 1, 1.5, and 2) are presented.

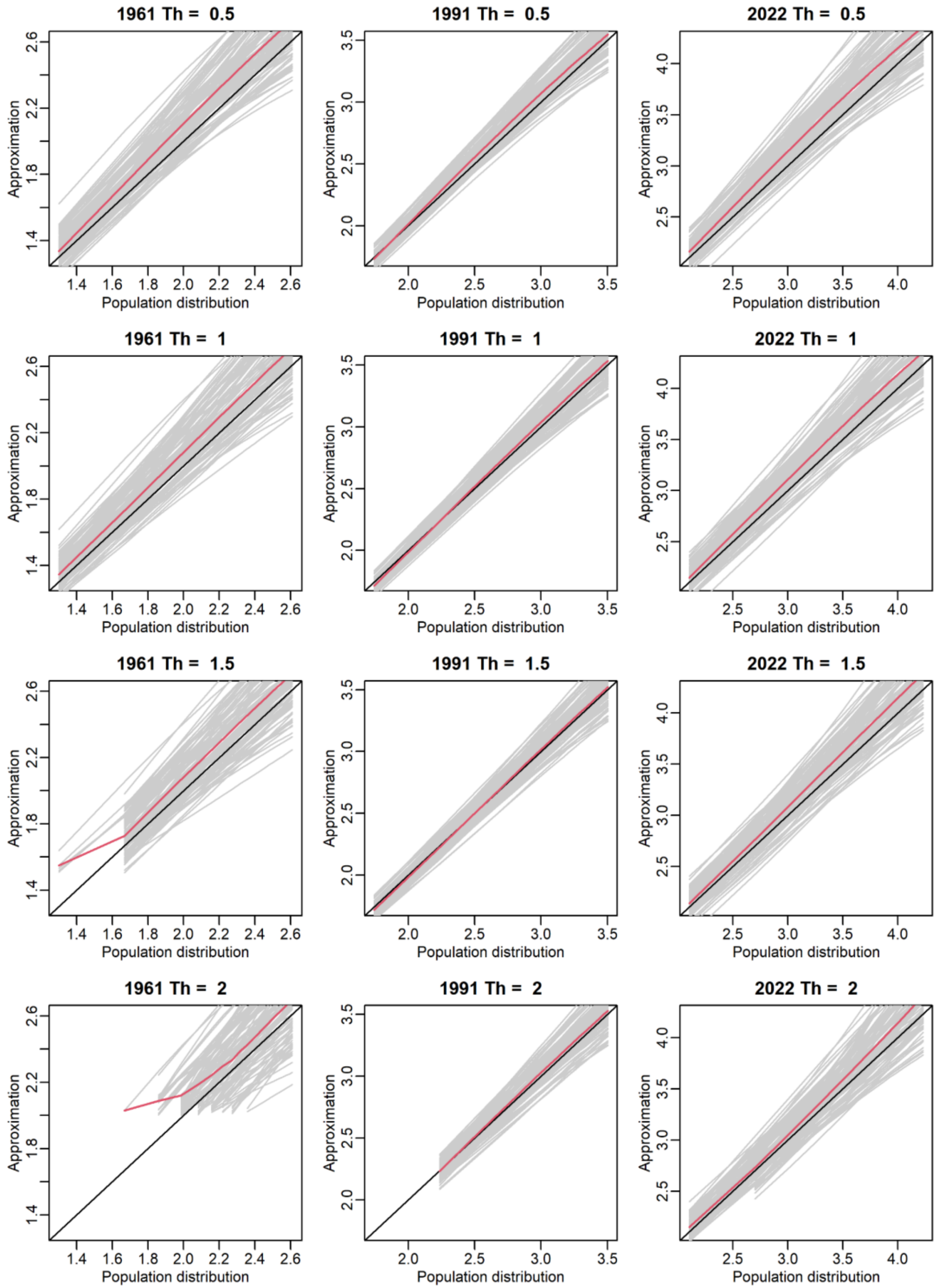


**Figure S22** Comparison of the *M*-year return levels in the simulations mimicking the maize yield in Africa. The y and x axes are the population (true) and approximated (estimated) return levels, respectively. The left, middle, and the right panels are for the oldest (1961), middle (1991), and latest (2022) years, respectively. The gray lines are the trajectories of each replication, and the red lines are the averages. The results obtained for four threshold values (0.5, 1, 1.5, and 2) are presented.

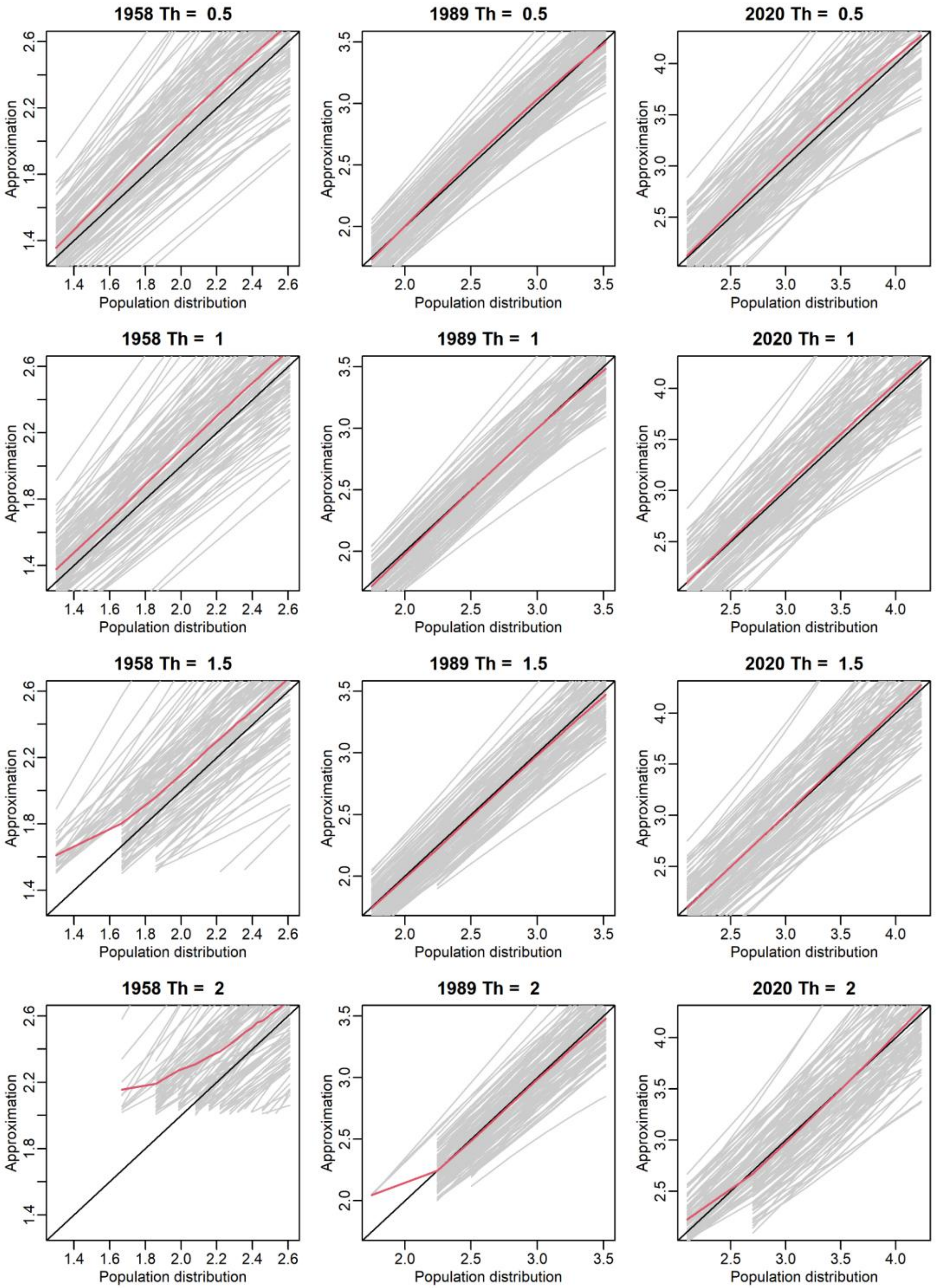


**Figure S23** Comparison of the *M*-year return levels in the simulations mimicking the wheat yield in the local yield data (Japan). The y and x axes are the population (true) and approximated (estimated) return levels, respectively. The left, middle, and the right panels are for the oldest (1958), middle (1989), and latest (2020) years, respectively. The gray lines are the trajectories of each replication, and the red lines are the averages. The results obtained for four threshold values (0.5, 1, 1.5, and 2) are presented.

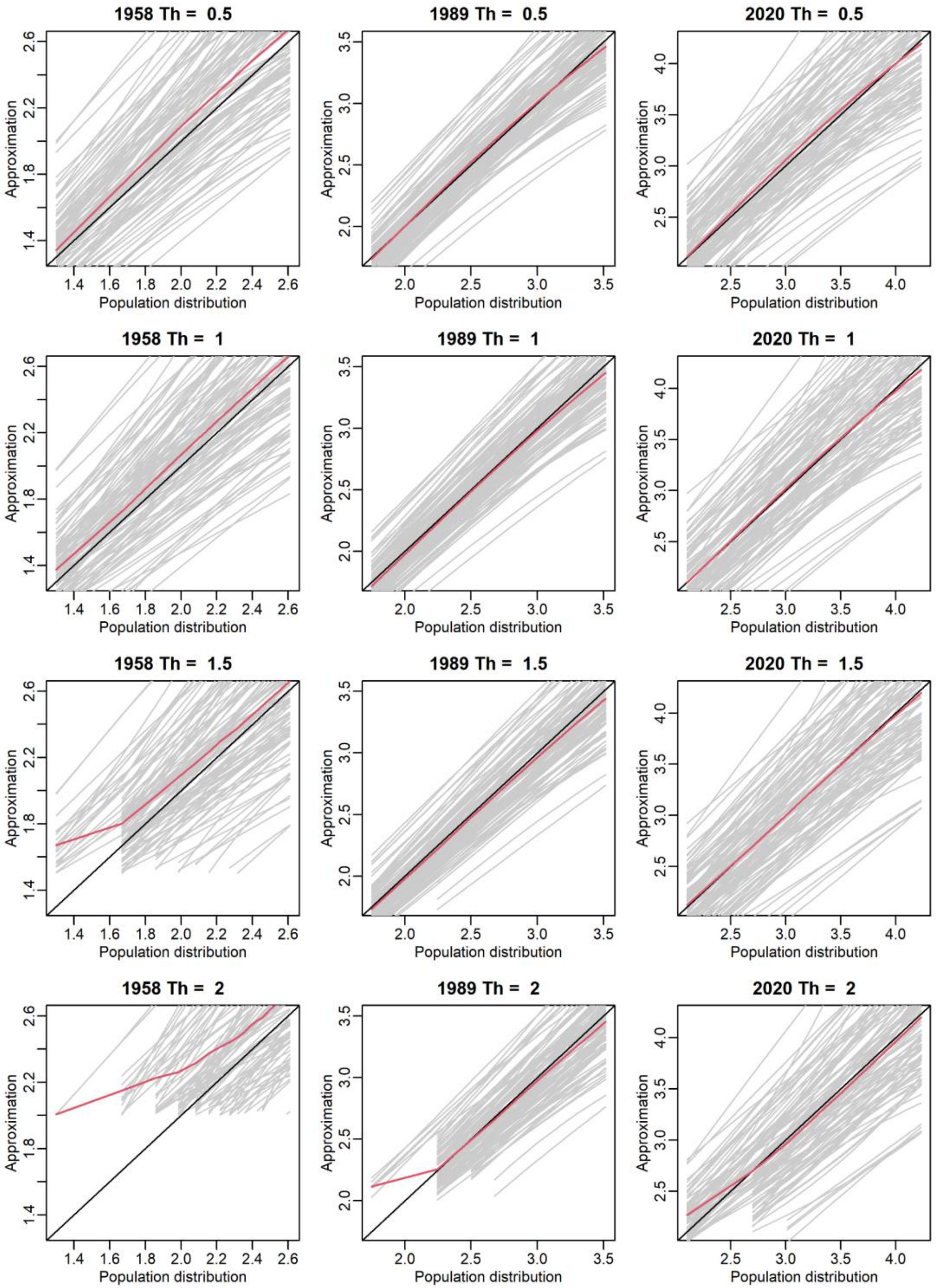


**Figure S24** Comparison of the *M*-year return levels in the simulations mimicking the rice yield in the local yield data (Japan). The y and x axes are the population (true) and approximated (estimated) return levels, respectively. The left, middle, and the right panels are for the oldest (1958), middle (1989), and latest (2020) years, respectively. The gray lines are the trajectories of each replication, and the red lines are the averages. The results obtained for four threshold values (0.5, 1, 1.5, and 2) are presented.

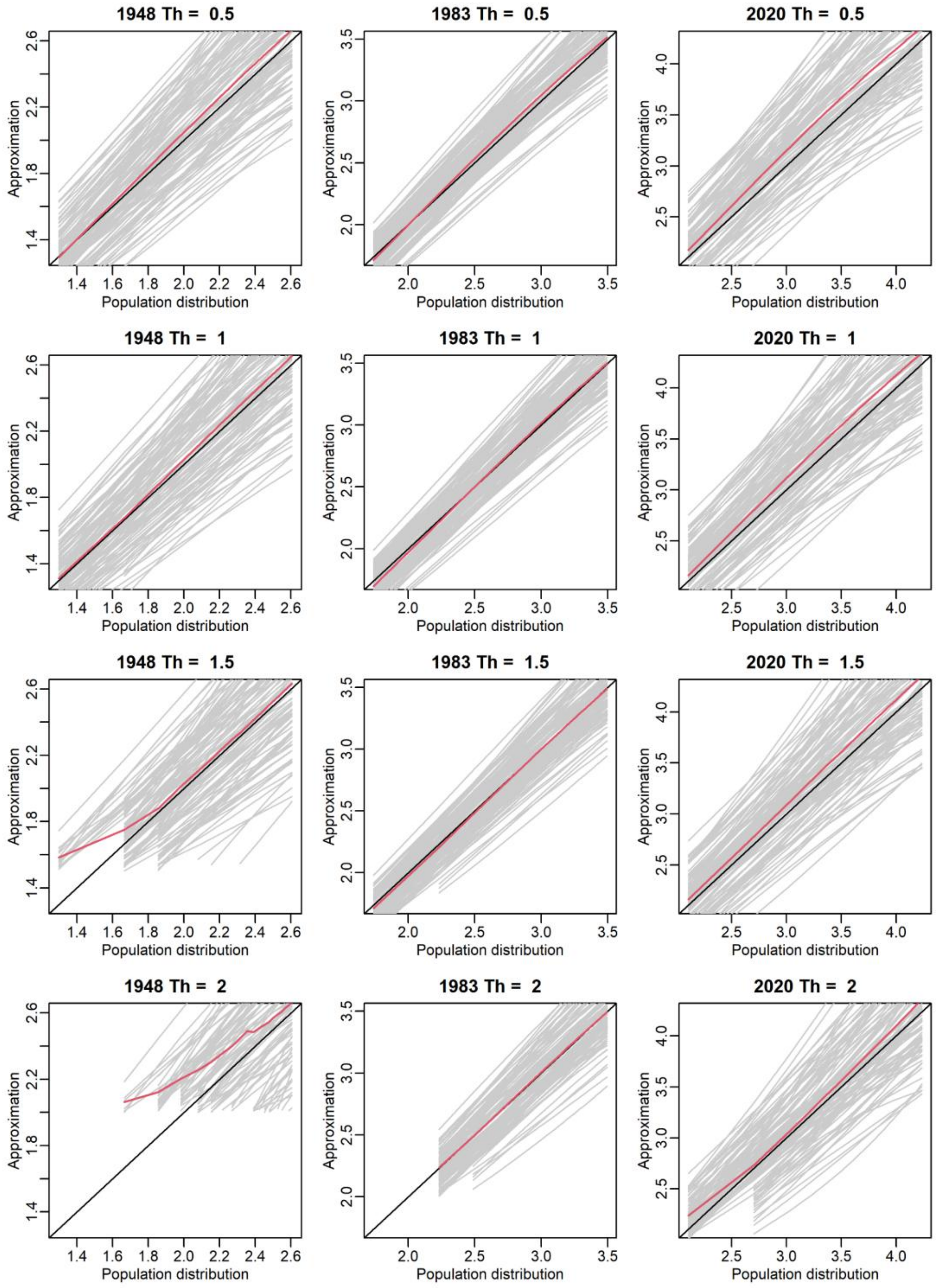


**Figure S25** Comparison of the *M*-year return levels in the simulations mimicking the soybean yield in the local yield data (Japan). The y and x axes are the population (true) and approximated (estimated) return levels, respectively. The left, middle, and the right panels are for the oldest (1948), middle (1983), and latest (2020) years, respectively. The gray lines are the trajectories of each replication, and the red lines are the averages. The results obtained for four threshold values (0.5, 1, 1.5, and 2) are presented.
